# High-throughput serial two-photon tomography with BakingTray and BrainSaw

**DOI:** 10.64898/2026.09.28.748989

**Authors:** Robert AA Campbell, Kurt I Anderson, Asli Ayaz, Emmanuel Beaurepaire, Hugo Blanc, Antonin Blot, Juan Carlos Boffi, Vincent Bonin, Howard HW Chan, Beverley Clark, Georgios Foustoukos, Peter Gordon, Del Halpin, Yunyun Han, Michael Häusser, Sonja B Hofer, Friedrich W Johenning, Stephan Junek, Loukia Katsouri, Michael Krumin, Pierre Mahou, Troy W Margrie, Adrien Philippon, Viktor M Plattner, Claudio Polisseni, Solène Prudhomme, Rajeev Rajendran, Simon Townsend, Nicole Vissers, Florian Vollrath, Simon Weiler

## Abstract

Volumetric imaging is increasingly important in fields such as neuroscience, where quantifying structure and mapping features are inherently three-dimensional problems. Block-face serial two-photon tomography (STPT) has emerged as a powerful approach for high-resolution, whole-organ volumetric imaging. STPT avoids complex sample preparation and balances image quality, speed, and scalability. However, STPT remains niche, in part because of the absence of flexible, general-purpose control software. Here we present BakingTray, an open-source software platform for STPT that extends ScanImage. BakingTray’s modular framework accommodates diverse hardware configurations and allows users to build systems customised to their needs. Twelve such systems, which we call ‘BrainSaws’, have been constructed worldwide since 2016, with more under construction, and have so far contributed to 35 research papers. The three systems at the Sainsbury Wellcome Centre alone have acquired over 7,000 samples in a busy multi-user facility. We demonstrate the platform across its core neuroscience applications: electrode track localisation, brain-wide cell counting, and bulk projection mapping. Beyond these, we map the cell-type-specific synaptic output of inhibitory neurons in the ventral lateral geniculate nucleus and, with whole-mount staining, amyloid plaques brain-wide in a mouse model of Alzheimer’s disease. BakingTray is a mature platform for groups or facilities needing a customised volumetric imaging solution, whether built new or converted cheaply from an existing two-photon microscope.

## 1 Introduction

### 1.1 Volumetric imaging

Modern anatomy increasingly relies on registering samples to a three-dimensional reference atlas, so that structures can be delineated automatically and results compared across animals in a common coordinate space (Osten and Margrie 2013; Tyson and Margrie 2022). Registration is most straightforward when the sample has been imaged as an intact volume, since the geometric relationship between planes is preserved. Cutting a sample into sections and mounting them on slides breaks that relationship, as sections are independently distorted and may even be damaged. Aligning such datasets to a 3D atlas is not straightforward (Puchades *et al*. 2019; Puchades *et al*. 2025). Volumetric imaging sidesteps the problem, but remains technically challenging for larger structures such as mouse brains, with no single solution optimal across all sample types.

Rendering tissue transparent with optical clearing (Richardson and Lichtman 2015) and subsequent imaging on a lightsheet microscope (Stelzer *et al*. 2021) is a popular method for volumetric imaging. This process avoids the need for slow manual sectioning and is compatible with antibody labelling (e.g. Renier *et al*. 2014). Clearing can optionally be coupled with tissue expansion (Chen, Tillberg, and Boyden 2015), which is especially powerful when combined with microscopes tailored for the technique (Glaser *et al*. 2025). However, clearing can be time-consuming and requires significant expertise for optimal results. Serial-sectioning techniques offer an alternative, particularly for samples that do not require whole-mount labelling.

### 1.2 Serial-section microscopy

Serial-section imaging techniques bypass the need for clearing because the sample is sliced or milled and only the exposed face is imaged. Slices are typically discarded and, since the block is near-rigid, the images can simply be ‘stacked up’ in order to produce a 3D volumetric image. Serial-section techniques are easy to scale, as multiple samples can be arranged in a grid and processed simultaneously with little increase in human labour time. Early systems (Figure 1–figure supplement 1) used episcopic (incident-light) imaging to acquire the block face of samples prepared by resin embedding (Odgaard *et al*. 1990; Ewald *et al*. 2002), paraffin embedding (Weninger *et al*. 1998; Weninger and Mohun 2002; Weninger *et al*. 2006), or cryo-sectioning (Spitzer *et al*. 1996; Zhang *et al*. 2003; Roy *et al*. 2009). The same approach works with electron microscopy (Leighton 1981; Soto *et al*. 1994; Denk and Horstmann 2004; Kasthuri *et al*. 2015). In knife-edge scanning microscopy the sample is imaged as it is cut, whilst the section lies over the blade (Mayerich, Abbott, and McCormick 2008; Pesavento *et al*. 2017). Thus, the physical section thickness sets the *effective* optical section thickness: slices as thin as 1 *µ*m can be imaged with conventional brightfield (Li, Gong, *et al*. 2010) or fluorescence (Gong *et al*. 2013) microscopy, with no need for elaborate optical sectioning techniques. Cutting reliably at these thicknesses requires hard blocks generated with paraffin or resin embedding. However, such embedding is time-consuming and may, depending on the protocol, quench fluorescent proteins. This creates a need for alternative serial-sectioning approaches based on true optical sectioning with minimal tissue processing.

### 1.3 Block-face imaging with optical sectioning

Agarose-embedded fixed tissue is easy to prepare and often cuts well with a vibratome, but the resulting sections are relatively thick and highly deformable, and so are poorly suited to knife-edge imaging. High-resolution imaging of agarose-embedded samples therefore requires a microscope with intrinsic optical sectioning capability. This can be achieved in a variety of ways, such as serial-section lightsheet (Narasimhan *et al*. 2017; Yang *et al*. 2018) and systems based on UV excitation epifluorescence microscopy (Kolluru *et al*. 2022; Yu *et al*. 2022). Confocal approaches have also been used, with systems built using both spinning disk (Seiriki *et al*. 2017; Seiriki *et al*. 2019) and slit confocal (Deng *et al*. 2022, where a cryostat was used). Whilst these confocal techniques excel at optical sectioning, for block-face imaging this comes at the cost of poor depth penetration and potentially low contrast due to scattering and pinhole cross-talk.

Two-photon microscopy allows imaging deeper into scattering tissue than confocal (Denk, Strickler, and Webb 1990), and in a block-face system it need only reach 50–100 *µ*m below the surface (a little more than one section thickness); even dense organs or large specimens therefore remain accessible. Two-photon microscopy has consequently been used in a variety of serial-section block-face imaging systems. The earliest such systems removed imaged tissue by laser ablation (Tsai *et al*. 2003) or milling (Ragan *et al*. 2007). However, these processes are more destructive than sectioning and are difficult to work with. Contemporary serial two-photon tomography (STPT, Fig. 1) microscopes use a vibratome, and have been built using linear scanners (Ragan *et al*. 2012), acousto-optical deflectors (Zheng *et al*. 2013), resonant scanners (Economo *et al*. 2016), and multi-focal scanning (Chen *et al*. 2020). Scanner-based approaches, such as STPT, acquire multiple channels simultaneously without the alignment concerns inherent in multi-camera approaches, such as spinning-disk confocal. Conventional STPT systems can readily handle three simultaneous fluorophores; specialist systems generate trichromatic two-photon excitation by combining pulses from two lasers, exciting a broader fluorophore palette (Abdeladim *et al*. 2019). STPT has been applied to a number of fields, including cancer biology (Grauel *et al*. 2020; Taranda *et al*. 2021) and immunology (Li *et al*. 2021; Li *et al*. 2024). It has in particular revolutionised neuroscience, where it was used to generate the Allen mouse brain atlas (Wang *et al*. 2020) and, along with related techniques (Gong *et al*. 2013; Gong *et al*. 2016; Gao *et al*. 2022), pioneered brain-wide single-axon tracing in mouse (Han *et al*. 2018; Winnubst *et al*. 2019).

**Figure 1.**
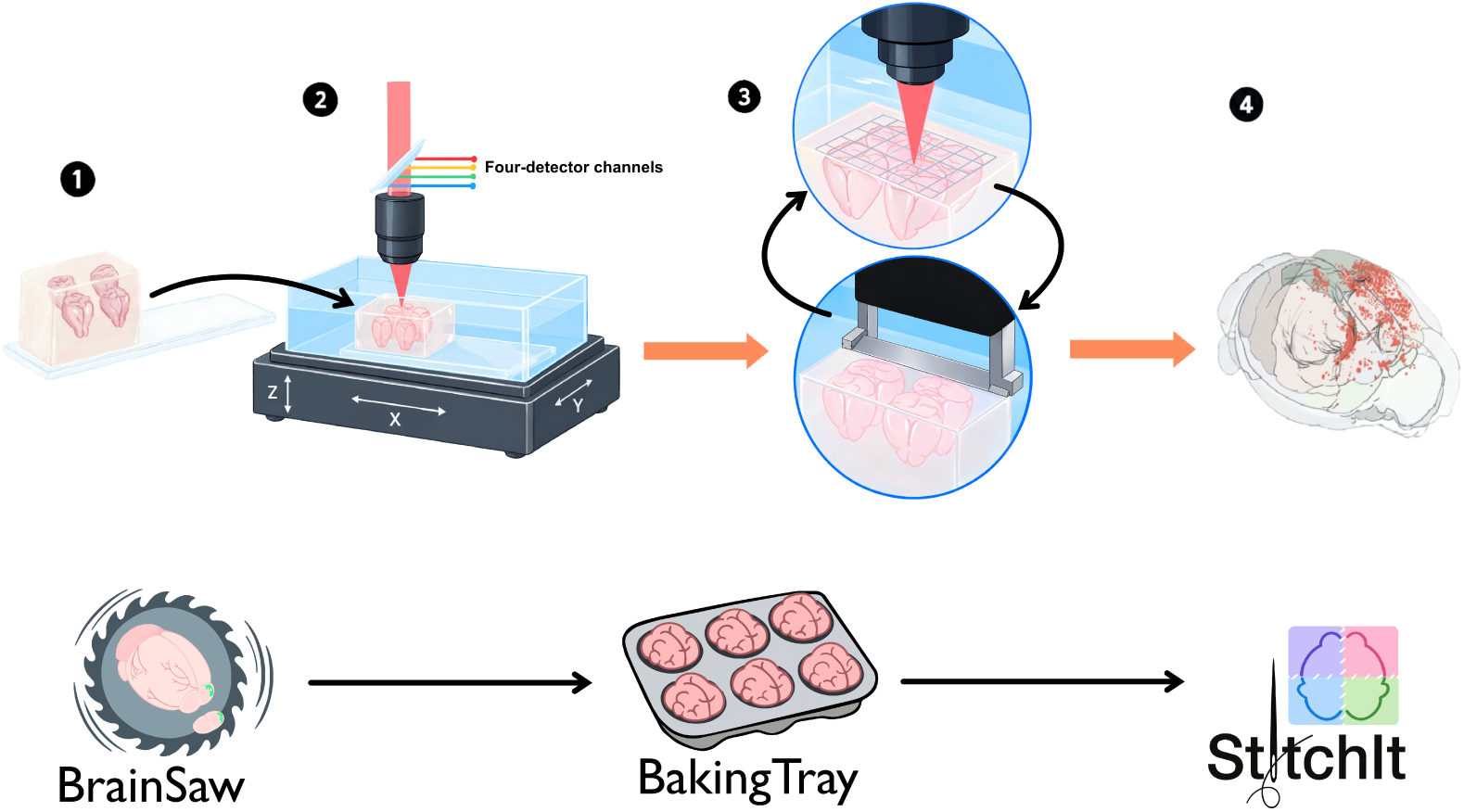
Procedure for block-face serial-section imaging. STPT systems slice and image fixed, agarose-embedded samples using a two-photon microscope coupled to a vibratome. The blade and objective are fixed, although the objective can take a small z-stack with a piezo focuser during imaging. **Step 1**. In this case four brains are embedded in agarose and the agarose block is glued to a microscope slide. **Step 2**. The slide is clamped into a water bath filled with phosphate buffer; the water bath rides on a 3-axis stage. The stage provides for tile-scanning as well as driving the sample into the vibratome for slicing. The sample is imaged with a two-photon microscope. **Step 3**. The system tile-scans over the exposed surface of the sample at the block face. Once images have been acquired, the sample is sliced. The cutting cycle raises the sample by the thickness of one physical section, drives it into the vibrating blade at low speed, then returns it to the imaging position. The cut sections sink to the bottom of the chamber and are discarded. A new tile scan is then performed across the exposed block face and the cycle repeats until the whole organ has been imaged. **Step 4**. At the end of the acquisition, data are stitched and assembled into a 3D volumetric image. The resulting data can be analysed using automated approaches, with results displayed in a common coordinate space. See also: a historical timeline of serial-sectioning approaches is presented in Fig. 1–figure supplement 1.

### 1.4 BakingTray

Despite its impact, STPT remains confined to relatively few laboratories. Commercial instruments are costly and may not offer the required features, which has driven some groups to build their own systems. Researchers wishing to self-build can draw on published hardware designs, including plans from the Teravoxel project (Economo *et al*. 2016) and the excellent hardware description in the Seiriki *et al*. 2019 spinning-disk methods paper. Control software, however, has lagged: the Teravoxel project makes available fetch, a C++ software package for running STPT, but it was written around that instrument’s specific digitiser and motion-control hardware and has not been updated since 2015. Its MouseLight fork was last updated in 2020 but remains hardware-specific. Here we present BakingTray, which is, to our knowledge, the only hardware-agnostic, actively maintained control platform for STPT. BakingTray extends ScanImage (Pologruto, Sabatini, and Svoboda 2003), the popular two-photon acquisition software, via its API, adding tile-scanning, automatic sample detection, slicing, and automation of sample setup tasks. Furthermore, it allows users to build a system tailored to their needs and budget, rather than adopting a fixed design. We refer to any STPT system running BakingTray as a ‘BrainSaw’.

We show that BakingTray is an established and reliable platform for 3D imaging of a variety of samples. In particular, it excels in three key neuroscience applications: electrode track localisation, brain-wide cell counting, and bulk projection mapping. We also demonstrate dual-colour brain-wide cell counting, explore projections and synaptic sites of multiple inhibitory cell types of the ventral lateral geniculate nucleus (vLGN), and show brain-wide staining of amyloid plaques in a mouse model of Alzheimer’s disease. BakingTray brings automated, high-throughput volumetric imaging of whole organs within reach of research groups and core facilities alike, and is particularly cost-effective where an existing two-photon microscope can be repurposed.

## 2 Results

### 2.1 Design overview

BakingTray is a general-purpose STPT software solution based around ScanImage. ScanImage handles image acquisition, control of laser power and PMTs; BakingTray handles setting of imaging parameters, tile scanning, slicing, and automatic sample detection. BakingTray supports acquisition of up to four detector channels and multiple laser lines via ScanImage.

BakingTray is not tied to a specific hardware configuration. The software can be extended to incorporate new hardware components (stages, lasers, vibratome controllers, etc.) and the code is extensively documented to make this straightforward. The core acquisition class contains no hardware-specific commands: each category of hardware is instead represented by an abstract class that declares a small, documented interface. Support for a new device requires only a single class that implements it (Supplementary Table 1). To aid development, a simulated mode allows BakingTray to be run without any hardware present (e.g. see video on the documentation pages). Currently supported hardware is listed in Supplementary Table 2. The modular nature of BakingTray means that it is possible to convert an existing resonant scanning two-photon microscope to an STPT system for around 30,000 EUR (see Appendix 1), or to upgrade legacy TissueCyte 1000 systems (Appendix 2).

### 2.2 BakingTray is production-ready and stable

Twelve BrainSaws have been constructed worldwide (Supplementary Table 3) and have contributed to 35 publications (Supplementary Table 4). We estimate that about 40 research groups have published with BakingTray or are actively working with it.

To quantify system throughput and reliability we analysed historical acquisition log files for the three BrainSaw systems at the Sainsbury Wellcome Centre (SWC) by scraping data from our Centre’s storage platform (Methods Section 4.3). The Centre initially had one microscope; this was joined by a second in 2019 and a third in 2021. We found 7,090 samples imaged across 3,251 unique acquisitions; roughly half of these acquisitions (1,676) contained two or more samples and 678 contained four or more (Fig. 2A&B), reflecting the routine use of multi-sample runs to raise throughput. Acquisitions at around 2 and 4 *µ*m/pixel are the most common (Fig. 2C), as these are the resolutions we recommend for our main use cases (see Results Section 2.6).

**Figure 2.**
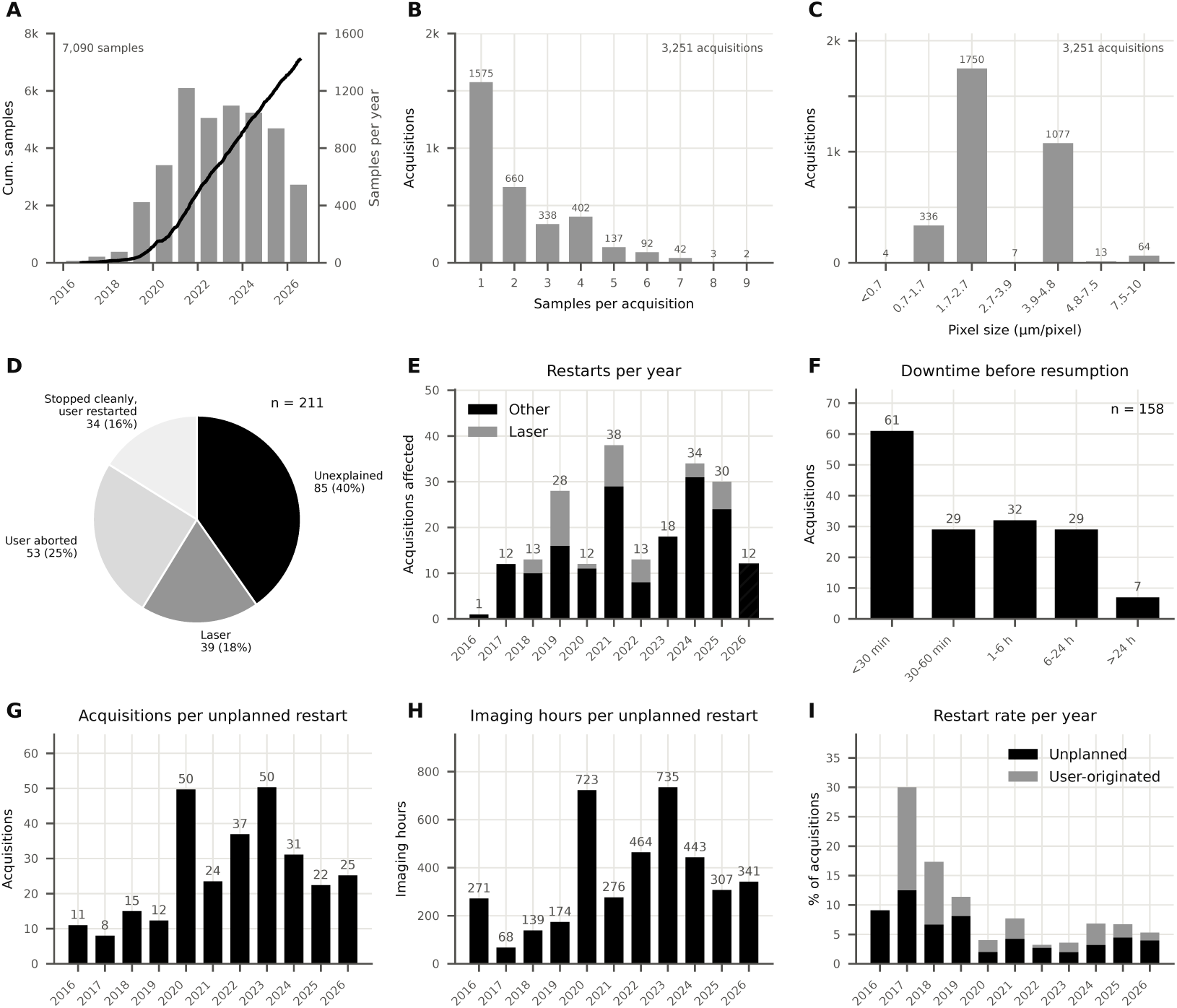
Acquisition throughput and reliability of the three microscopes at the SWC. Data before 2018 were acquired by one of the microscopes now at the SWC before it was moved there. Graphs show data up to mid-July 2026. **A**. Line shows the cumulative number of imaged samples as a function of time (left axis). Grey bars show the total number of samples imaged each year (right axis). **B**. BakingTray supports multiple samples per acquisition. Panel shows the distribution of the number of samples imaged in a single acquisition, e.g. in 139 acquisitions six or more samples were imaged. **C**. Lateral (X/Y) pixel size at which these acquisitions were conducted. Around 2 *µ*m/pixel is suggested for cell counting, and 4–8 *µ*m/pixel suggested for mapping electrode tracks and fibre tracks (Supplementary Table 5). **D**. In 211 cases acquisitions were restarted for some reason (restarts after five or fewer sections, which reflect setup errors, are not counted; see Methods). On 53 occasions the user manually aborted then restarted the acquisition. On 34 occasions the acquisition stopped as planned but was then restarted. On 39 occasions BakingTray stopped acquiring because the laser was reported as not running. In the remaining 85 cases the acquisition stopped for an unknown reason and was then restarted. **E**. Breaks down the number of restarts by year. Restarts due to the laser are highlighted; these were more likely due to a bug in the serial communications protocol (fixed in 2026) than to a true laser problem. **F**. The time between an acquisition stopping and its restart. These data do not include ‘user aborted’ restarts, as typically those were restarted immediately. **G**. Shows a time to failure measure: the number of acquisitions per unplanned restart in each year. **H**. Re-casts the data in G in units of acquisition hours per unplanned restart. **I**. Restart rate per year. User-originated restarts followed a stop that the user requested or that BakingTray made by design (a manual abort, reaching the requested number of sections, or finding no tissue). Unplanned restarts followed a stop forced by an error, such as a laser fault, a hard crash of MATLAB or a power cut. The years 2016–2019 had fewer acquisitions, so their rates are less reliable, and BakingTray was under heavier development at that time.

Setting up a sample is fast but the acquisition itself runs for several hours, often well into the night. It is therefore important that BakingTray is stable and does not require excessive intervention for acquisitions to complete successfully. We assessed reliability by further mining the log data on our storage platform. The acquisition metadata files record not only resolution and the number of samples but also how long they took to acquire and whether acquisitions were manually aborted, restarted, or otherwise did not terminate as expected. Of these 3,251 unique acquisitions, 94.5% completed without requiring a restart; the remaining 180 interrupted acquisitions (5.5%) were restarted 211 times (Fig. 2D), for two kinds of reason: stops the user requested or BakingTray made by design (87; user-originated) and genuine failures (124; unplanned). Restarts that occurred within five sections of an acquisition start event were excluded from these numbers. Such restarts were highly likely to be associated with a user error and do not reflect system reliability.

The 87 user-originated restarts were of two kinds. In 53 cases the user *manually* aborted and restarted the acquisition, for example to change imaging settings or correct sample mounting issues. In 34 cases the acquisition stopped naturally without error but was subsequently restarted by the user. Reasons for this include the user asking for too few sections or BakingTray being unable to find the tissue and therefore stopping for safety reasons. A previously cut section floating between the objective and sample can cause a safety stop of this sort.

Acquisitions stopped prematurely on 124 occasions due to an error. There were 39 failures associated with the laser; these were more likely a loss of serial communications due to a now-resolved bug than a true laser failure (Fig. 2E). Finally, on 85 occasions acquisitions stopped abruptly with no logged error but were then restarted. These reflect events such as power cuts, random hard-crashes of MATLAB, very rare ScanImage frame logging errors, or Windows update reboots that proceeded despite the update policy being set to avoid unattended reboots. In most cases acquisitions were resumed in under an hour (Fig. 2F), partly because an acquisition that ends early is reported by a Slack message. The rate at which the systems generated errors that required an unplanned restart (i.e. due to the laser [n=24] or for unknown reasons [n=69]) has stayed fairly constant since 2020 (Fig. 2I), with about 1 in 34 acquisitions requiring intervention to complete (Fig. 2G; mean of yearly values, 2020–2026). This corresponds to one restart intervention for every 470 hours of acquisition time (Fig. 2H; again the mean of yearly values), roughly once per month of nightly use.

### 2.3 BakingTray user experience

To set up a sample, the user begins by defining the sample ID, imaging resolution, and section thickness in the main BakingTray window (Fig. 3A), with channels to acquire defined in ScanImage. BakingTray’s ‘Prepare’ GUI (Fig. 3B) is used to trim the block down to the starting depth, and slicing is checked for consistency (see Appendix 5). The ‘Acquisition’ GUI (Fig. 3C) allows the user to take a fast low-resolution tile scan, in order to assess image quality, set the imaging depth, and check for alignment of the vibratome blade (see Appendix 4). Once ready, the user performs an ‘Auto-Thresh’ (Fig. 3C) to automatically identify the tissue and fit it with a tile pattern (see Methods Section 4.2), then the acquisition is initiated. After each section has been imaged, the microscope slices the top of the block; the acquired section is removed and sinks to the bottom of the water bath. The tissue boundaries are re-detected automatically after each section, so the imaged area tracks the changing shape of the sample at the block face. The acquisition proceeds until the requested number of sections has been acquired or no further tissue is found; the laser is then automatically switched off and PMT gains set to zero. BakingTray communicates with the laser (Fig. 3D), setting the wavelength and stopping the acquisition automatically if the laser loses mode lock. Safeguards the software imposes (Appendix 3–table 1), along with other operating principles, are described in Appendix 3 and Methods Section 4.2.

**Figure 3.**
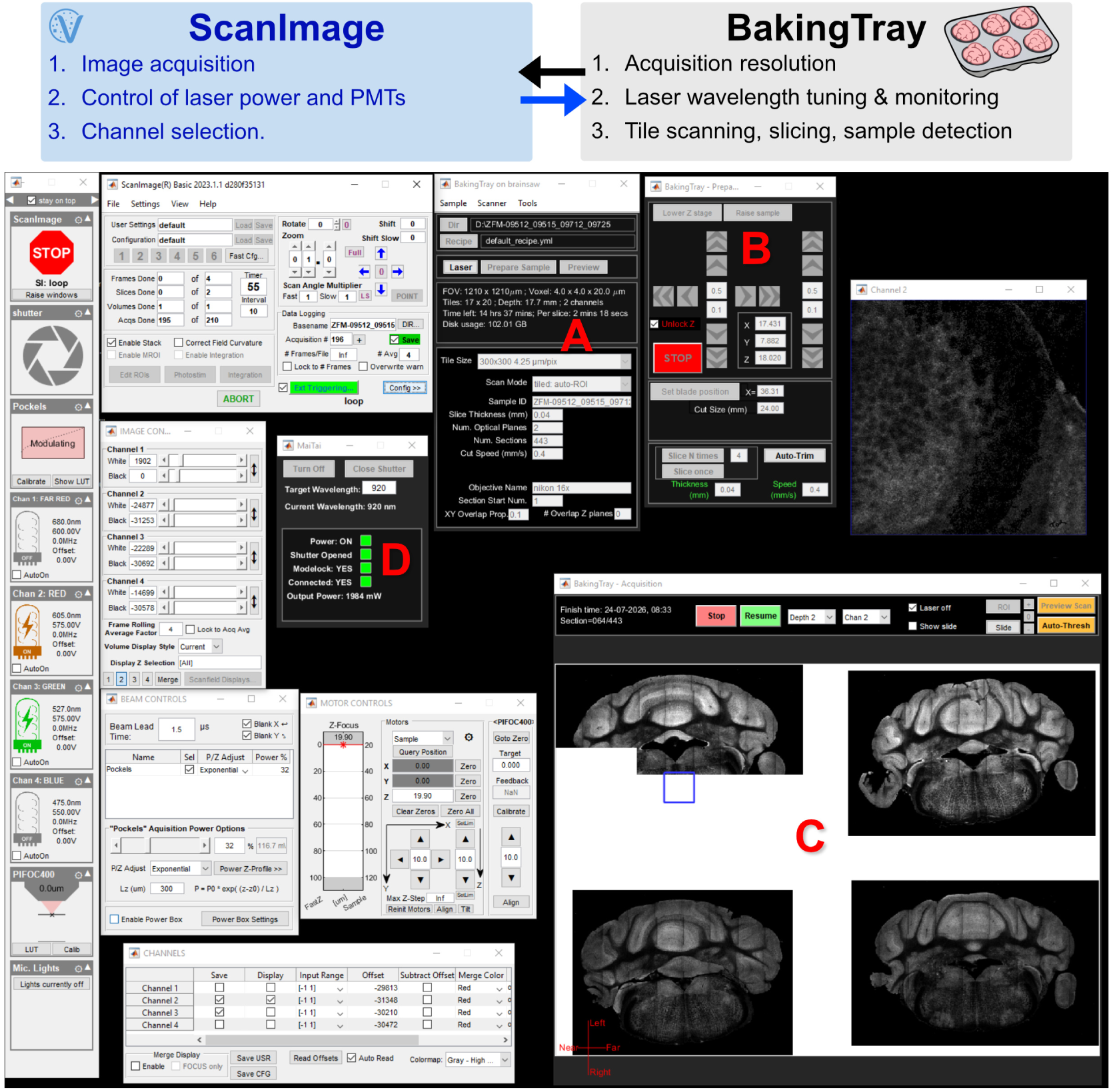
Acquisition screenshot showing ScanImage & BakingTray. Screenshot shows a typical view of the software during acquisition: grey ScanImage windows are on the left and the black BakingTray windows (labelled **A**–**D**) are on the right. ScanImage and BakingTray interact bidirectionally with each other to form a cohesive user experience. ScanImage is unmodified and handles image acquisition, control of laser power, control of PMTs, fast and coarse focus, and channel selection. **A**. Main BakingTray window where the user sets imaging parameters such as the sample save path, image resolution, number of slices, etc. **B**. The ‘Prepare’ GUI moves the XYZ stages interactively during sample set up and handles initial sample trimming. **C**. The ‘Acquisition’ GUI displays a live preview of the sample, which is used to set the initial imaging ROI and monitor progress locally during acquisition. **D**. The ‘Laser’ GUI interfaces with the laser, allowing the user to change wavelength, gate the shutter, and power the laser up/down.

Setup is the only hands-on stage of an acquisition and takes about 45 minutes; the remaining acquisition is automated, and throughput is determined by the chosen resolution, frame averaging, and the number of samples. For lower resolution tasks, such as mapping electrode tracks in mouse brains, up to six brains can be imaged in 24 hours. For higher resolution tasks, such as cell counting in mouse brains, up to four brains can be imaged in the same time period, depending on the degree of frame averaging desired (Figure 2–figure supplement 1). Acquisition time per brain is roughly 4–15 hours depending on the resolution and number of samples imaged simultaneously (Figure 2– figure supplement 1C). The imaging, sectioning, and throughput parameters we routinely use are summarised in Supplementary Table 5.

Assembly of individual tiles into stitched planes is handled by our StitchIt software package (see Methods Section 4.5). Data are stitched automatically at the end of acquisition; during acquisition, StitchIt also generates a temporary stitched preview that is streamed to a website, allowing remote monitoring of the acquisition.

### 2.4 BakingTray for volumetric imaging

BakingTray enables high-quality 3D imaging of a variety of samples, including mouse and rat brains, songbird brains, and mouse kidneys and spleens (Fig. 4 and Figures 4–figure supplement 1, 4–figure supplement 2, & 4–figure supplement 3). The approach is compatible with any sample that can be agarose-embedded, cut cleanly on a vibratome, and imaged with a two-photon microscope. STPT produces seamless 3D datasets at cellular resolution (Fig. 4A, Figures 4–figure supplement 1 & 4–figure supplement 2), with three fluorescent labels excited simultaneously at 780 nm using tdTomato or mCherry, GFP, and BFP (Fig. 4B). Fixed tissue exhibits rich autofluorescence under two-photon excitation, so label-free structural imaging (including second harmonic generation, Fig. 4C) is possible without additional staining. The autofluorescence signal can be used for automated registration to a standard atlas, which delineates brain areas without a dedicated cytoarchitectural stain such as DAPI, or to build templates for new atlases (e.g. Wang *et al*. 2020; Gustison *et al*. 2024; Sirmpilatze *et al*. 2026). We use the BrainGlobe tools (brainglobe.info), but the data are equally compatible with other atlas-mapping pipelines such as ANTsX (Tustison *et al*. 2025), D-LMBmap (Li, Shang, *et al*. 2023), and BIRDS (Wang *et al*. 2021). We typically image samples that were already fluorescently labelled *in vivo*, whether by transgene expression, virus injection, or dye-labelled electrodes.

**Figure 4.**
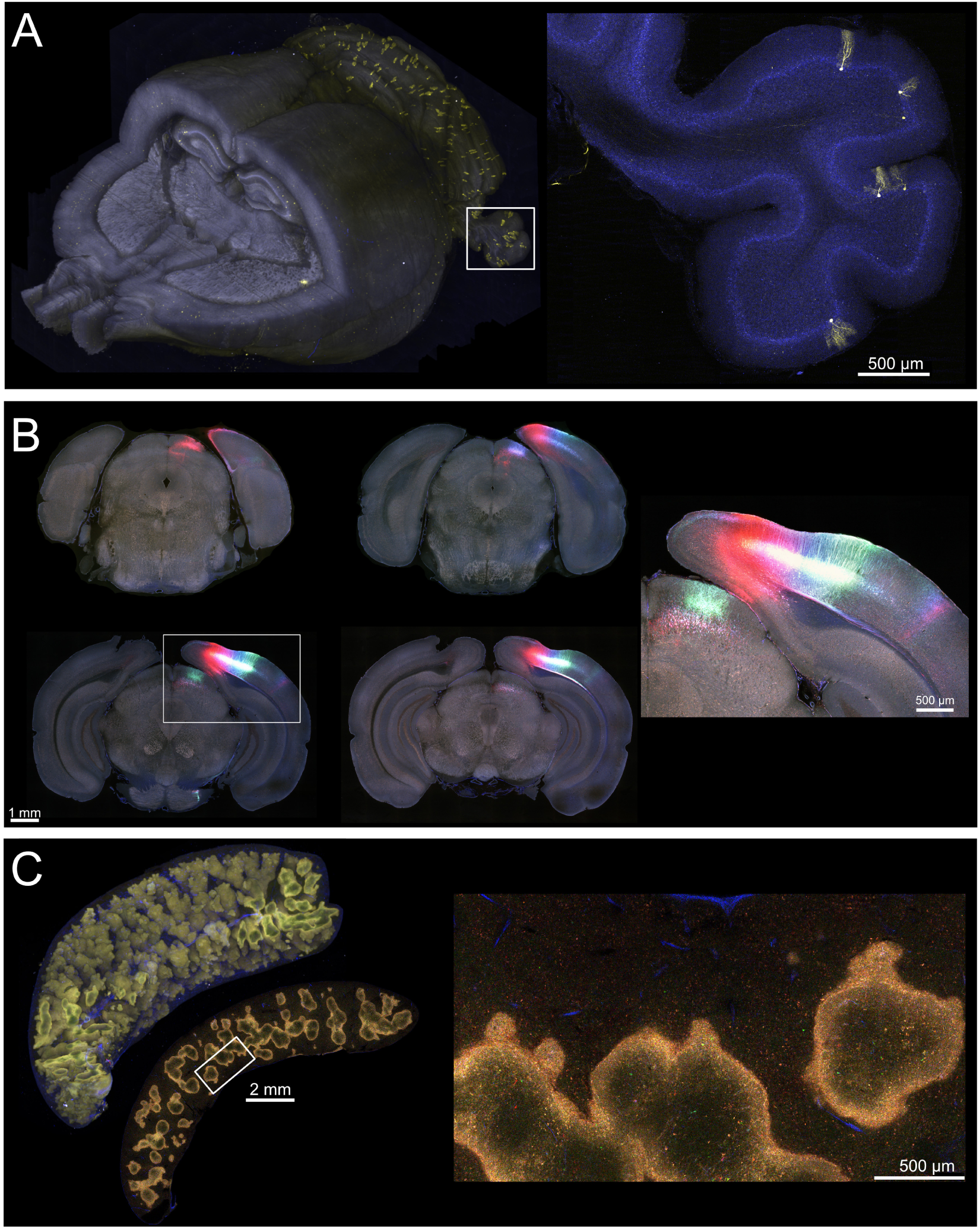
High-quality volumetric imaging. **A**. Rat brain sparsely expressing mNeonGreen (yellow); autofluorescence shown in blue. Right: a 100 *µ*m maximum intensity projection showing sparse labelling of individual Purkinje cells. Data from Chen *et al*. 2023: mNeonGreen expression delivered by systemic AAV injections. **B**. Four optical planes from a mouse brain labelled with three AAV injections in layer 5 neurons of primary visual cortex: mCherry, GFP, and BFP imaged at 780 nm. Inset panel shows boxed region at higher resolution. **C**. Mouse spleen with sparse confetti labelling. A 3D view of the organ is shown at the top left, adjacent to a representative 2D slice. A detail of that slice is shown on the right. The white pulp has stronger autofluorescence and appears yellow in the 3D panel. Collagen in blood vessel walls is an efficient second harmonic generator, producing blue photons (∼460 nm) from the same 930 nm femtosecond pulses used for two-photon imaging.

### 2.5 STPT imaging of whole-mount labelled tissue

Targets with no *in vivo* label, such as endogenous proteins or disease pathology, can be whole-mount stained before STPT imaging. We show two proof-of-principle examples: an antibody stain and a histological dye.

Figure 5A–C shows labelling with an antibody against the enzyme tyrosine hydroxylase (TH), conjugated to the far-red fluorescent dye Vio667. Far-red fluorescent proteins are less commonly used, so on a rig with a far-red channel, Vio667 staining can supplement genetically encoded or *in vivo* labels without competing for their detector channels. The horizontal plane in Fig. 5A is dominated by the dense TH-positive axonal terminals in the striatum, as expected. The midbrain dopaminergic nuclei that project to the striatum are visible at the top of the image, and seen again in the sagittal cross-section (Fig. 5B). The coronal plane (Fig. 5C) is at the level of the substantia nigra pars compacta. Labelling is high contrast, with good penetration and uniformity, and no rim of non-specific labelling around the perimeter of the tissue.

**Figure 5.**
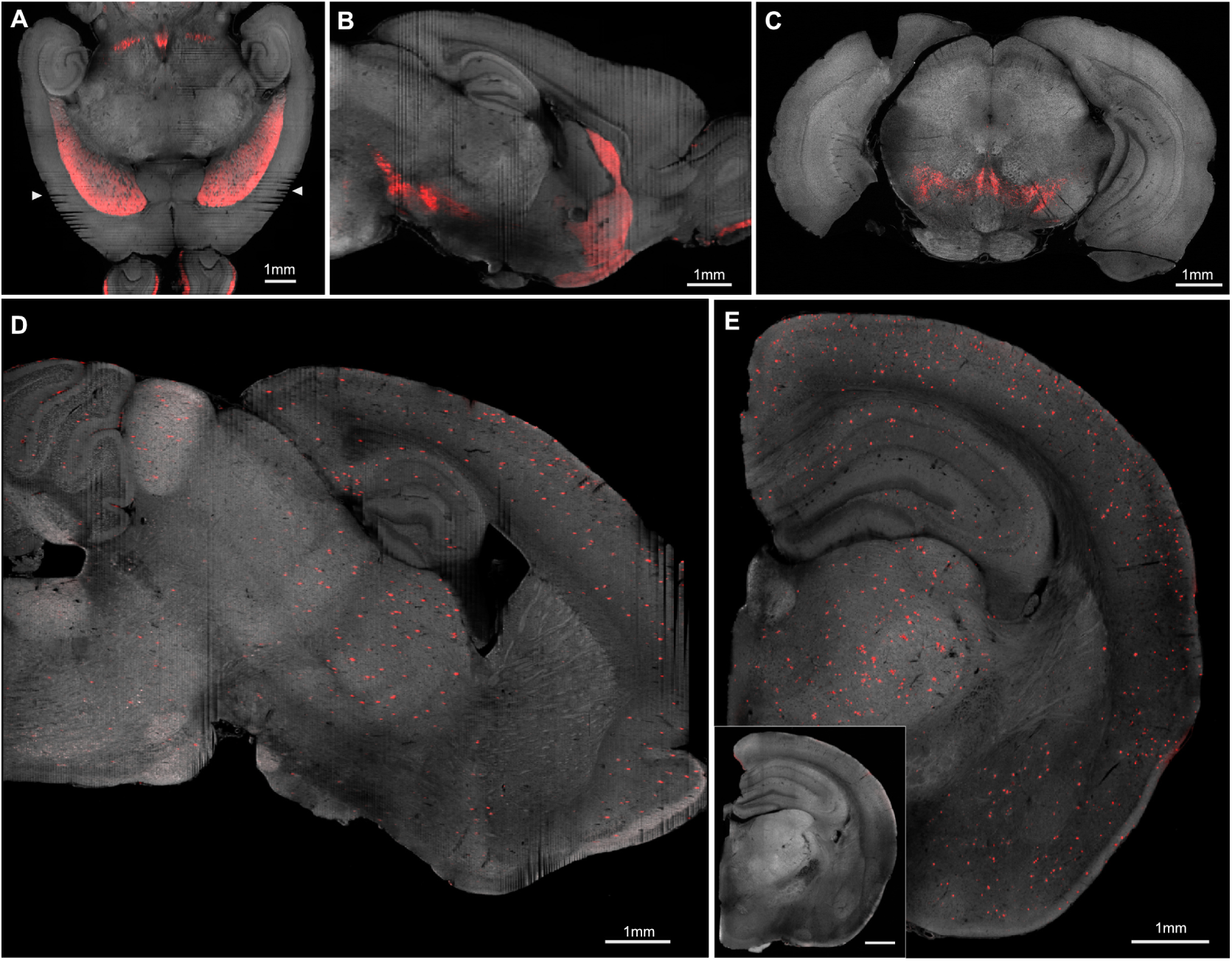
Whole-mount labelling. **A**. Labelling against tyrosine hydroxylase (TH) using a primary antibody conjugated to Vio667. Horizontal view obtained from a re-slice of a 25 *µ*m/voxel isotropic downsampled stack. Autofluorescence channel (green) shown in greyscale and the far-red channel showing Vio667 signal in red. Arrows indicate sectioning artefacts due to the permeabilised tissue being softer than a typical brain. **B**. As A but sagittal view. **C**. Coronal view downsampled to 4.4 *µ*m/pixel from 2.2 *µ*m/pixel raw data for display purposes. The far-red channel has very little autofluorescence so was passed through Fiji’s ‘de-speckle’ filter to clean up shot noise. **D**. Amyloid plaques labelled with Congo red in a half-brain. This sagittal view is obtained from a re-slice of a 10 *µ*m/voxel isotropic downsampled stack (original pixel spacing: 2.0 × 2.0 × 5 *µ*m). Autofluorescence channel (blue) shown in greyscale and the red signal channel containing Congo red labelling is overlaid. **E**. Coronal section derived from the same 10 *µ*m/voxel stack shown in D. Abundant plaque labelling is visible. Inset panel shows control brain processed the same way and with a similar look-up table. There is no evidence of plaque labelling in this brain. All scale bars are 1 mm.

To allow for whole-mount antibody labelling, the brain was delipidated using EZ Clear (Hsu *et al*. 2022) and labelled using the approach from iDISCO (Renier *et al*. 2014). The full protocol, which takes roughly two weeks, is described in Appendix 6. Tissue processed this way becomes softer and therefore harder to slice evenly. The arrows in Fig. 5A point at slicing artefacts caused by tissue movement relative to the agarose block; the same is visible in other panels. The brain was cut using our standard sectioning protocol, which was not adapted for this softer tissue. Slower cutting, softer agarose, and strengthening the tissue with gelatin (see Appendix 7) would therefore likely help. Alternatively, the iDISCO delipidation protocol produces firmer brains that slice more reliably.

Amyloid plaques, a hallmark of Alzheimer’s disease (AD), can be stained with Congo red. Half-brains were processed to reduce labelling time. The tissue was delipidated with CUBIC-L prior to staining, using a protocol that takes just under two weeks (Appendix 7). We labelled two groups of aged (22-month-old) mice: *App*^NL-G-F^ knock-in mice (Saito *et al*. 2014), which carry a humanised gene bearing mutations linked to familial AD, and *App*^hu/hu^ mice (Serneels *et al*. 2020), their age-matched control, which carries the humanised gene without the mutations. One mouse from each group is shown here; the animals formed part of a larger behavioural study (in preparation) comparing humanised AD models. An example labelled *App*^NL-G-F^ brain is shown in Fig. 5D&E; plaques (red) are profuse, spread brain-wide, and readily distinguished from background autofluorescence (grey). The age-matched control (inset Fig. 5E) shows no plaque labelling. Plaques can also be labelled *in vivo* with methoxy-X04 (Klunk *et al*. 2002), and such brains would feed directly into an STPT pipeline without the post-mortem staining step (e.g. Whitesell *et al*. 2019; Oblak *et al*. 2021).

Whilst BrainSaw is compatible with whole-mount labelling, the much longer preparation times and the challenges of sectioning softer tissue mean that STPT works best with tissue labelled *in vivo*.

### 2.6 Main use cases

We categorised the 35 publications that used BakingTray (Supplementary Table 4) to find the most common use cases of the approach. BakingTray was most often used to map the location of dye-labelled electrode tracks (13 publications) or fibre tracks (two publications). Whole-brain cell counting was conducted in 10 publications, and finally researchers mapped bulk projections (five publications) and lesions (one publication). The remaining use cases were mostly methods-adjacent, such as atlas generation, screening AAV expression, or hardware development. Below we illustrate the three most common applications.

#### Use case – Electrode tracks

Electrodes used for *in vivo* recordings are dipped in a lipophilic dye, such as CM-DiI, which remains after the electrode is withdrawn. The animal is perfused and the brain is imaged via STPT to precisely map the location of the dye track. We image such samples at lower resolution (often 4 × 4 × 20 *µ*m/voxel; about 6 GB per channel for a mouse brain), and typically process 4–6 in a single acquisition. The electrode tracks can be traced and the brain registered to a standard atlas (Fig. 6A). This objectively locates electrode tracks in anatomical space and allows researchers to overlay the locations of different tracks in the same standard space, as done at scale by the International Brain Laboratory (Int. Brain Lab. *et al*. 2025, Fig. 6A&B), where the locations of 699 Neuropixels probe insertions across 139 mice were mapped using BrainSaw.

**Figure 6.**
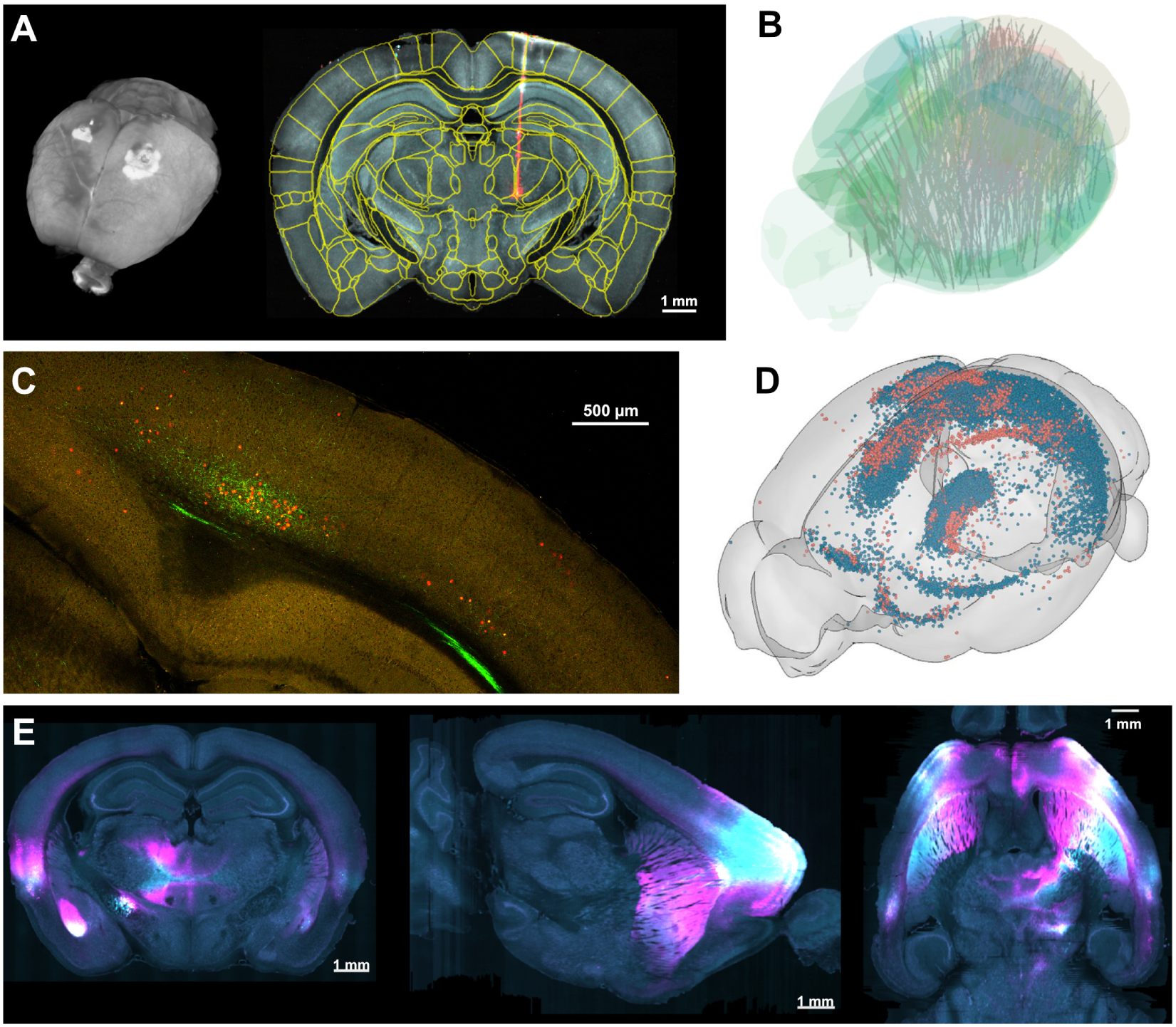
Main use cases. This figure shows example raw data underlying the three most common use cases for BakingTray. **A**. Raw data from an individual brain containing a DiI-labelled electrode track. A 3D view of the whole brain is shown to the left and a section registered to the Allen mouse brain atlas containing the electrode track is shown to the right. Data in A are from one animal that contributed to the IBL Brain-Wide Map (Int. Brain Lab. *et al*. 2025). **B** shows all 699 electrode tracks overlaid in atlas space. **C**. Detail from a section imaged at 2 × 2 *µ*m/pixel for a cell-counting experiment. The nuclear-targeted H2B-tdTomato cells (red) are to be automatically counted by the cellfinder pipeline. The processes of a separate neural population filled with eGFP are shown in green. This image is intended to provide the reader with an expectation for what a good-quality, ‘cellfinder-ready’ dataset looks like. **D**. The coordinates of mapped neurons from different animals (red and blue) can be overlaid in atlas space and analysed together, adapted from Figure 1 of Tyson and Margrie 2022. **E**. Three orthogonal views of mouse brain with two large virus injections leading to expression of eGFP (cyan) and mCherry (magenta). Overlapping regions are rendered white. Images are derived from the isotropic 25 *µ*m/voxel downsampled stacks generated at the end of acquisition. This resolution is often adequate for quantifying data of this type. *Permissions*: panel **B** is adapted from Figure 2A of Int. Brain Lab. *et al*. 2025 and panel **D** from Figure 1 of Tyson and Margrie 2022. Both articles are published under a CC BY 4.0 licence.

#### Use case – Cell counting

STPT datasets allow users to quantify the number and distribution of labelled cells across the whole brain using automated pipelines. For this application, brains are imaged at 2 × 2 × 5 *µ*m/voxel (40 *µ*m sections, eight optical planes). At these higher resolutions, 2–4 brains can be imaged in 24 hours and the resulting data occupy roughly 120 GB per channel per mouse brain. Figure 6C shows a detail from a single plane of one such acquisition; one population of neurons expresses nuclear-targeted tdTomato (red); the other is filled with eGFP (green). The BrainGlobe tool cellfinder (Tyson *et al*. 2021) can be used to count all labelled cells brain-wide and display these in atlas space (Fig. 6D). The same pipeline works in other species given a suitable reference atlas: we ran cellfinder on a rat brain sparsely labelled by systemic AAV injection (as in Fig. 4A) and registered the detected cells to a rat brain atlas (Fig. 6–figure supplement 1). Further cell-counting data are provided in Results Section 2.7, below.

#### Use case – Bulk projections

Projections of *populations* of labelled neurons can be mapped without tracing individual axons. Single-cell axonal tracing is time-consuming and requires very large datasets (typically 5–30 TB), so it is only appropriate if the scientific question demands it (see Discussion Section 3.2). Bulk-labelling data are acquired at around 4 × 4 × 20 *µ*m/voxel to 2 × 2 × 20 *µ*m/voxel (roughly 6–24 GB per channel per mouse brain). Projection locations can be established simply by thresholding for expression. Figure 6E shows an example with two viral injections; the data are displayed as three orthogonal views at 25 *µ*m isotropic voxels. Even this low resolution can be adequate for quantifying data of this type, accelerating analysis (e.g. see Esmaeili *et al*. 2022 Fig. 5, and El-Boustani *et al*. 2020 Fig. 1).

### 2.7 Dual-colour cell counting

Comparing the brain-wide inputs to two regions is most powerful when both populations are labelled in the same animal, removing inter-animal variability. Here we demonstrate that two differently labelled neuronal populations can be counted in one brain imaged on BrainSaw. We used nuclear-targeted retrograde viral tracing with rAAV2-retro-EF1a-H2B-eGFP and rAAV2-retro-EF1a-H2B-tdTomato (Tervo *et al*. 2016; Weiler, Teichert, and Margrie 2025), injected adjacent to one another along the medial-lateral axis of primary visual cortex in mice (Fig. 7A&B). tdTomato and eGFP are generally well co-excited at a single wavelength around 920–950 nm (Figure 7–figure supplement 1A), with rich tdTomato signal often seen even at the shorter end of the range (Figures 4– figure supplement 1 & 4–figure supplement 2). Nonetheless, tdTomato can optionally be boosted by adding a second laser line at around 1030–1090 nm. In this pilot dataset we co-aligned a 1064 nm laser and scanned it simultaneously with the system’s main tunable line set to 920 nm to demonstrate multi-wavelength imaging. We then performed dual-colour cell detection using cellfinder (Fig. 7C&D), which correctly rejected the bleed-through of strong eGFP emission into the adjacent red channel (Figure 7–figure supplement 1B). This approach allowed us to visualise the topographic distribution and relative fraction of retrogradely labelled neuronal nuclei across different brain areas projecting to two neighbouring sites within primary visual cortex (Fig. 7E&F).

**Figure 7.**
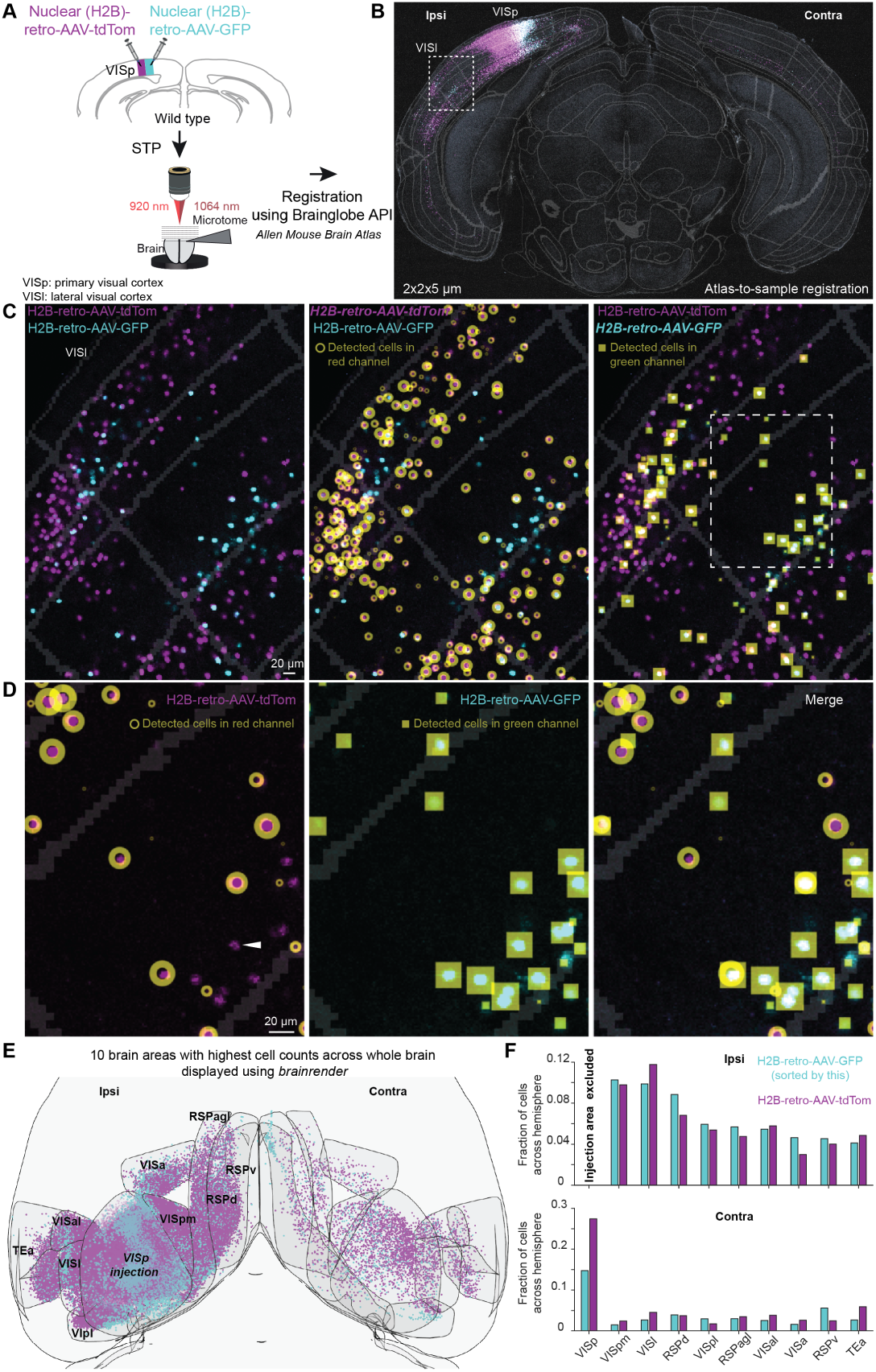
Dual-colour nuclear retrograde brain-wide tracing. **A**. Schematic depicting the dual-colour viral tracing approach. After viral expression, the entire brain was cut and simultaneously imaged at 920 and 1064 nm using STPT. **B**. Atlas-to-sample registration. Raw two-photon image overlaid with the brain region segmentation boundaries based on the Allen Mouse Brain Common Coordinate Framework, showing injection sites and labelled nuclei of retro-GFP (cyan) and retro-tdTom (magenta) labelled cells. **C**. Zoomed-in area, highlighted in B, showing retro-tdTom+ and retro-GFP+ labelled nuclei in area VISl and neighbouring areas (left), alongside automatically detected retro-GFP+ (middle) and retro-tdTom+ (right) nuclei using cellfinder. **D**. Zoomed-in area, highlighted in C, displaying the tdTom+ channel (left), the GFP+ channel (middle) and the merge of both channels (right) together with detected nuclei. The white arrowhead indicates bleed-through of a strong GFP+ signal into the tdTom+ channel, without being incorrectly detected as a tdTom+ nucleus. **E**. Visualisation of detected retrogradely labelled nuclei warped into the 3D rendered space of the two hemispheres. Note that only the ten brain areas with the highest nuclei counts are displayed. **F**. Bar graphs of the normalised fractional cell counts for each of the ten brain areas shown in E, for ipsi- and contralateral hemispheres. Fractional counts are ranked by retro-GFP+ nuclei.

### 2.8 Mapping synaptic sites brain-wide

Tracing with a cytoplasmic fluorophore reveals neurons’ axonal spread, but not where they are likely to make synaptic contacts. The locations of putative synaptic sites can be revealed using fluorescent labelling approaches, such as expression of synaptophysin-eGFP (Li, Tasic, *et al*. 2010). In an analogous way to bulk tracing of axons (Fig. 6E), STPT can be used to map the distribution of synaptophysin-eGFP without resorting to extreme magnifications (here we use 2 × 2 × 5 *µ*m/voxel) or counting individual terminals. This allows the whole brain to be acquired at a moderate resolution whilst capturing both axonal projection patterns and the distribution of putative synaptic output, with the data placed in a common atlas space. We demonstrate this in the ventral lateral geniculate nucleus (vLGN), a prethalamic nucleus containing at least 18 inhibitory neuronal cell types (Stebbins *et al*. 2026) that form long-range projections throughout the brain. Whilst axonal spread from the vLGN has been mapped previously (Moore, Weis, and Moga 2000; Fratzl *et al*. 2021; Salay and Huberman 2021; Li, Peng, *et al*. 2023; Mederos *et al*. 2025), it has not been broken down by cell type, and the locations of putative synaptic sites have not been assessed. The vLGN is therefore a good target for illustrating the benefits of this synaptic labelling approach. We do so here descriptively, using one animal per driver line; quantification is beyond the scope of this manuscript.

We virally labelled neurons with AAV[2/1]-hSyn-DIO-tdTomato-Synaptophysin-eGFP, which fills somata, dendrites, and axons with tdTomato while simultaneously marking putative presynaptic terminals with synaptophysin-eGFP (Fig. 8A; Methods Section 4.4). This virus is Cre-dependent, which allowed us to target its expression to defined cell types using Cre driver lines: vesicular GABA transporter (vgat), parvalbumin (PV), proenkephalin (Penk), and somatostatin (SOM). We show example images from each driver line to demonstrate combined mapping of axons and putative synaptic sites (Fig. 8). In all cases we registered data to the Enhanced and Unified Mouse Brain Atlas (Chon *et al*. 2019) using the BrainGlobe tools, allowing synaptic output patterns to be compared based on objectively defined brain region borders (Fig. 8B&C).

**Figure 8.**
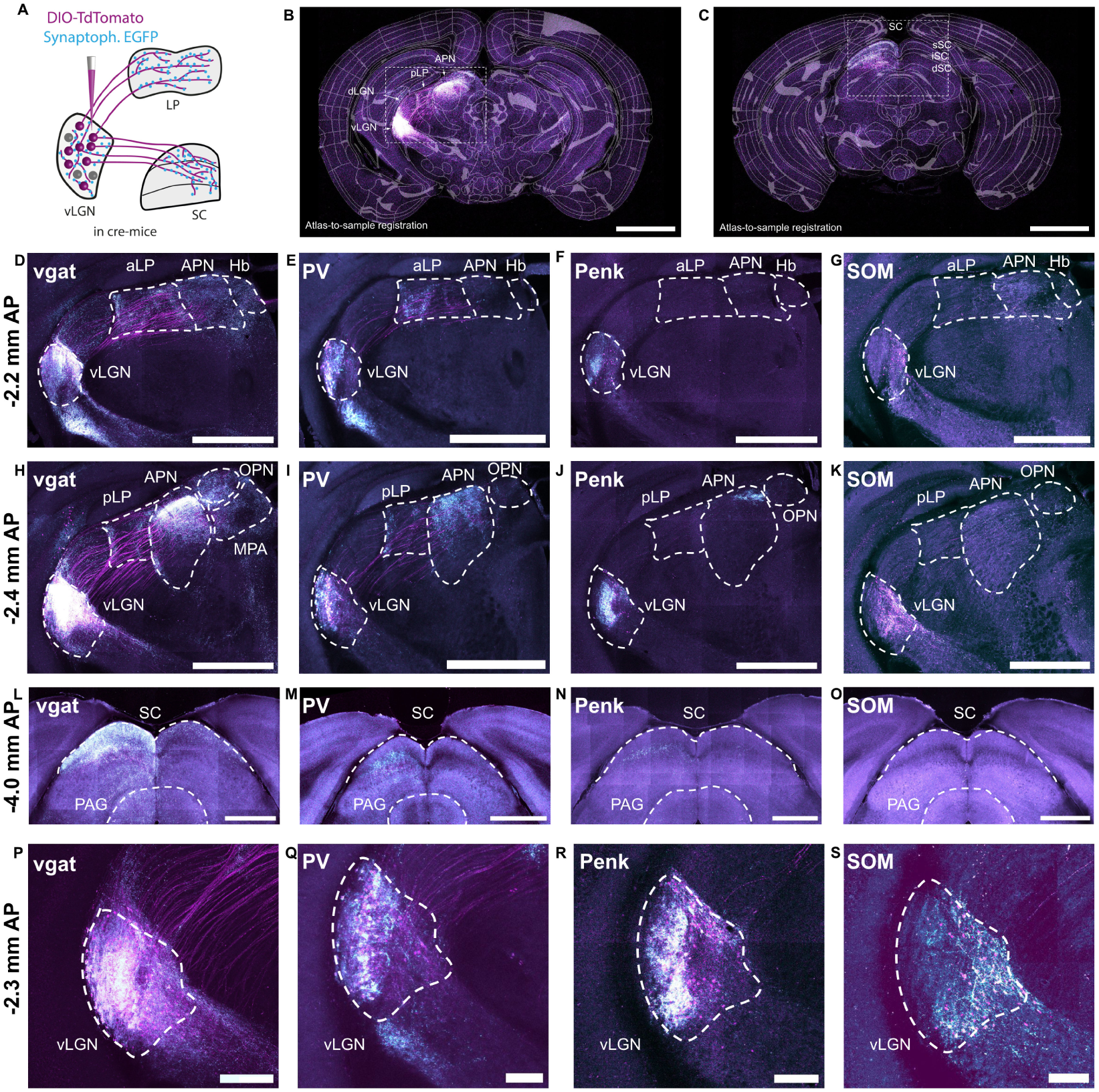
Brain-wide synaptic mapping. **A**. Injection of a Cre-dependent virus into the vLGN fills neurons with tdTomato and labels presynaptic terminals using synaptophysin-eGFP. **B**. Atlas-to-sample registration. Raw two-photon image overlaid with the brain region segmentation boundaries based on the Enhanced and Unified Mouse Brain Atlas, showing the injection site in the vLGN and labelled somata and axons in tdTomato (magenta) and their presynaptic terminals in eGFP (cyan). Scale bar is 500 *µ*m. **C**. As B but showing projections and putative presynaptic terminals in the SC. Scale bar is 500 *µ*m. **D–G**. Local and long-range projections shown by expression of the virus in vgat, PV, Penk, and SOM vLGN neurons in the region of the anterior LP. Scale bar is 500 *µ*m. **H–K**. As D–G but in the region of the posterior LP. **L–O**. Long-range projections to the SC. Scale bar is 500 *µ*m. **P–S**. Local presynaptic labelling within the vLGN. Scale bar is 200 *µ*m. Further images from a single animal are shown in Figure 8–figure supplement 1. *Abbreviations:* APN, anterior pretectal nucleus; dLGN, dorsal lateral geniculate nucleus; dSC, deep layers of the superior colliculus; iSC, intermediate layers of the superior colliculus; LP, lateral posterior nucleus of the thalamus; SC, superior colliculus; sSC, superficial layers of the superior colliculus; vLGN, ventral lateral geniculate nucleus.

Our data show clear examples of regions through which axons merely pass, as opposed to those where putative synaptic contacts exist (e.g. Fig. 8H&I, and Fig. 8–figure supplement 1). Projection patterns in the ipsilateral thalamus and pretectum differed across inhibitory cell classes (Fig. 8D–K). The general population of vgat-expressing GABAergic neurons formed putative synaptic sites in multiple thalamic and pretectal nuclei, in particular the anterior pretectal nucleus (APN), while subtypes of GABAergic neurons, such as PV- and Penk-expressing neurons, appeared to only innervate a subset of these regions. Within the superior colliculus (SC), likewise, vgat-, PV-, and Penk-expressing neurons all formed putative synaptic contacts but targeted different subregions (Fig. 8L–O).

The spatial resolution of the data was sufficient to examine local presynaptic labelling within the vLGN itself. Local synaptophysin-eGFP puncta were present in all four cell types, but their distribution within the vLGN differed. Neurons in vgat-Cre mice exhibited extensive local axonal labelling, together with dense synaptophysin-eGFP puncta throughout the vLGN, indicating sub-stantial local inhibitory output (Fig. 8P). The remaining three groups had different distributions of synaptophysin-eGFP puncta (Fig. 8Q–S), with SOM-expressing neurons having puncta and somata located most medially. These observations illustrate the local and long-range connectivity patterns of defined vLGN inhibitory subtypes, and demonstrate the strength of STPT for studying synaptic organisation across multiple spatial scales within a single brain.

## 3 Discussion

We have presented BakingTray, a complete open-source acquisition platform for STPT, with an established user base and proven track record. This manuscript has focused on the software, as it is the enabling factor for opening up STPT to the DIY builder community, and on how the resulting systems perform in practice. We describe the hardware only briefly, as this has been documented previously (Ragan *et al*. 2012; Zheng *et al*. 2013; Economo *et al*. 2016; Seiriki *et al*. 2019) and our BrainSaw systems are not of a single design. The critical aspect of the hardware is how the imaging plane, three-axis stage, and vibratome are co-aligned. We describe this process in Appendix 4, have made available CAD drawings detailing the relevant key features, and provide online documentation to serve as a dynamic information repository. BakingTray runs alongside StitchIt (stitchit.swcmicroscopy.com), which performs various pre-processing operations during acquisition and assembles image tiles into stitched planes. Together, BakingTray, StitchIt, and the BrainSaw hardware form an ecosystem that delivers STPT from acquisition to downstream analysis. Here we put these tools into context, in particular exploring the limitations of the approach and future directions of the project.

### 3.1 Building on ScanImage

The first development decision was whether BakingTray should acquire images itself or delegate the task. Basic scanning software is trivial to write (e.g. SimpleMScanner), but software that is stable, well tested, and supports a broad range of acquisition hardware is a far larger undertaking. BakingTray was therefore built on ScanImage, which had already solved that problem. This confined the scope of the project to tile-scanning, sample detection, slicing, and a smooth user experience. ScanImage runs in MATLAB, so BakingTray does too. This has proved an advantage for longevity: MATLAB introduces very few breaking changes between releases, making maintenance easy.

BakingTray therefore depends on two commercial products: MATLAB and ScanImage. An annual academic MATLAB licence costs about the same as the SWC’s annual spend on BrainSaw vibratome blades. The ScanImage licence is more expensive, but buys professional support at a price well below a commercial microscope service contract. Since low-level hardware control libraries (NI-DAQmx, PI’s motion control APIs, etc.) are closed source whatever language is chosen, we did not view the proprietary nature of MATLAB and ScanImage as a constraint. BakingTray and StitchIt are themselves released under an open-source licence.

### 3.2 Where BrainSaw fits

We use BakingTray for moderate-resolution, high-throughput volumetric imaging where the primary consumer of the data is often an automated analysis pipeline rather than a human observer. BrainSaw systems excel at resolutions adequate for electrode track localisation, cell body counting, and bulk projection mapping – the workhorse applications of systems neuroscience. At these resolutions, datasets are generally modest in size, and multiple samples can be imaged overnight on a single microscope. This allows samples from different experimental groups to be imaged simultaneously under the same conditions.

Although brain-wide tracing of individual axons is achievable with STPT (Zheng *et al*. 2013; Han *et al*. 2018; Winnubst *et al*. 2019) and BakingTray (Liu *et al*. 2022), we have not optimised our platform for the sub-micron voxel sizes this requires, as there has been little demand for it within our user community. Systems designed for single-axon tracing, such as the ExA-SPIM (Glaser *et al*. 2025; Su *et al*. 2026), are probably the best choice at present.

Despite our focus on neuroscience applications, STPT can be used to image other structures such as spleens (Fig. 4C, Li *et al*. 2021), prostate glands and livers (Taranda *et al*. 2021), tumours (Grauel *et al*. 2020), and kidneys (Figure 4–figure supplement 3). Any organ or structure that cuts well on a vibratome can be imaged on a BrainSaw system (see fresh carrot, Figure 4–figure supplement 3C).

### 3.3 System reliability

In Results Section 2.2 we quantify the reliability of the three BrainSaw systems at the SWC by analysing the number of times they failed to complete an acquisition unattended. Reliability data were gathered only from the SWC because this was the one site where we had ready access to a large pool of data. It is unlikely users are simply not uploading failed acquisitions to the storage platform: resuming an acquisition is straightforward and there is no motivation for discarding partial acquisitions and losing data. We have no reason to think reliability differs at other sites, with one known exception: a site that uses Zaber linear stages for tile-scanning found these need regular lubrication (Methods Section 4.1) and tend to stall, stopping the acquisition.

The acquisition statistics cannot capture more subtle errors, such as users setting up samples incorrectly, thereby producing the wrong data or data of poor quality. Less experienced users or those in a rush do make errors and this underscores the need for good user training and a vigilant system administrator or ‘superuser’, who is also responsible for alignment, calibration, and troubleshooting. To assist with this we provide extensive documentation and additional resources.

### 3.4 Who should build a BrainSaw?

A BrainSaw is not a turnkey instrument. It requires building – or at least maintaining – a high-quality two-photon microscope, involving alignment of dangerous Class IV lasers. An STPT system as a whole is composed of many interacting parts, and will require regular oversight for trouble-free long-term operation. The need for expert oversight is not unusual; the same is true of many lightsheet and other high-end commercial microscopes.

For those with the appropriate expertise, BakingTray not only solves the crucial hardware control problem but also provides a well-tested acquisition process that would otherwise take many months of development to achieve. Operators can proceed directly to hardware design and assembly, knowing that BakingTray’s modular architecture means that hardware choices are not dictated by the software. In addition to this manuscript, we make available online documentation (bakingtray.swcmicroscopy.com) to help with construction. The corresponding author can be contacted for assistance with planning, building, or running a BrainSaw.

Prospective adopters should also plan for analysis and storage infrastructure. Datasets range in size from about 6 to 120 GB per channel per sample, depending on the resolution chosen (Supplementary Table 5). We use a Linux workstation with 12–24 TB of local storage for stitching and pre-processing. This PC in turn sends data to a managed storage platform for long-term storage and tape archiving.

### 3.5 Limitations

Because STPT is destructive, one cannot re-acquire samples that were imaged incorrectly. Settings must be correct at the start and obvious issues, such as inconsistent sectioning, must be resolved before initiating acquisition (Appendix 5). End-users less familiar with two-photon imaging sometimes acquire data with obvious defects (e.g. laser power being too low) without realising. Whilst problems associated with user inexperience are not unique to STPT, they are particularly important to eradicate due to the technique’s destructive nature.

BrainSaws have no way of automatically collecting slices as they come off the block face. Users occasionally gather slices post-acquisition in order to antibody stain and image a small subset of sections. Those needing routine *post hoc* labelling of many sections could instead stain a smaller number of thicker cleared slices (e.g. 500 *µ*m sections) and image them on a confocal.

Most users image samples labelled with genetically encoded fluorophores or organic dyes deposited *in vivo*. One of the main advantages of STPT is the minimal sample preparation, making such samples a natural fit. Whilst whole-mount labelling followed by STPT is possible (see Results Section 2.5), and increases the flexibility of the technique, it remains technically demanding to do well and we do not perform it routinely. Groups whose experiments depend on whole-mount labelling are likely better served by clearing and lightsheet imaging, for which these protocols were developed.

Samples that image poorly on an STPT system include large or gyrencephalic brains, or those that are too soft to be reliably cut on a vibratome. Brains from larger animals, such as ferrets, have thick dura, which is impossible to cut with a vibratome; it forms long streamers that occlude the sample surface causing shadowing artefacts, and complete dura removal is not possible given the folded cortical surface. Rat brains are more prone to dural shadows than mouse brains, especially around the caudal hippocampus where membranes emerge from the ventricles mid-acquisition. Mouse brains cut well, although a small minority show shadowing artefacts when dura is not removed prior to imaging (Fig. 9). Soft samples, such as embryos, can be challenging to cut and may pop out of the agarose prematurely or shift during slicing. Whilst there are techniques for covalently cross-linking samples to agarose (Ragan *et al*. 2012), they significantly lengthen the sample preparation time and are not completely effective at mitigating these issues. In our hands mouse and rat brains, and other organs of comparable size and firmness, can be imaged reliably and routinely.

**Figure 9.**
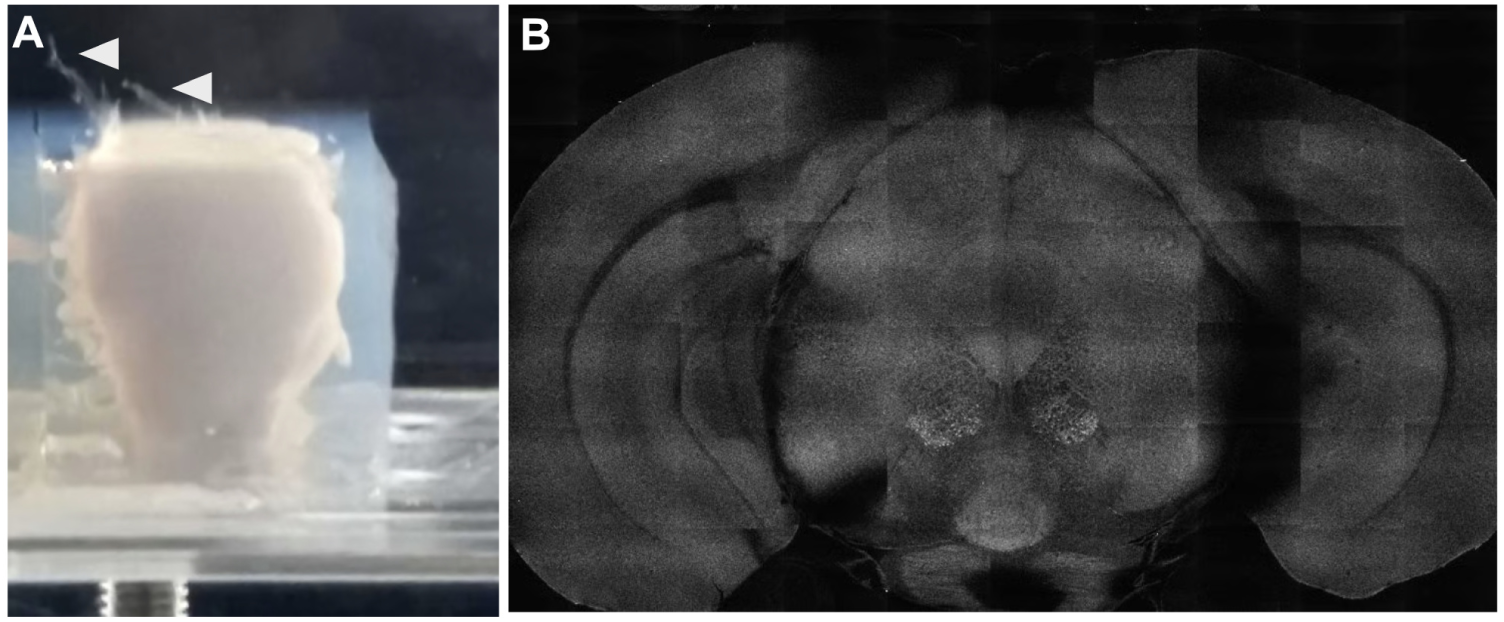
Dura not removed prior to imaging can cause shadowing artefacts even in mouse brains. **A**. A mouse brain embedded in agarose in the BrainSaw water bath. The dura was not removed prior to imaging and now extends in long streamers beyond the block face (arrowed). **B**. Detail taken during acquisition from the BakingTray GUI showing how the exposed block face in A appears once it has been imaged. A particularly prominent shadow can be seen to the right of the image. The tile pattern is visible because vignetting is not corrected at this stage of acquisition.

For simplicity, BakingTray does not track the absolute height of the block face and so cannot image a fixed distance below it. Remarkably, once the block has been correctly trimmed (Appendix 5), the distance between the objective and the block face usually shows no detectable drift over the course of an acquisition. Occasionally it drifts by 10–20 *µ*m, which is a substantial fraction of the 25–35 *µ*m depth at which we typically acquire the first optical plane. The user can correct this by adjusting the objective height live, without pausing or restarting the acquisition (see Appendix 3 for more information on which settings can be changed during acquisition). Automatic surface detection could be added to the software, but would need to be implemented carefully to avoid introducing more problems than it solves.

Laser power at the sample can be changed manually during acquisition (Appendix 3) but not in an automatic manner based on local signal strength. Bright viral injection sites may therefore become saturated (e.g. Fig. 4B, inset), as fluorophore density is very high in these areas. This is a constraint worth bearing in mind, although it is rarely a problem.

### 3.6 Maintenance and future directions

BakingTray is an integral part of the SWC’s mission and will be actively maintained for the foreseeable future by the Centre’s staff scientists, who are core-funded by the Wellcome Trust and Gatsby Charitable Foundation (see Funding). The growing base of installed systems also constitutes an additional level of support.

BakingTray is largely feature-complete, but we are constantly making usability improvements and bug fixes. The only significant new feature currently planned is the introduction of interleaved multi-laser acquisition within a single section. This will allow the same region to be imaged rapidly in sequence with different excitation lines and acquisition channels, minimising cross-talk. The approach will be an extension of the dual-laser excitation data presented in Results Section 2.7, where two laser lines were scanned simultaneously with no multiplexing.

We hope that by sharing BakingTray and the BrainSaw ecosystem we lower the barrier for groups wishing to adopt STPT and encourage a wider community of developers to extend the platform in directions we have not anticipated.

## 4 Methods

### Key resources table

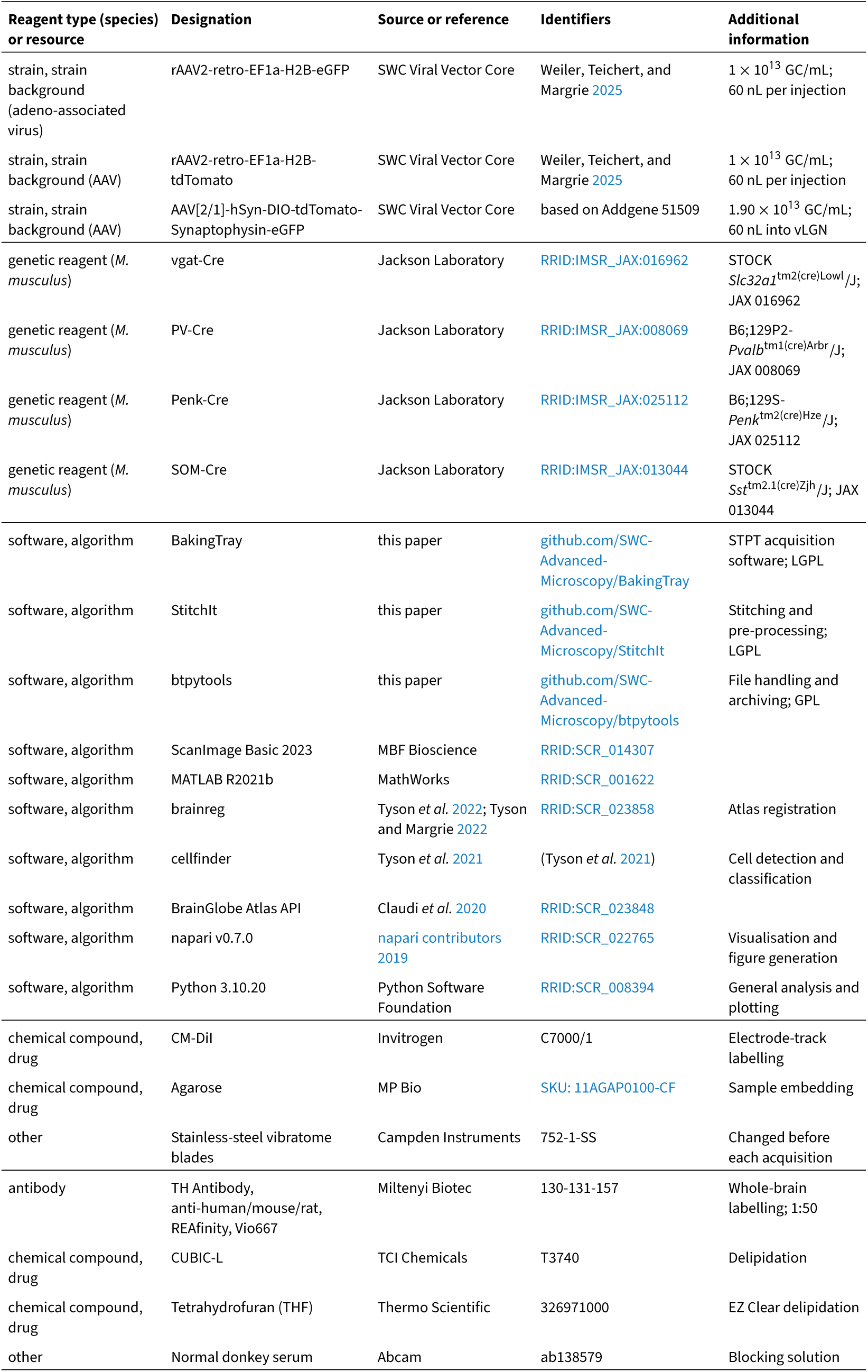
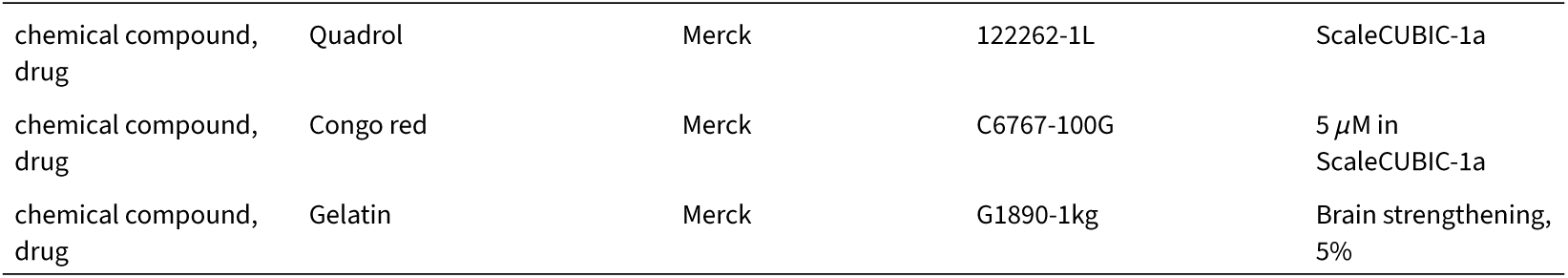

### 4.1 System hardware

Since BakingTray is not tied to a specific hardware configuration, the BrainSaw systems we have constructed are not identical to each other (Fig. 10). Below we provide an overview of the core hardware and highlight choices that are particularly effective. A CAD model of a generic system from the objective down is provided in our BrainSaw_CAD GitHub repository.

**Figure 10.**
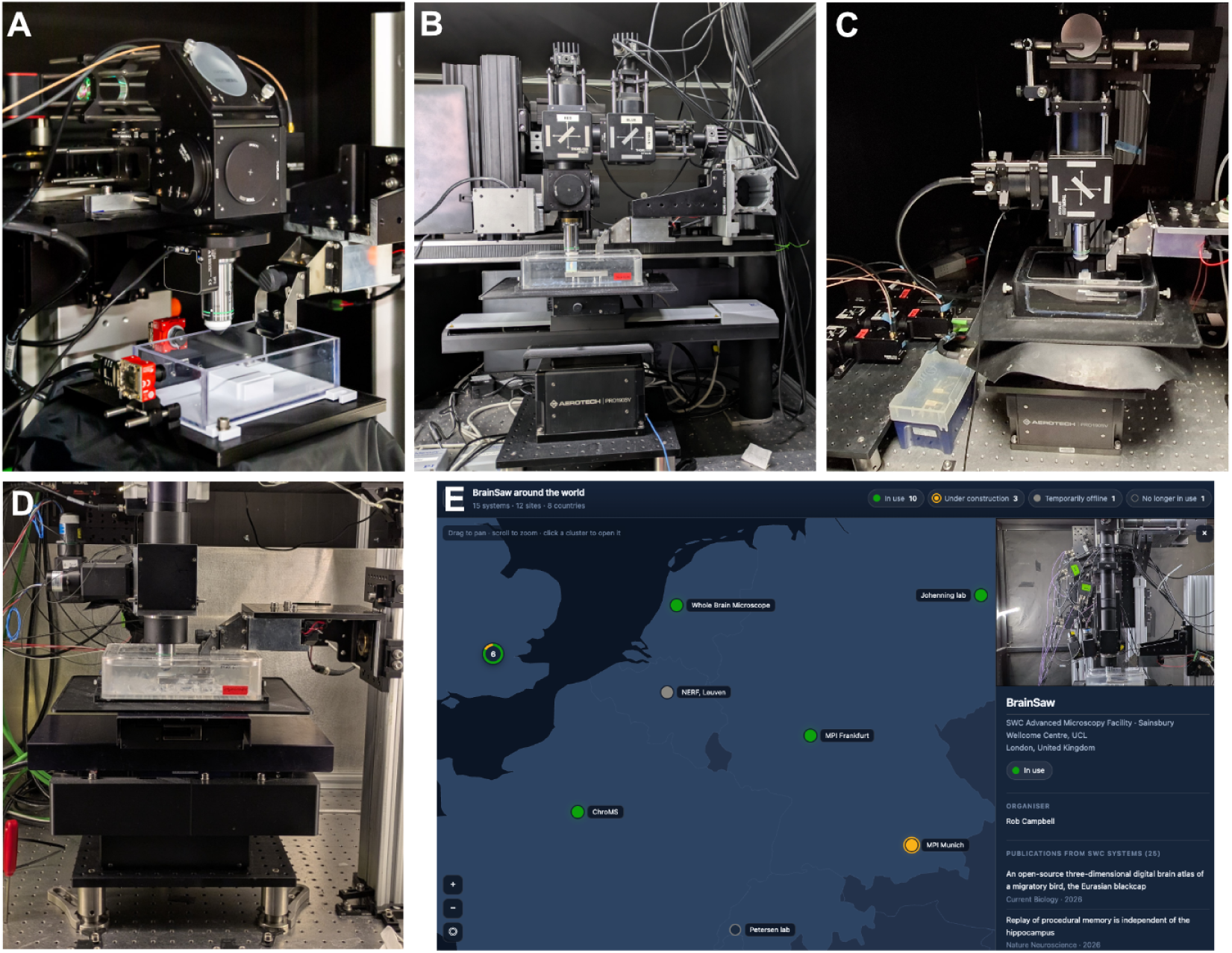
The diversity of BrainSaw systems. **A**. The Frankfurt Max Planck system has cameras on the water bath, which is of a unique design. **B**. The ‘NeuroVision’ system at the SWC is a modified TissueCyte 1000; it is the only one equipped with SiPMs. **C**. ‘CrickSaw’ at the Francis Crick Institute lacks tilt adjustment on the sample stage and has a liquid light-guide-based collection path. **D**. The ‘OrganGrinder’ system at the SWC is equipped with a single-line Axon 1064 nm laser in addition to a tunable Mai Tai eHP DS. **E**. Our interactive BrainSaw system map (found at swcmicroscopy.com/brainsaw) shows all systems built to date, their status, and associated publications.

#### Scanners and optics

We recommend resonant scanning over galvo scanning. Resonant scanners achieve higher frame rates at the cost of shorter pixel dwell times and therefore lower signal-to-noise ratio (Nguyen *et al*. 2001), but frame averaging is available within ScanImage for samples with lower fluorescence levels. Galvo scanning is supported but not recommended: galvo mirrors driven too fast are prone to variable bidirectional scan phase artefacts that cannot be corrected *post hoc*. We use 4 kHz or 8 kHz resonant scanners (Cambridge Technology or Thorlabs) with the slow axis galvo sourced from ScannerMax. We have systems with both close-coupled and optically relayed scan heads. Optical relays create a more uniformly illuminated field of view at the cost of more glass and additional complexity. We match the scan and tube lens choice to the objective back aperture in order to achieve the widest scan angle. Scan optics are generally assembled using Thorlabs lenses. We typically use Nikon W 16x/0.8 objectives, which provide a field of view of approximately 1.15 mm with roughly 20 *µ*m of field curvature. The Zeiss W Plan-Apochromat 20x/1.0 is a viable alternative for higher-resolution applications.

#### Detectors and pre-amps

Our systems use up to four detection channels (Fig. 11), which are acquired using either NI DAQ or ScanImage’s ‘vDAQ’ hardware. A range of different detector types are used across systems. Hamamatsu R10699 multi-alkali PMTs are adequate, but higher sensitivity can be achieved with GaAsP PMTs, such as Hamamatsu H16201P-40. GaAsPs without a trip circuit are preferred, as the trip can be triggered by bright sample regions. Note that GaAsPs are much more sensitive to degradation when exposed to higher light levels. Sensitivity of GaAsPs should be logged over time so that they can be replaced when necessary (Lees *et al*. 2025). Although multi-alkali PMTs such as R10699s have lower initial quantum efficiency than their GaAsP counterparts, we have not noticed them degrade over time (even after almost 10 years of service). We use Thorlabs TIA60 or Femto DHPCA-100 pre-amplifiers.

**Figure 11.**
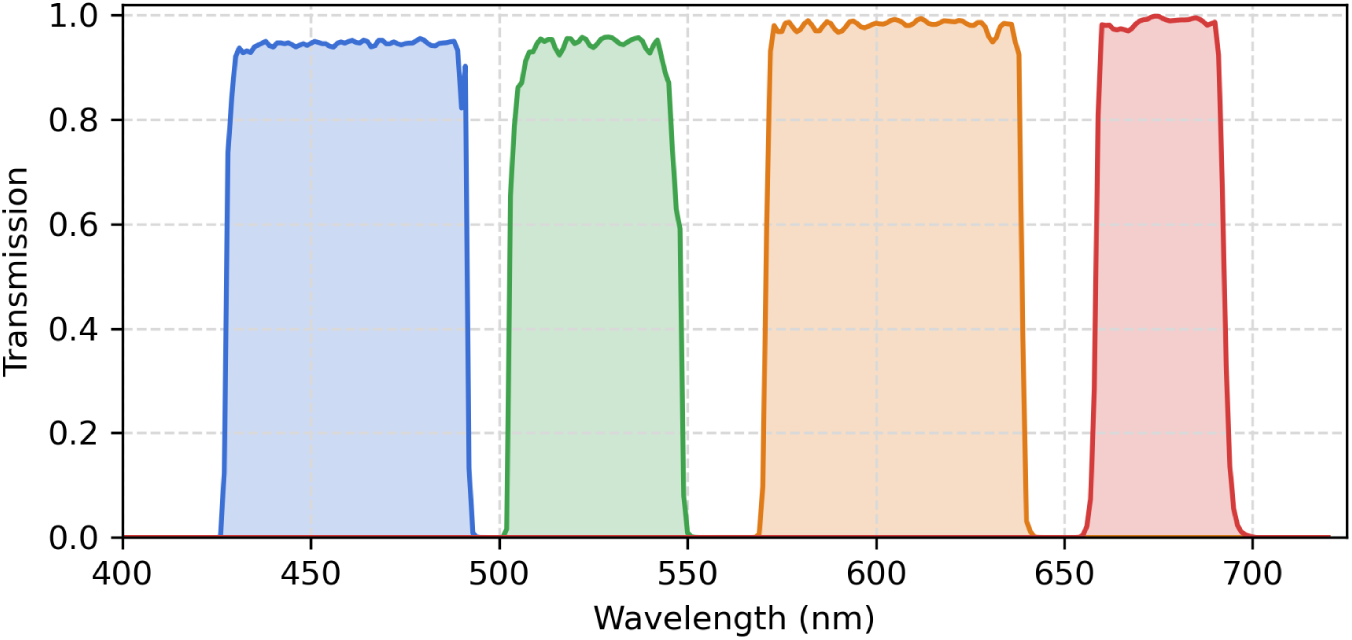
Filter pass bands used on a typical 4-channel BrainSaw system. Blue: Semrock FF01-460/60. Green: Semrock FF01-525/39. Red: Chroma ET605/70m. Far-red: Semrock FF01-676/29.

Silicon photomultipliers (SiPMs), such as Thorlabs PDA45, are an alternative to conventional PMTs. Whilst these detectors have a small 3 mm active area, they can be integrated into systems with large back aperture objectives (Modi *et al*. 2019) and have quantum efficiencies comparable to GaAsP detectors. SiPMs cannot be damaged by high light levels and may be permanently left switched on, making them well-suited to facility use. These detectors are especially cost-effective as they have built-in pre-amps (a feature that significantly reduces their sensitivity to electrical noise). In our tests, five of six Thorlabs PDA45 detectors had significant offset drift over the course of a typical acquisition. This is inconvenient but can be completely eliminated from the data using a moving baseline correction prior to stitching (see StitchIt commit 8676986).

#### Vibratome

We use the section head and knife holder from Leica’s VT1000S vibratome (P/N 14047248229 and 14046230131, respectively), sourced as spare parts from the manufacturer. The section head contains a Faulhaber DC motor which we drive with a Faulhaber MCDC3006 S RS controller. Setup instructions are available in the BakingTray documentation. The MCDC3006 S RS controller is, unfortunately, discontinued, with no direct replacement from the manufacturer; we recommend finding refurbished units on the secondary market, although BakingTray does support alternative controllers (Supplementary Table 2). We source high-quality stainless-steel blades from Campden Instruments (model 752-1-SS) and change them before each acquisition.

#### Water bath

Slicing and imaging take place in a water bath filled with approximately 750 mL of 50 mM phosphate buffer. The water bath is milled out of a single acrylic block with internal walls sufficiently polished to allow visualisation of the sample from the side during setup (Fig. 12C). The microscope slide is clamped to a stainless steel pedestal fixed to the bottom of the water bath (Fig. 12). The pedestal position is consistent across sessions, so the slide position is precisely known to BakingTray, simplifying sample setup. CAD models for the water bath are available at SWC-Advanced-Microscopy/BrainSaw_CAD on GitHub.

**Figure 12.**
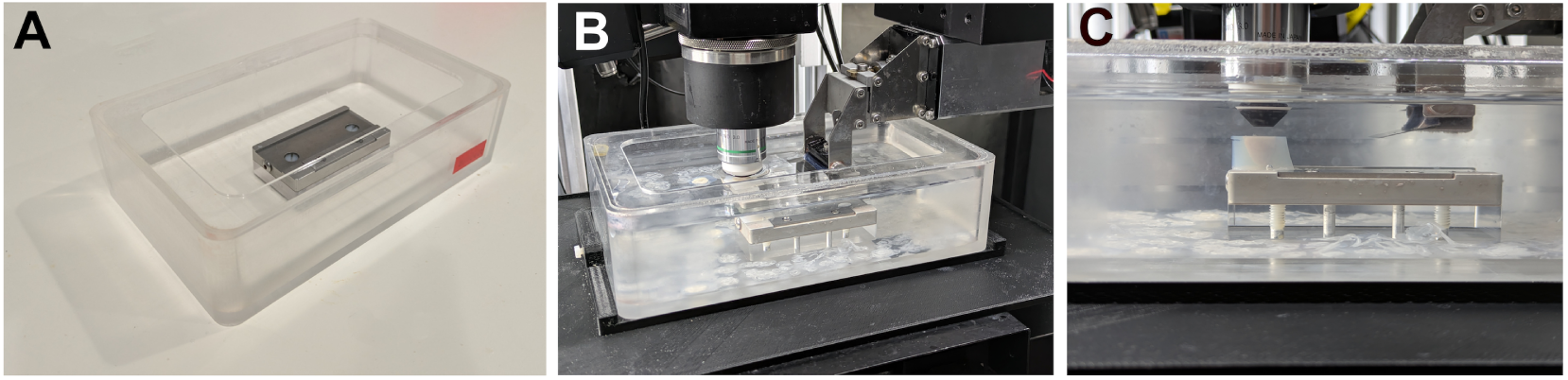
Water bath. **A**. Unfilled BrainSaw water bath. **B**. Water bath on a microscope containing a sample. **C**. Side view of the sample and blade through the wall of the water bath.

#### Motion control: X/Y tile-scanning

We recommend Physik Instrumente (PI) linear stages for X and Y motion. Many systems use V-551.4B and V-551.2B direct-drive stages, driven by C-891 controllers. We have also had good results with the legacy PI M-531.DD and M-605.2DD stages that came with TissueCyte 1000 systems. The stages used for tile scanning are a critical component and cheaper alternatives are not recommended given the workload these stages will accumulate (Fig. 2). BakingTray supports Zaber stages, but we have found the X-LRM series requires periodic lubrication and the lead screws wear over time, making them less suitable.

#### Motion control: Z lift stage

The X/Y stages are mounted on top of a Z ‘lift’ stage. The Z lift stage is responsible for moving the sample upwards during acquisition by the thickness of one section in order for it to be sliced. The easiest way to construct this X/Y/Z platform is to use a high-load lift stage capable of accurately pushing the tile-scanning stages and water bath. Few such stages are available on the market. We have successfully built systems with Aerotech PRO190SV-035 or Prior FB204E lift stages. The Prior stage coupled with their OptiScan III controller works well and is cost-effective. Around 2024/2025 Aerotech discontinued their Ensemble and Soloist motor controllers, replacing them with the Automation1 series and dropping MATLAB support. Aerotech reported that MATLAB support was reinstated for Automation1 in mid-2026 (personal communication), but we have yet to test this and there is currently no hardware control class for the new controller. Zaber’s VSR lift stages are another possibility and are currently supported by BakingTray, but they have not been tested in this application.

#### Motion control: objective coarse and fast focus

The objective is mounted on a ‘coarse’ focus motor, which is used to find the sample surface during setup. This process is controlled entirely via ScanImage and any ScanImage-supported hardware will work. We have had success with Zaber stages and controllers for this application. Fast Z-stacks during imaging are also controlled via ScanImage: we use PI’s PIFOC systems.

### 4.2 Software

#### Installation

BakingTray requires MATLAB (we use 2021b currently) and ScanImage Basic 2023 (earlier versions should work, but the latest 2023 versions have relevant bug fixes and useful features). ScanImage is a robust, well-tested product that runs on a range of image acquisition hardware and comes with technical support. Full installation instructions, including ScanImage configuration, BakingTray hardware class setup, and calibration procedures, are available at bakingtray.swcmicroscopy.com.

#### General operating principle

ScanImage and BakingTray communicate bidirectionally, with ScanImage handling image acquisition tasks and BakingTray handling the X/Y/Z stage, slicing, laser comms, and the setting of most acquisition parameters. The ScanImage source code is unmodified as BakingTray interfaces with it via API calls. Acquisition of a sample is dominated by tile-scanning epochs and slicing epochs. A ‘tile’ constitutes data from a single X/Y stage position comprising all optical planes and all acquired detector channels. In a three-channel acquisition with five optical planes per physical section, each tile is saved as a small TIFF stack containing 15 images. All tiles from one physical section are stored within a single sub-directory.

When an acquisition is started, BakingTray determines the number of tile positions and requests that ScanImage perform a ‘Loop’ over that many ‘Acquisitions’. Once each tile has been acquired, a BakingTray callback function (SIBT.tileAcqDone) is run by ScanImage. This initiates an X/Y stage move to the next tile position, updates the local tile-scan preview window, then triggers acquisition of the next tile. Once all tiles have been acquired, ScanImage returns to ‘Idle’ mode and BakingTray initiates a cutting cycle. The sample is moved upwards by the thickness of one section, driven into the blade at a pre-defined speed (typically ∼0.4 mm/s), then moved back under the objective. The tile-scanning and slicing process repeats until the requested number of sections has been obtained.

#### ‘autoROI’ automatic sample detection

Acquisitions may be of two types: ‘manual ROI’ or ‘autoROI’. A manual ROI acquisition is one where the user takes a tile-scan preview of the sample at the block face and draws a bounding box around the area they wish to image. This fixed ROI is applied to all sections. Manual ROI acquisitions of whole organs are inefficient since the area of most samples at the block face will change dramatically during the acquisition. They are particularly inefficient when a single acquisition includes multiple samples. To solve this problem, BakingTray has a threshold-based ‘autoROI’ feature to automatically detect the sample and calculate the ROIs as the acquisition proceeds.

Most acquisitions are conducted in autoROI mode. The user takes a preview tile-scan image of the sample at the block face as before. This image is downsampled to 50 *µ*m/pixel and median filtered (filter size: 5 by 5 pixels), and the pixels lying within 200 *µ*m of the image border (4 pixels) are taken as background. The tissue/background threshold is set to the median of this background distribution plus *k* standard deviations. If *k* is too high, some of the sample may be treated as background and not imaged. If *k* is too low, the resulting ROI tends to expand until it encompasses the entire agarose block. To choose *k*, the segmentation is run repeatedly for *k* decreasing from 12 to 2; *k* is set to the centre of the widest range over which the number of detected regions is invariant. This selection criterion is effective at finding tissue and *also* centres *k* in the region where the segmentation is least sensitive to the threshold. As a result, the procedure is tolerant of imperfect background estimates caused by tissue intruding into the border band.

The chosen *k* is applied to the median-filtered image to generate a binary mask. This is cleaned with morphological operations. Objects with high eccentricity are removed, as long thin structures are the signature of electrical noise in poor SNR images. The binary image is further median filtered (filter size: 5 by 5 pixels), and an erode–dilate cycle removes remaining speckle using a disk-shaped structuring element of 50 *µ*m radius, which at this sampling is a single pixel and so produces a 3-by-3 cross. The mask now represents real tissue, with rare exceptions. It is dilated by 600 *µ*m to provide a safety buffer. Minimum bounding boxes are computed around the surviving regions, boxes are merged where doing so increases the imaged area by less than 30%, and each box is finally rounded up to the nearest whole number of tiles for imaging.

After the user presses ‘Bake’, the acquisition begins and the tile pattern is imaged. Once each section is complete the sample location is recalculated and the block is sliced. On these subsequent sections the analysis is confined to the previously imaged ROIs: background statistics are drawn only from pixels near each ROI edge *that the previous section’s mask identified as non-tissue*. Thus, after the acquisition has begun, it is no longer assumed that all border pixels are background pixels. The absolute threshold is smoothed as a rolling median over the last five sections. If tissue is found abutting a ROI edge over a length greater than 350 *µ*m, that ROI is grown by 450 *µ*m in the corresponding direction, which allows the imaged area to track a sample that is growing or drifting fast at the block face. If the ratio of tissue to background area changes by more than tenfold between consecutive sections – as happens if the laser power or wavelength is altered mid-acquisition – the threshold is recalculated from scratch. The process repeats until the requested number of sections has been obtained or no further tissue is found.

To develop this process we collected a diverse set of 133 past acquisitions encompassing over 300 samples. This library included both typical samples and unusual ones (eyes, spinal cords, ferret hemispheres), as well as poor acquisitions with very low signal (e.g. laser power too low, imaging plane too deep, or incorrect wavelength). The location of the tissue in each section was manually annotated to obtain a ground truth and the autoROI algorithm was developed heuristically against this.

BakingTray’s autoROI has acquired about 6,000 samples at the SWC since 2020 with, to our knowledge, only one unusual incident where a sample was lost. In typical use, the autoROI loses no tissue or, in rare cases, 2–6 tiles’ worth of data out of the roughly 20,000 that make up a mouse brain. Overall, the autoROI has proved central to decreasing not only acquisition time but also sample setup time, as well as to streamlining user training.

### 4.3 Log analysis

We mined BakingTray log files on our storage platform to derive throughput and reliability statistics. BakingTray acquisition directories were identified by the presence of a ‘recipe’ file and a data folder. To simplify analysis we made a copy of the storage platform’s directory tree, retaining only the BrainSaw directories and the metadata files: the ‘recipe’ files and the ‘acquisition log files’. We analysed all available samples up to mid-July 2026.

#### Samples and acquisitions

The recipe file records information such as the start date and time, BakingTray imaging settings, key ScanImage settings, and the microscope name. The file is written at the start of the acquisition and a copy with a new time stamp is written upon restart of a stopped acquisition. Selected lines from one of these files are shown here:

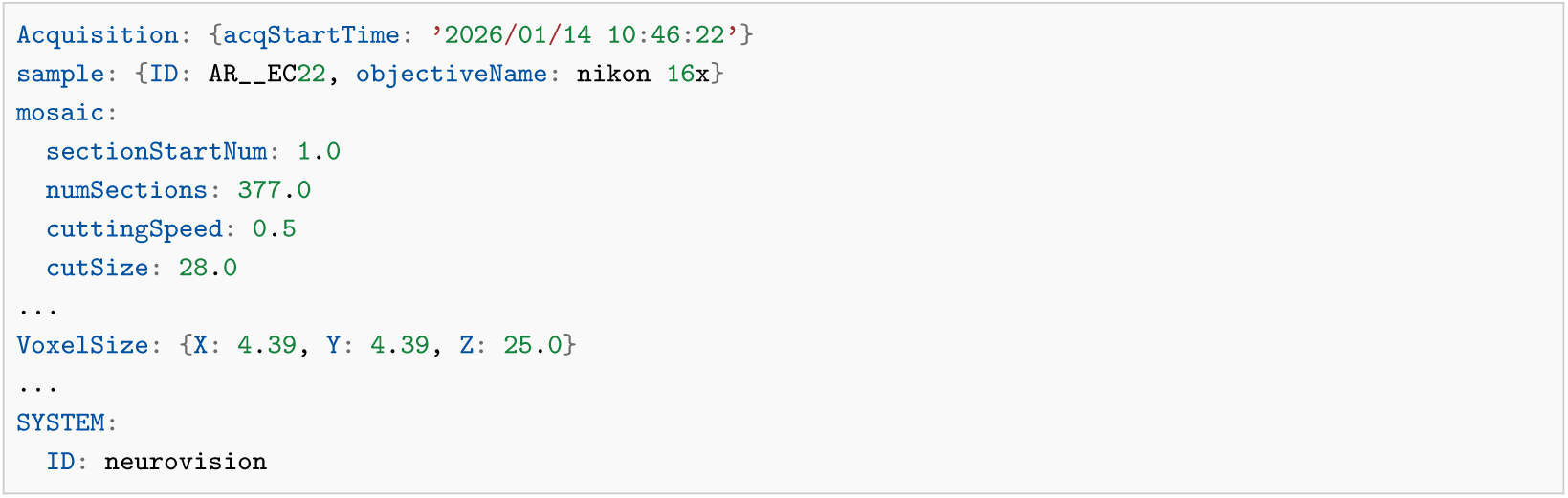

Individual acquisitions may contain multiple samples and we were interested in extracting statistics at both levels (samples and acquisitions). Sample directories were spread across the whole storage platform, spanning multiple research groups, with differing data management practices. As a result, it was challenging to ensure our final dataset did not contain duplicate acquisitions. A sample was defined as a directory containing a recipe file, and samples were counted on this basis. Because some samples were present in more than one path on the system, each was counted only once. Two recipe files were treated as duplicates if they had identical content and the same sample name. They were also treated as duplicates if they had the same microscope and sample name and belonged to the same acquisition. The sample name was taken from the file name. Backup copies of recipe files that BakingTray keeps in its logfiles directory were ignored. Where a sample directory held more than one recipe file, usually because a restarted acquisition writes a new date-stamped recipe file, we kept only the one whose start time was closest to the first start recorded in the directory’s acquisition log. For an acquisition resumed into a new directory, the samples of only one of the two runs were counted (see below). An acquisition that cut five or fewer sections in total was considered a test or a failed setup, and was counted neither as an acquisition nor as a sample.

StitchIt always produces one TIFF per optical plane per channel, so when several samples are imaged together they all appear in the same image. A software tool allows users to crop the individual samples into separate directories with unique sample names. Each holds a copy of the recipe file renamed for that sample and with the sample ID appropriately modified in the body text. These split samples do not carry an explicit record indicating with which samples they were acquired. In order to determine which samples were acquired together, and therefore the total number of acquisitions, we had to identify samples acquired at the same time on the same microscope. This was done based on the Acquisition.acqStartTime and SYSTEM.ID fields in the recipe file. Together, these data allowed us to plot the cumulative number of samples over time and to determine the distribution of the number of samples per acquisition. Similarly, the distribution of image resolutions across acquisitions could be extracted from the recipe files.

#### Restarts

The acquisition log file is a running log to which BakingTray appends progress during the acquisition. It begins with the acquisition start time, software versions, and laser wavelength and power, then records time-stamped progress messages. These include the laser state and enough information to recover the total acquisition time and how it was distributed across sections (directory paths are clipped below to reduce line length). The start and end of this message block are shown below. The final line indicates that the acquisition stopped at section 373 because no further tissue could be found. The number of requested sections appears to change from 377 to 368 because this acquisition was restarted after nine sections.

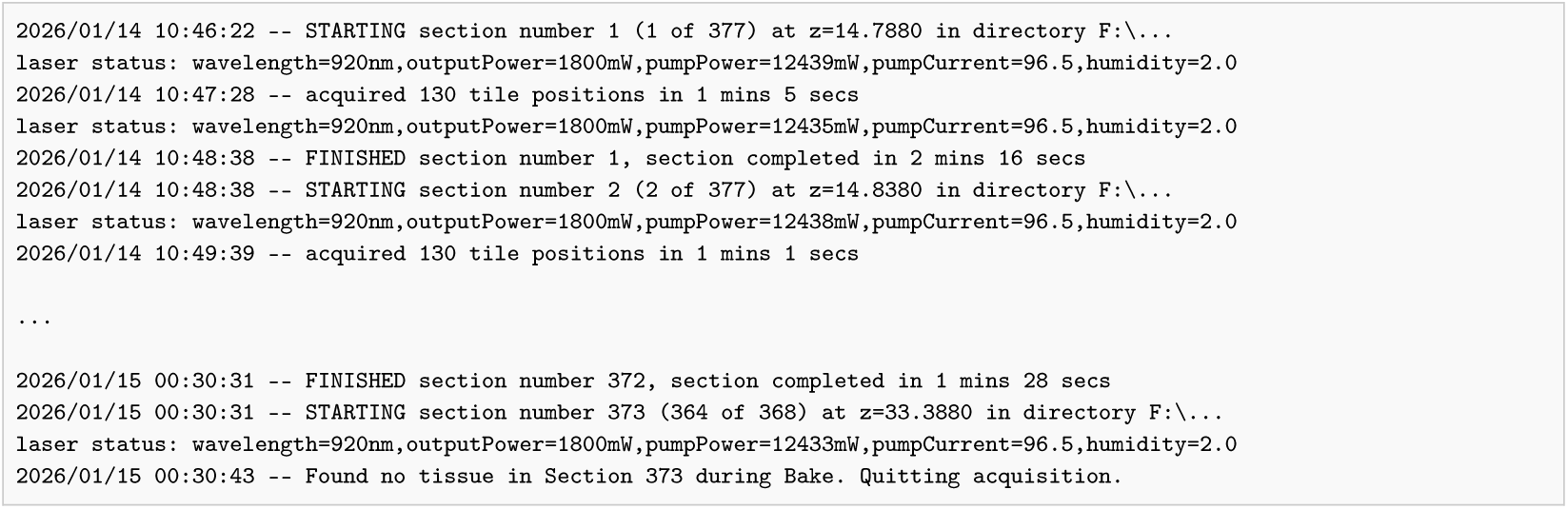

When a stopped acquisition is resumed, BakingTray appends a new header block, beginning STARTING NEW ACQUISITION, to the existing acquisition log. Every STARTING NEW ACQUISITION line after the first in a log was taken to be a restart, and each restart was classified from the log lines that preceded it, back to the previous start. Cases where the user manually aborted and restarted an acquisition are clear. In the following instance the user requested that the acquisition stop after section 13 and then restarted it within 5 minutes. This is indicated by the abortAfterSectionComplete message at 15:47:21.

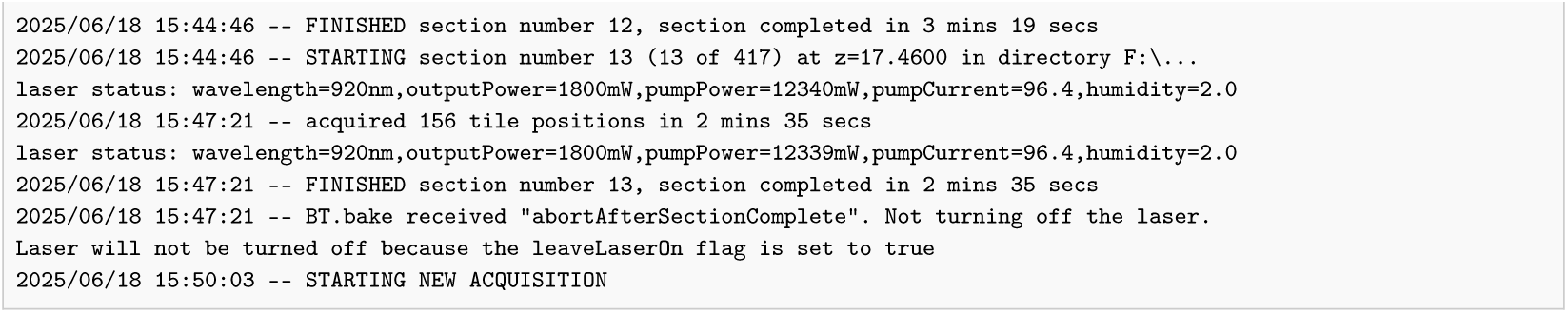

By contrast, the following log lacks the abort message but was restarted by the user about an hour later. This restart was classed as a stop for unknown reasons.

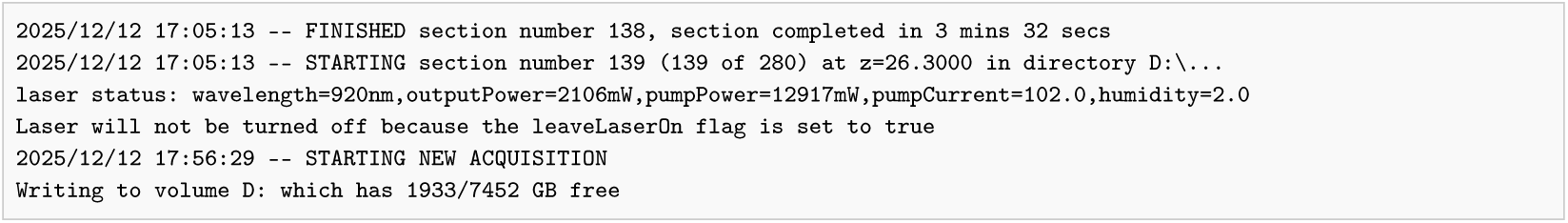

Restarts were placed in one of four categories based on the log file messages.

1. Manually aborted acquisitions: abortAfterSectionComplete
2. Acquisitions that stopped as planned and were then restarted by the user: FINISHED AND COMPLETED ACQUISITION, or Found no tissue … Quitting acquisition
3. Laser-related: STOPPING ACQUISITION DUE TO LASER or LASER NOT RUNNING
4. Stopped for unknown reasons

Note that the laser failures in most cases reflected a now-resolved bug in serial communications rather than a mode lock or power failure. Each restart was identified by its microscope and time, so it was counted once however many co-imaged samples’ logs recorded it; an acquisition restarted twice contributed two restarts.

Occasionally a stopped acquisition was resumed under a new name, so that the resumed run had its own directory, recipe file and acquisition log. The acquisition log records, for each section, the cutting depth and the directory on the acquisition PC to which the data were written. We treated two consecutive acquisitions on the same microscope as a single acquisition that had been restarted if the second began at the depth at which the first would have cut its next section. We also merged two such acquisitions if all three of the following were true. First, the later run began within 24 hours of the end of the earlier one. Second, it was written to a directory with the same name as the earlier run, or that name with a suffix (e.g. AB_rab0405 and AB_rab0405_NEWLASER). Third, it either continued the earlier run’s section numbering or followed an earlier run of fewer than 80 sections. In total, 22 pairs of runs were merged in this way (12 by depth, 10 by directory name). Both runs contain the same samples, so only one set was counted: that of the run split into more sample directories (an uncropped run holds all its samples in one directory). The stop between the two runs was treated as a restart, classified from the end of the first run’s log, and the imaging time of the two runs was summed.

A restart was not counted if five or fewer sections had been cut since the acquisition was started or last restarted, whatever the reason for the stop. Such stops occur during setup, for example when the user has requested the wrong number of sections (in some cases just one section) or notices a problem with the first sections. These rapid restarts reflect the correction of a setup error rather than the reliability of the system. Of the 311 restarts found, 100 were excluded on this basis, leaving 211. From these data we could calculate the number of restart events, their distribution over time, and their causes. In some cases the nature of the restart event was ambiguous. For example, some logs lacked an abortAfterSectionComplete message, yet the acquisition was restarted almost immediately, suggesting the user was present at the system the whole time. We classed these events as unplanned restarts, even though they may not have been. Thus, whilst the exact nature of some restarts remains unclear, every restart after more than five sections is included in the statistics.

#### Acquisition duration

The total duration of each acquisition was calculated from the log files. The time taken for each section is printed on each FINISHED line, and these times were summed. This approach excluded any downtime in acquisitions that were stopped and then restarted. The thickness of tissue cut varied by sample, as some users do not acquire the whole brain. We therefore divided each acquisition’s imaging time by the depth cut and multiplied by 12.5 mm, to report the time taken to image 12.5 mm of depth. Runs that cut less than 9.5 mm were dropped rather than scaled. For the graph showing acquisition time per brain we divided the total normalised time by the number of samples in the acquisition. To reduce variance we filtered the data in the following ways. We excluded rat brains based on the initials of users known to run these samples, read from both the sample directory name and the sample ID in the recipe file. We discovered the presence of uncropped multi-brain acquisitions in the dataset, based upon their sample directory names. We excluded these, and other such potential acquisitions, by removing all single-sample acquisitions whose mean number of tile positions per section was more than 1.6 times the median of that value across single-sample acquisitions at the same resolution. The median was about 58 tile positions per section at both resolutions, giving an exclusion threshold of about 92. These 40 excluded acquisitions include acquisitions of several brains that were never cropped into separate directories, often with a recipe sample ID naming several animals, as well as single brains imaged over an unusually large area. They remain counted as single samples elsewhere, as the number of brains they contained is not known. We restricted the analysis to the two microscopes with 8 kHz resonant scanners; the third microscope in the Facility has a 4 kHz scanner and so ran more slowly. We used data from 2020 onwards, once both microscopes had become well-established in the Facility. We make available the data and code used for the above analyses at github.com/SWC-Advanced-Microscopy/BrainSaw_Log_Analysis.

### 4.4 Imaging and sample processing methods

This section describes the methods used to process samples presented in this manuscript.

#### Surgical procedures and perfusions

Surgical procedures for virus injections were carried out under isoflurane (2–5%) with oxygen as carrier gas at a flow rate of 2 L/min and after carprofen (5 mg/kg, s.c.) had been administered. For all surgical procedures eye gel was applied (Lubrithal) and body temperature was controlled via a rectal probe and an electrical heating pad set to ∼35^◦^C. All surgeries were performed on a stereotaxic frame (Benchmark). For perfusions, mice were deeply anaesthetised using sodium pentobarbital (80 mg/kg, i.p.). A blunt needle was placed in the left ventricle and an incision made in the right atrium of the heart. Using handheld 10 mL syringes, blood was cleared with 100 mM PBS (10 mL, 2.5 mL/min) and the animal was then perfused with 10 mL PBS containing 4% PFA. After successful fixation assessed via muscle rigidity, the head was removed and the brain carefully extracted. The brain was further fixed by immersion in 4% PFA overnight at 4^◦^C, and then stored in PBS at 4^◦^C until ready for imaging.

#### Sample preparation for imaging

Detailed sample preparation protocols are maintained in the BakingTray user guide. Samples were embedded in 5% agarose made in 50 mM phosphate buffer (Campbell 2024a), which served as a pH buffer during imaging. Samples were embedded either singly (Campbell 2024c) or in groups of four (Campbell 2024b) using custom embedding moulds. The agarose blocks were removed from the moulds and super-glued to the frosted portion of a conventional glass microscope slide.

#### Spleen

The spleen sample in Fig. 4C is from a Tlr7-GoF Gcsam-ert2cre Confetti (Het) mouse treated with a single dose of tamoxifen (15 mg) seven days prior to tissue harvesting. This mouse line generates an autoimmune phenotype causing the spleen to produce spontaneous germinal centres. Germinal centre B cell confetti-expression was induced by Cre expression from the Gcsam promoter (specific to germinal centre B cells). The sample was imaged with two laser lines simultaneously: a Spectra-Physics Mai Tai eHP DS (930 nm, 97 mW at the sample) and Coherent Axon 1064 (1064 nm, 141 mW at the sample); the image resolution was 2 × 2 × 10 *µ*m/voxel.

#### Whole-mount labelling

Tyrosine hydroxylase (TH) was labelled after tetrahydrofuran delipidation (EZ Clear; Hsu *et al*. 2022) with a recombinant monoclonal antibody conjugated to Vio667 (Miltenyi Biotec). The brain was imaged with 800 nm excitation, 125 mW at the sample, collecting the far-red and green channels. The protocol is described in detail in Appendix 6. Congo red labelling of amyloid plaques was performed on half-brains delipidated with CUBIC-L (Tainaka *et al*. 2018), stained with Congo red in ScaleCUBIC-1a (Susaki *et al*. 2014), and then strengthened with gelatin cross-linked by PFA to counteract the softening caused by delipidation (Appendix 7). Brains were imaged with 820 nm excitation, 80 mW at the sample, collecting the blue and red channels. We initially tried an iDISCO-based approach (Liebmann *et al*. 2016), but this resulted in a bright Congo red rim on the sample surface.

#### Dual-colour retrograde viral tracing

For dual-colour retrograde viral tracing, we used rAAV2-retro-EF1a-H2B-eGFP and rAAV2-retro-EF1a-H2B-tdTomato (1 × 10^13^ GC/mL) (Tervo *et al*. 2016; Weiler, Teichert, and Margrie 2025). After shaving the dorsal surface of the head with clippers and performing an antero-posterior incision through the midline of the head using a scalpel, a small craniotomy was performed using a handheld dental drill (Osada). Virus injections were then performed using borosilicate glass injection pipettes (Wiretrol II; Drummond Scientific) pulled to a taper length of ∼30 mm and a tip diameter of ∼50 *µ*m using a vertical puller (PC-10, Narishige). Virus volumes were 60 nL each and viruses were delivered at a rate of 1–2 nL/s using Nanoject III (Drummond Scientific) and injected at a cortical depth of 650 *µ*m. The injection coordinates relative to bregma were 2.1 mm lateral for retro-GFP, 2.25 mm lateral for retro-tdTom, and 3.8 mm posterior for both viruses. After injection, the craniotomy was sealed with silicone (Kwik-Cast), the skin was re-sealed using tissue adhesive (3M Vetbond), and animals were left for 2–3 weeks to allow for expression before being perfused. We imaged brains on a BrainSaw system equipped with a Coherent Axon 1064 co-aligned with the main tunable Mai Tai laser. For these specific experiments the Mai Tai ran at 920 nm and 75 mW, with the Axon 1064 nm laser at 150 mW.

#### Mapping synaptic sites

To map putative synaptic sites across the brain, we injected 60 nL of AAV[2/1]-hSyn-DIO-tdTomato-Synaptophysin-eGFP (1.90 × 10^13^ GC/mL) into the vLGN. Four Cre driver lines were used, all from the SWC Mouse Colony: vgat-Cre (STOCK *Slc32a1*^tm2(cre)Lowl^/J), PV-Cre (B6;129P2-*Pvalb*^tm1(cre)Arbr^/J), Penk-Cre (B6;129S-*Penk*^tm2(cre)Hze^/J), and SOM-Cre (STOCK *Sst*^tm2.1(cre)Zjh^/J). Head shaving, midline incision, craniotomy and post-injection wound closure were as described in Section 4.4. Injections were made using glass pipettes (4.2 *µ*L capillaries with ID 0.53 mm, Drummond Scientific) pulled on a P-1000 micropipette puller (Sutter Instrument) and sharpened using a micro-grinder (EG-45, Narishige). Viral injections were performed at 1–2 nL/s using a Nanoject III (Drummond Scientific). The injection coordinates relative to bregma were −2.35 mm anterior-posterior, 2.43 mm medial-lateral and −3.44 mm dorsal-ventral. Animals were left for 7 weeks to allow for expression before being perfused. Brains (n=1 from each line) were imaged using STPT at 2 × 2 × 5 *µ*m/voxel, with 40 *µ*m sections, eight optical planes, and four-frame averaging. Some initial pilot brains were imaged at higher resolutions (e.g. see Figure 8–figure supplement 1). We used 920 nm excitation at about 130 mW at the sample.

#### STPT imaging procedures

All imaging data presented in this manuscript were obtained from one of the three BrainSaws at the SWC Advanced Microscopy Facility (AMF). Systems were equipped with PIFOC fast focusing systems (either 100 *µ*m or 400 *µ*m travel range). Images were acquired with a Nikon 16x/0.8 water-immersion objective. Systems were equipped with a 4 kHz or 8 kHz resonant scanner from Cambridge Technology. We used Hamamatsu R10699 PMTs selected for higher gains. Signals were pre-amplified using Femto DHPCA-100 (switched to 80 MHz bandwidth) or Thorlabs TIA60 transimpedance amplifiers. Data acquisition was controlled using NI DAQ hardware. All rigs had Spectra-Physics Mai Tai eHP lasers with DeepSee. Beam intensity was modulated using a Conoptics Pockels cell and ScanImage was calibrated to report power at the sample across all wavelengths employed in these experiments. We typically image at around 120 mW at the sample but the range of power values used spans roughly 75–175 mW. The first optical plane was typically taken 25–35 *µ*m below the sample surface.

### 4.5 Data processing during acquisition and stitching

We acquire data to a dedicated 8 TB SSD on the Windows acquisition PC. Each acquisition has its own folder. The folder contains metadata associated with the acquisition parameters, log files, and a rawData folder where images are stored. Each physical section has its own sub-folder within rawData. Each X/Y tile position is saved as a small TIFF stack comprising all channels and optical planes acquired at that position.

We have a separate Linux PC running our ‘StitchIt’ analysis package to service all three acquisition systems. Once the acquisition has begun, the StitchIt command ‘syncAndCrunch’ is initiated. This automatically searches for a currently running acquisition and then repeatedly runs rsync to copy data from the acquisition PC to a local RAID array on the analysis PC. After each section is complete, the tiles are loaded, statistics are extracted, and average images from that section are saved. Grand averages of these images are periodically generated. These grand averages are used to correct for vignetting and other artefacts that introduce systematic inhomogeneity to the tiles. A preliminary image is stitched and sent to a web-server, providing users with a way of monitoring live acquisitions. At the end of the acquisition, a file called ‘FINISHED’ is written to the acquisition folder. The analysis PC detects this and initiates stitching of the whole dataset. Stitched data are saved as one 16-bit TIFF per optical plane and channel. StitchIt also generates downsampled stacks with isotropic voxels (e.g. 10 or 25 *µ*m), saved as multi-page 16-bit TIFFs, which are convenient for visualisation and for re-slicing in other planes.

Our stitching pipeline is fast and works purely from the recorded stage positions. Stitching errors are typically under 30 *µ*m, which is adequate for our use cases. More elaborate but more accurate stitching approaches have been created by others for STPT (see Economo *et al*. 2016; Chen *et al*. 2020, and in particular González-Solares *et al*. 2023).

### 4.6 Post-acquisition data processing

After stitching is complete, users interactively run a command called stitchit.sampleSplitter to crop samples (for a single-sample acquisition) or to split the acquisition into separate directories (for a multi-sample acquisition). For larger datasets, considerable space can be saved by this cropping operation.

A pip-installable Python package called btpytools is used to simplify file handling. This provides a shell command called compressRawData that initiates parallel bzip compression (pbzip2) of the rawData folder for users who wish to archive the raw tiles rather than relying solely on the stitched output. It also provides a transferToServer command that is an rsync wrapper for users to copy acquisitions from the analysis PC to a mounted storage server. This provides a simpler interface than rsync, double-checks that the transfer was successful, and writes an empty file to the acquisition directory indicating that the transfer succeeded. Successfully transferred folders are deleted automatically from the analysis PC via a cron job based upon the presence of this file.

#### Dual-colour automatic registration and cell detection

Images were registered to the Allen Mouse Brain Common Coordinate Framework (Wang *et al*. 2020) (10 *µ*m resolution) using the software tool brainreg (Tyson *et al*. 2022; Tyson and Margrie 2022), which is based on aMAP (Niedworok *et al*. 2016). All atlas data were provided by the BrainGlobe Atlas API (Claudi *et al*. 2020). Automated cell detection and deep-learning-based cell classification were performed by the cellfinder software (Tyson *et al*. 2021). Dual-colour cell detection (Results Section 2.7) was achieved by running cellfinder twice, once with the red channel as signal and green as background, and once with the channels reversed.

### 4.7 Animal welfare

All animal procedures were conducted in compliance with the guidance and regulations set forth by the UK Home Office according to the Animals (Scientific Procedures) Act 1986 and approved by the Animal Welfare and Ethical Review Body (AWERB; SWC) and in compliance with ARRIVE guidelines. PPL numbers associated with experiments that appear in this study and not elsewhere are:

- Dual-colour cell counting: PP1819274
- TH labelling, synaptic labelling, and bulk projection tracing: PP4459082
- Congo red labelling: PP2015516

The mouse spleen was supplied by the Francis Crick Institute in compliance with the aforementioned Act under PPL number PP2867252. Data from bird brains were obtained under animal procedures approved by the German local and national authorities for the use of animals in research (Niedersächsisches Landesamt für Verbraucherschutz und Lebensmittelsicherheit/LAVES, Oldenburg, Germany, Az.: 33.19-42502-04-20/3492; 33.19-42502-04-21/3650). All remaining data are taken from previous publications with associated animal use statements as cited.

### 4.8 Analysis

Whole brain data were visualised in napari (napari contributors 2019), where scale bars were added. Images were exported to Affinity Designer for sub-panel layout by taking a screenshot. The original scale bar was converted to a vector element before being cropped out. Line graphs were generated using Python.

#### Use of AI

BakingTray, StitchIt, and btpytools were written without the use of AI. Since Q2 2026, Claude Opus 5.0–5.5 and Claude Fable 5 have been used for code review of BakingTray and StitchIt. Proposed changes were usually applied manually based on a work-package provided by Claude and sanity checked before deployment.

This manuscript was written by the human authors. Claude was used to check it for errors and flow. In addition, Claude drafted Supplementary Table 1 and generated Figure 1–figure supplement 1 based on the Introduction. The autoROI algorithm is long and was written in 2020. Therefore, Claude used the source code to generate a prose summary, which was confirmed and manually converted into Methods Section 4.2. The analyses of the BakingTray log files were written in Python by Claude and checked thoroughly by a human. All AI-associated contributions were verified, edited, and approved by the authors, who take full responsibility for the content of this manuscript. The ‘channel chooser’ tool at swcmicroscopy.com/brainsaw_channel_chooser was written entirely using Claude Opus 5 and rigorously checked by a human. Source code is available at github.com/SWC-Advanced-Microscopy/brainsaw_channel_chooser.

## 5 Author Contributions

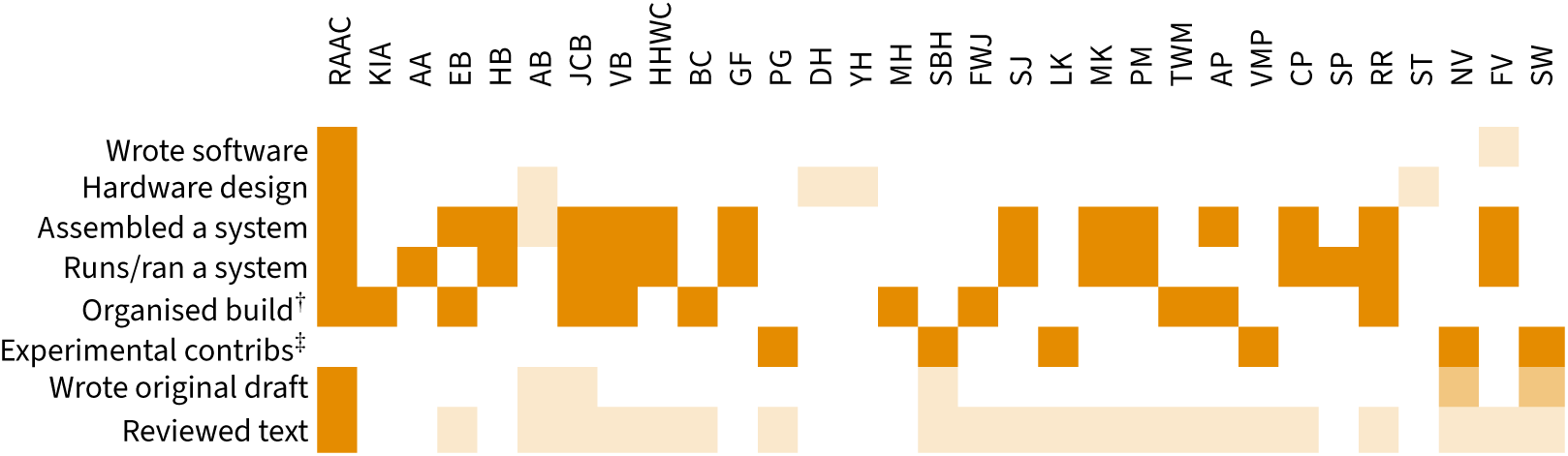

^†^**Organised a build** These authors were involved in roles such as funding acquisition, project leadership, or logistics of a BrainSaw install. Emmanuel Beaurepaire and Troy Margrie purchased TissueCyte 1000 systems that were later converted to BrainSaws.

^‡^**Experimental contributions** Four figures and associated data were generated by specific authors. The two whole-mount labelling protocols associated with Fig. 5 were designed by Peter Gordon. Peter Gordon performed the TH antibody labelling and acquired the data. Loukia Katsouri generated and imaged the Congo-red-labelled brains from the Alzheimer’s model and its control as part of a larger study currently under preparation. The rat cell counting and atlas registration experiment shown in Fig. 6–figure supplement 1 was the work of Viktor Plattner, who also made the figure and analysed the data. The cell counting experiment shown in Fig. 7 was the work of Simon Weiler, who also made the figure, analysed the data and wrote the associated Results text. The vLGN synaptic mapping experiment shown in Fig. 8 was the work of Nicole Vissers and Sonja Hofer, who also made the figure, analysed the data and wrote the associated Results text.

## 7 Acknowledgements

This project was conceived and initiated in the lab of Tom Mrsic-Flogel, whom we thank for providing the time, space, and funding necessary to build the first BrainSaw.

Funding for the BrainSaw system at the EPFL was obtained by Carl Petersen. Dan Gunton, now at Cairn Research, helped set up the CrickSaw system at the Francis Crick Institute when he was a member of the Institute’s imaging core facility. The BrainSaw team in Leuven thank Shu-Hao Yeh for early testing and Norma Kühn for microscopy help. Claudio Polisseni at the Max Planck Institute, Munich, was assisted in converting a Sutter MOM microscope to BrainSaw by Maximiliane Heckmann. Rajeev Rajendran at Netherlands Institute for Neuroscience was assisted in setting up BrainSaw by Roeland Lokhorst. The Ecole Polytechnique group thanks Jean Livet, Lamiae Abdeladim, Katie Matho, Gabriel Kaddour, Clément Caporal, and Anatole Chessel for discussions on implementation and benchmarking of multicolour STPT imaging.

We thank Mitchell Sandoe, Georg Jaindl, and Boris Djiguemde on the ScanImage team at MBF for assistance with ScanImage. Help with machining early in the project was provided by the Central Mechanical Workshop at the Biozentrum, University of Basel.

Numerous people, especially at the SWC, have made suggestions for new BakingTray features, and supported the project in a variety of ways. Igor Tatarnikov and Isabell Whiteley at the SWC AMF have helped troubleshoot the equipment and assisted users. User training was also provided by Patty Blakely, Ryan Cini, and Panagiota Iordanidou. Jessica Broni-Tabi has helped users to *post hoc* antibody label slices collected from the AMF’s BrainSaw systems. Adam Tyson encouraged development of multi-sample acquisitions and promoted BrainSaw via the BrainGlobe analysis tools. The ‘Auto-Trim’ feature was the suggestion of Stephen Lenzi. Alex Fratzl provided a lot of encouragement early in the project, including insisting on frame averaging, which required software changes to ScanImage. The name ‘BrainSaw’ was the idea of Petr Znamenskiy.

Piotr Nowak (SWC Viral Core) generated the viruses used in this study. Christophe Galichet and Luke O’Hara perfused some of the animals from which organs were harvested for this study. The spleen sample was provided by Amalie Grenov. Existing datasets used here as examples were provided by Rosa Holder, Viktor Plattner, Alex Fratzl, and Chaofei Bao. Harry Carey helped survey the brain registration literature and provided valuable feedback on the manuscript. Olivier Winter helped us generate the IBL probe tracking figure. We thank Adam Tyson and Isabell Whiteley for their valuable feedback on the manuscript. We thank Michelle Ashraf, who designed the logos associated with this project and made Fig. 1.

## 8 Data Availability Statement

This project is developed entirely in the open and all code is available. BakingTray and StitchIt are released under the LGPL. btpytools is released under the GPL. The CAD models are released under the MIT licence. We will maintain current links and information at swcmicroscopy.com/brainsaw/.

The Gad2 tdTomato transgenic mouse brain data shown in Figure 4–figure supplement 2 are available for download at https://doi.org/10.5281/zenodo.22114627. The Sst tdTomato transgenic mouse brain data shown in Figure 4–figure supplement 1 are available for download at https://doi.org/10.5281/zenodo.22232268. The bulk projection data shown in Figure 6E are available for download at https://doi.org/10.5281/zenodo.22304676. The whole rat brain data shown in Figure 4A are available for download at https://doi.org/10.5281/zenodo.22304842. The whole spleen data shown in Figure 4C are available for download at https://doi.org/10.5281/zenodo.22302483. The raw data and analysis scripts for the acquisition log statistics are at github.com/SWC-Advanced-Microscopy/BrainSaw_Log_Analysis. Other datasets are available on request.

## 9 Supplementary figures and tables

**Supplementary Table 1.**
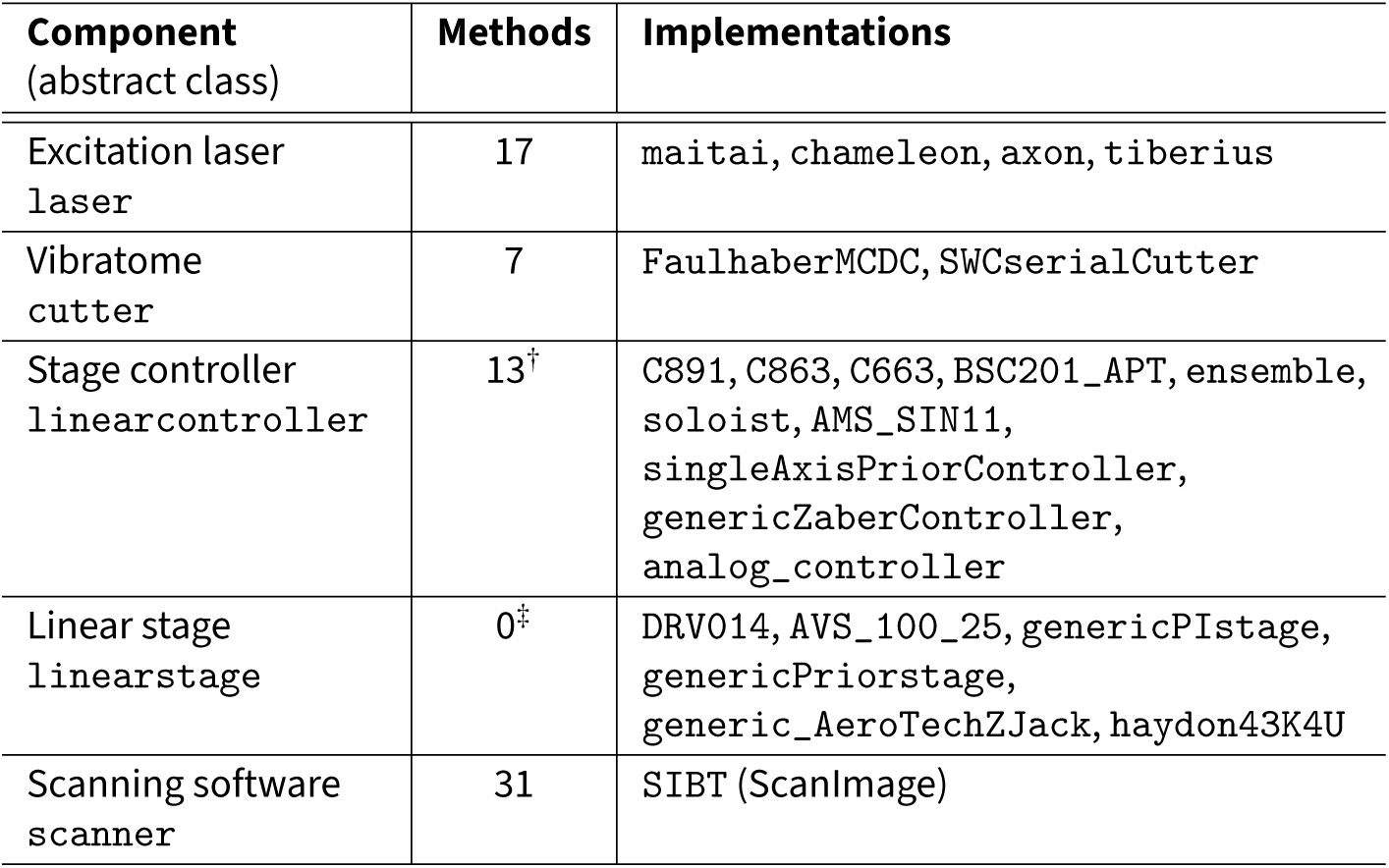
Component interfaces. . BakingTray’s core acquisition class, BT, contains no hardware-specific commands. Each category of hardware is instead represented by an abstract MATLAB class that declares a documented set of methods, and support for a new device is added by writing a single class that inherits it. “Methods” is the approximate number of methods a new class must define; the abstract class carries the documentation describing how each should behave. Each component directory contains a function that builds the requested class at BakingTray start-up, so the class is chosen by name in the componentSettings.m settings file. Simulated (“dummy”) implementations of every component are also provided, allowing the software to be run with no hardware attached. ^†^A further six methods relating to acceleration, initial velocity, and axis enabling are optional. ^‡^A linearstage declares no methods: it defines the physical properties of an axis, such as its travel limits and units, and is attached to a linearcontroller.

**Figure 1–figure supplement 1.**
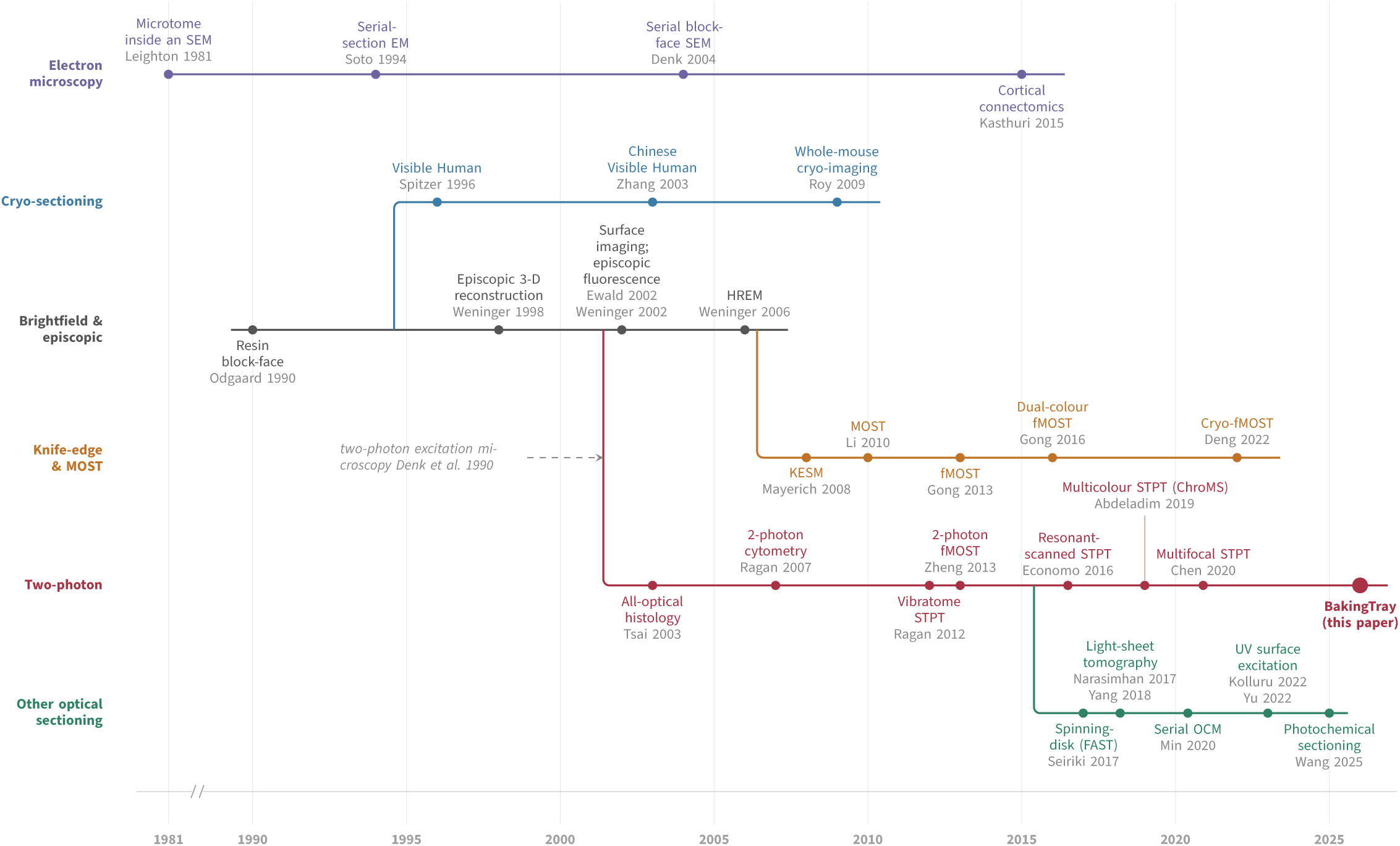
A timeline of block-face serial-section imaging. Each strand is a family of techniques; where one grew out of another it is drawn budding from its parent. Horizontal position is publication year, with a break in the axis before 1990. The strands differ chiefly in the imaging modality used. The figure is illustrative rather than exhaustive, and some instruments could reasonably be placed on more than one strand.

**Supplementary Table 2.** Hardware compatibility. . This table summarises the hardware compatibility of BakingTray. Where relevant, **bold** text indicates recommended options. The system works with either NI DAQ or vDAQ acquisition systems via ScanImage. We commonly use Nikon W 16x/0.8 objectives, but other objectives are also possible. ^†^There is currently no BakingTray support for Aerotech’s new Automation1 controllers (see Methods Section 4.1).

| Hardware class | Supported hardware |
| --- | --- |
| Lasers | • Coherent Chameleon and Discovery<br>• Coherent Axon fibre lasers<br>• Spectra-Physics Mai Tai eHP DS<br>• Thorlabs Tiberius (discontinued) |
| Vibratome | • Leica VT1000S section head (P/N 14047248229)<br>• Leica VT1000S knife holder (P/N 14046230131) |
| Vibratome controllers | • **Faulhaber MCDC3006 S RS**<br>• Custom SWC DC motor controller |
| X/Y Sample stage controllers | • **PI V-551.2B/.4B & 2x C-891**<br>• Zaber X-LRM (not recommended)<br>• PI M-531.DD (discontinued)<br>• PI M-605.2DD (discontinued) |
| Sample Z actuator & controller | • **Prior FB204E and OptiScan III ES11**<br>• Aerotech PRO190SV-035 and legacy Soloist/Ensemble controllers<sup>†</sup><br>• Zaber VSR40A-T3A (supported but untested) |
| Scanners | • **4 kHz and 8 kHz resonant scanners**<br>• Linear scanners |

**Supplementary Table 3.**
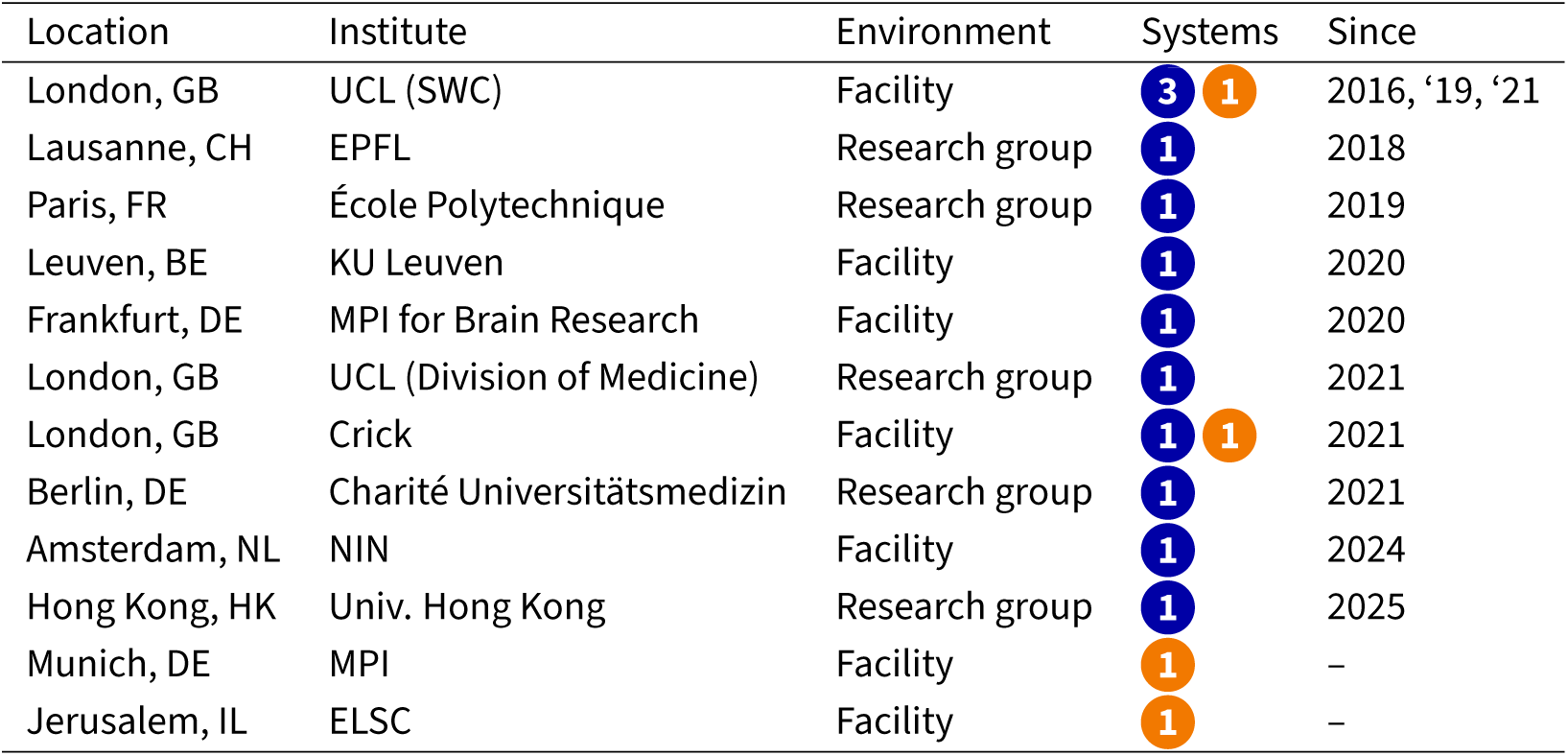
List of BrainSaw systems. . Circles indicate the number of systems at each site by status: 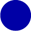 built; 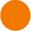 under construction. In total 12 systems have been built, the first in 2016. Years indicate when each system first became operational, and sites are ordered by that year. The system at EPFL is no longer in operation, but produced several publications; the system at KU Leuven was constructed but is currently being moved. The table is correct as of 1st September 2026.

**Supplementary Table 4.** List of the 35 publications that have used BakingTray.

| Study | Journal | Use case |
| --- | --- | --- |
| Thompson *et al.* 2026 | Nature Neuro. | Probe tracks |
| Sirmpilatze *et al.* 2026 | Curr. Biol. | Atlas generation |
| Lo *et al.* 2025 | Neuron | Bulk projections |
| Chen, Schmitz, and Tukker 2025 | Neuroscience Bulletin | Cell counting |
| Ahmadlou *et al.* 2025 | Nature | Probe tracks |
| Banga *et al.* 2025 | eLife | Probe tracks |
| Beau *et al.* 2025 | Cell | Probe tracks |
| Becalick *et al.* 2025 | bioRxiv | Cell counting |
| Edwards *et al.* 2025 | eLife | Cell counting |
| Greenstreet *et al.* 2025 | Nature | Fibre tracks |
| Int. Brain Lab. *et al.* 2025 | Nature | Probe tracks |
| Mederos *et al.* 2025 | Science | Probe tracks, fibre tracks |
| Vélez-Fort *et al.* 2025 | Cell | Probe tracks |
| Weiler, Teichert, and Margrie 2025 | eLife | Cell counting |
| Weiler *et al.* 2024 | Nature Comms. | Cell counting |
| Blanc *et al.* 2023 | ACS Photonics | Methods development |
| Ho *et al.* 2023 | Cell Reports Methods | Lesion mapping |
| Bimbard *et al.* 2023 | Nature Neuro. | Probe tracks |
| Campagner *et al.* 2023 | Nature | Cell counting |
| Chen *et al.* 2023 | Nature Comms. | Screening AAV expression |
| Tran-Van-Minh, Ye, and Rancz 2023 | PLOS ONE | Cell counting |
| Esmaeili *et al.* 2022 | PLOS Bio. | Bulk projections |
| Galloni, Ye, and Rancz 2022 | J. Neurosci. | Cell counting |
| Lai *et al.* 2022 | Nature Meth. | Whole-mount Ab. staining validation |
| Liu *et al.* 2022 | Front. Neuroanat. | Whole-brain single-axon tracing |
| Mazuski and O'Keefe 2022 | Cell Reports | Probe tracks |
| Tyson *et al.* 2022 | Sci. Rep. | Probe tracks |
| Keshavarzi *et al.* 2022 | Neuron | Probe tracks |
| Blot *et al.* 2021 | Neuron | Cell counting |
| Esmaeili *et al.* 2021 | Neuron | Probe tracks |
| Fratzl *et al.* 2021 | Neuron | Probe tracks, bulk projections |
| Orsolic *et al.* 2021 | Neuron | Mapping widefield data to Allen CCF |
| Tyson *et al.* 2021 | PLOS Comp. Bio. | Cell counting |
| El-Boustani *et al.* 2020 | Nature Comms. | Bulk projections |
| Mayrhofer *et al.* 2019 | Neuron | Bulk projections |

**Supplementary Table 5.**
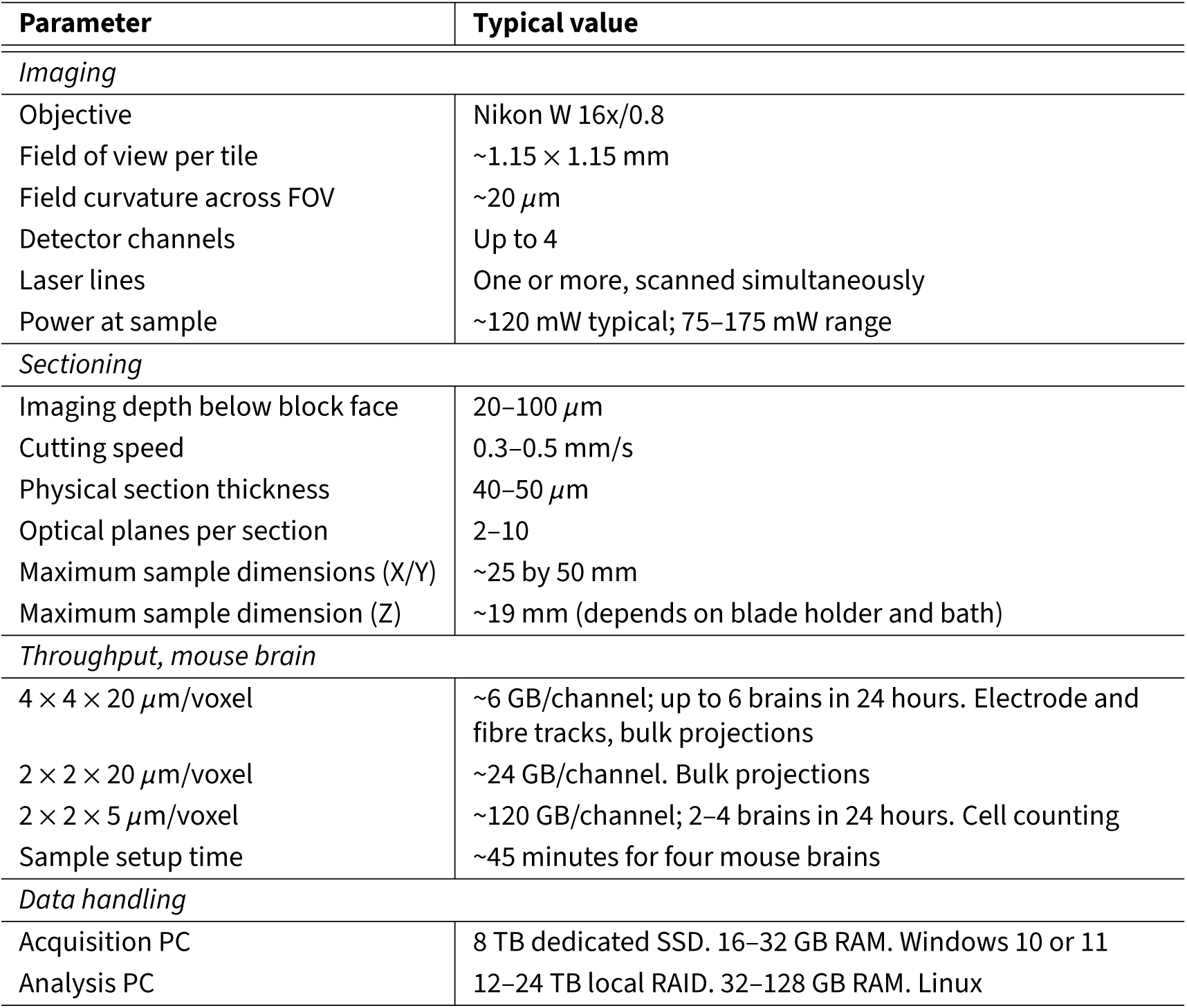
Typical acquisition parameters for a BrainSaw STPT system. . Values are those routinely used on the three systems at the SWC and are intended to give a prospective builder a sense of what the platform delivers in practice. They are not hard limits: resolution, section thickness, and sample number are all set by the user, and the achievable field of view depends on the objective and scan optics chosen. Sample size depends on water bath size, stage range, and blade holder length. Data volumes are quoted per channel per mouse brain; a four-channel acquisition of four brains scales linearly.

| Parameter | Typical value |
| --- | --- |
| *Imaging* |  |
| Objective | Nikon W 16x/0.8 |
| Field of view per tile | $\sim 1.15 \times 1.15$ mm |
| Field curvature across FOV | $\sim 20$ μm |
| Detector channels | Up to 4 |
| Laser lines | One or more, scanned simultaneously |
| Power at sample | $\sim 120$ mW typical; 75–175 mW range |
| *Sectioning* |  |
| Imaging depth below block face | 20–100 μm |
| Cutting speed | 0.3–0.5 mm/s |
| Physical section thickness | 40–50 μm |
| Optical planes per section | 2–10 |
| Maximum sample dimensions (X/Y) | $\sim 25$ by 50 mm |
| Maximum sample dimension (Z) | $\sim 19$ mm (depends on blade holder and bath) |
| *Throughput, mouse brain* |  |
| $4 \times 4 \times 20$ μm/voxel | $\sim 6$ GB/channel; up to 6 brains in 24 hours. Electrode and fibre tracks, bulk projections |
| $2 \times 2 \times 20$ μm/voxel | $\sim 24$ GB/channel. Bulk projections |
| $2 \times 2 \times 5$ μm/voxel | $\sim 120$ GB/channel; 2–4 brains in 24 hours. Cell counting |
| Sample setup time | $\sim 45$ minutes for four mouse brains |
| *Data handling* |  |
| Acquisition PC | 8 TB dedicated SSD. 16–32 GB RAM. Windows 10 or 11 |
| Analysis PC | 12–24 TB local RAID. 32–128 GB RAM. Linux |

**Figure 2–figure supplement 1.**
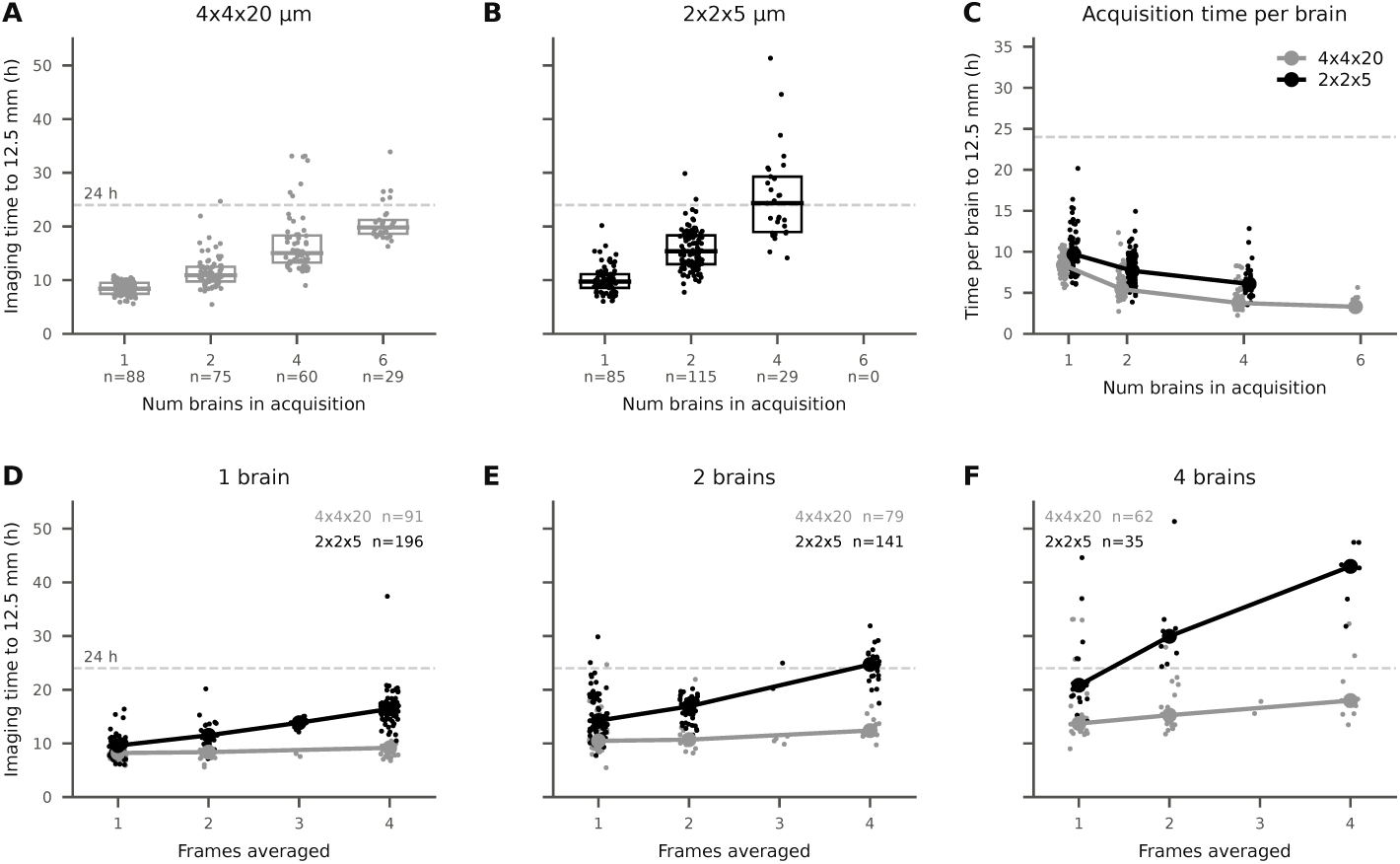
Acquisition time statistics. Data come from mouse brain acquisitions performed from January 2020 onwards from the two rigs at the SWC running 8 kHz resonant scanners. We did not include acquisition time data from our third microscope, which is equipped with a slower 4 kHz resonant scanner. **A&B**. Total acquisition time for a mouse brain normalised to 12.5 mm of imaged depth, plotted against the number of brains in the acquisition, for the two most common voxel configurations: 4 × 4 × 20 *µ*m (A, grey) and 2 × 2 × 5 *µ*m (B, black). **C**. Data from A and B normalised by the number of brains in the acquisition. Points are individual acquisitions, lines join the medians. Acquisition time per brain falls as more brains are added, mostly due to lower cutting time per brain. **D–F**. Acquisition time normalised to 12.5 mm of imaged depth plotted against the number of averaged frames, separately for acquisitions of one brain (D), two brains (E), and four brains (F). Points are individual acquisitions, lines join medians of averaging levels represented by at least five acquisitions. Throughout, points are jittered horizontally to reduce over-plotting, and the dashed line marks 24 h. Panels A–C include only data with ≤ 4 frames of averaging for the lower-resolution data and ≤ 2 frames for the higher-resolution data. Rat brains and single-sample acquisitions containing more than one mouse brain’s worth of tissue are excluded (see Methods). Sample sizes are *n*=582 acquisitions (A–C) and *n*=738 acquisitions (D–F). The variability in acquisition times is due to factors such as cutting speed, the amount of agarose being cut, frame averaging, and variability in autoROI behaviour.

**Figure 4–figure supplement 1.**
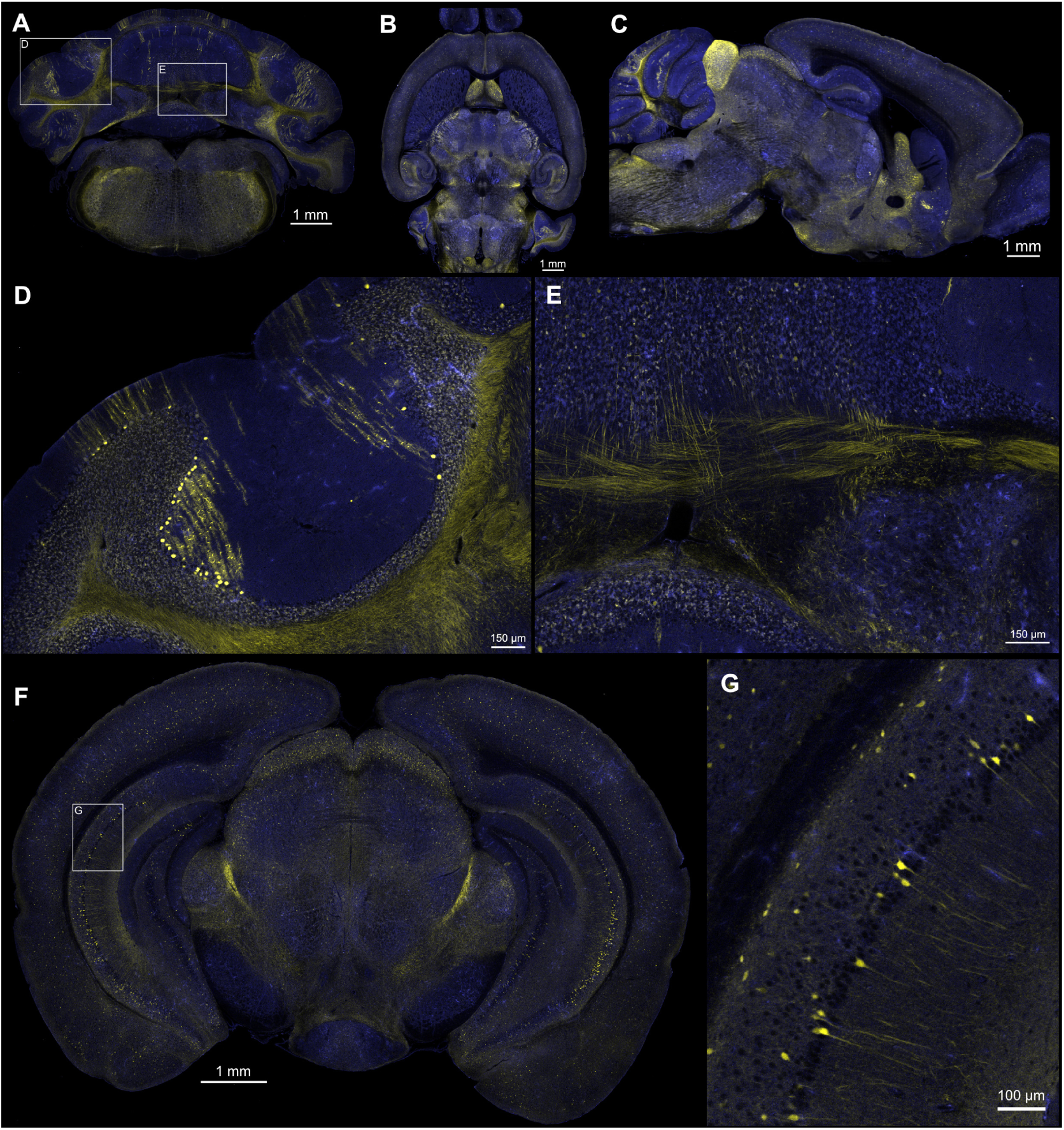
High-resolution mouse brain expressing tdTomato in Sst neurons. Dataset acquired at 920 nm, 155 mW at the sample, 1.2 × 1.2 × 5.0 *µ*m/voxel. Red channel shown in yellow, green channel shown in blue. **A**. Coronal view. Boxed areas marked ‘D’ and ‘E’ indicate the areas of the higher resolution views in those figure panels. **B**. Horizontal view derived from re-slicing a 10 *µ*m isotropic downsampled stack. **C**. Sagittal view derived from re-slicing the same stack. **D&E**. Detailed views from A where individual axons are clearly visible. **F**. A higher resolution coronal section at AP roughly −3.5 mm. **G**. Detail from F showing part of hippocampal area CA1. Data in this figure are available for download at https://doi.org/10.5281/zenodo.22232268.

**Figure 4–figure supplement 2.**
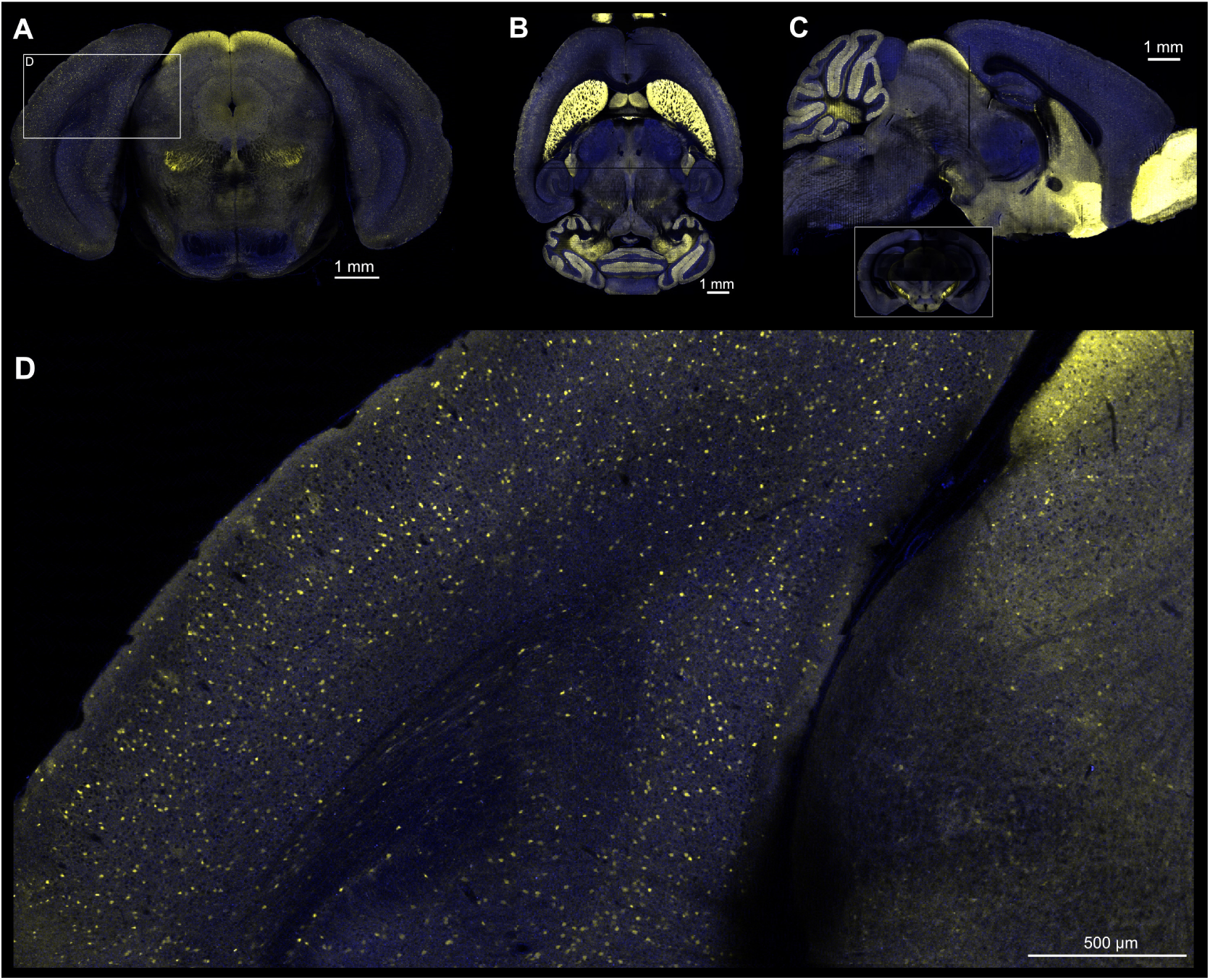
High-resolution mouse brain expressing tdTomato in Gad2 neurons. Dataset acquired at 920 nm, 117 mW at the sample, 2.5 × 2.5 × 5.0 *µ*m/voxel. Red channel shown in yellow, green channel shown in blue. **A**. Coronal view. Boxed area marked ‘D’ indicates the area of the higher resolution view in that figure panel. **B**. Horizontal view derived from re-slicing a 10 *µ*m isotropic downsampled stack. **C**. Sagittal view derived from re-slicing the same stack. Panels B&C show a dark plane just rostral to the hippocampus; this is due to one physical section that was partially obscured by a floating section (see inset figure below C). **D**. Detail from A. Data in this figure are available for download at https://doi.org/10.5281/zenodo.22114627.

**Figure 4–figure supplement 3.**
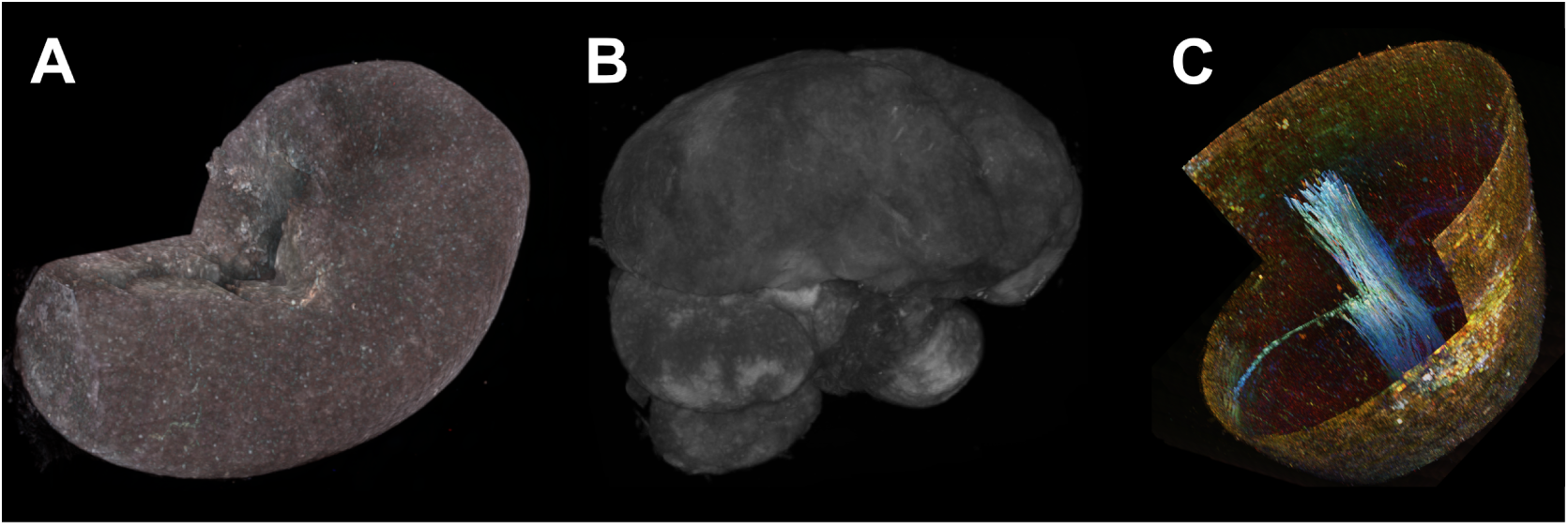
Diversity of imaged samples. STPT can be used to image any sample that cuts well with a vibratome. The main figures show data from rat brains, mouse brains, and mouse spleens. Here we show 3D summary images from three other sample types. **A**. Mouse kidney. **B**. Nightingale brain. **C**. The tip of a fresh carrot.

**Figure 6–figure supplement 1.**
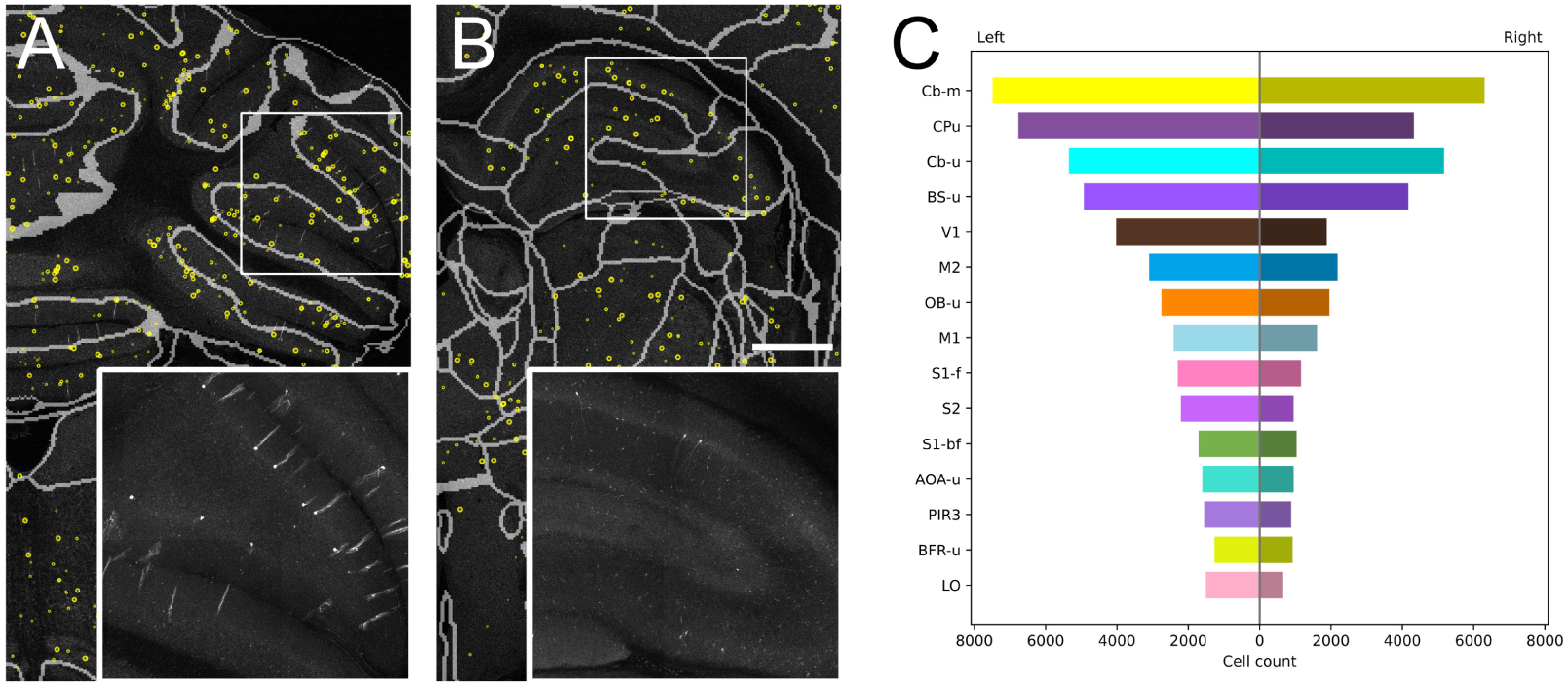
Cell counting and atlas registration in the rat brain. The data shown here are from a rat brain expressing mNeonGreen in a sparse subset of neurons following a systemic AAV injection (data originally gathered for Chen *et al*. 2023). Cellfinder was run on the dataset to locate the labelled neurons brain-wide, and the sample was aligned to a 25 *µ*m resolution rat brain atlas (Plattner *et al*. 2026). **A**. Detail from the cerebellum showing labelled Purkinje cells. Yellow circles mark cells detected by cellfinder; the smaller circles indicate cells whose centres are outside the displayed plane. The inset shows an unannotated maximum intensity projection (10 planes, 50 *µ*m) from the boxed region, centred around the plane of interest. **B**. As A, but for a more rostral section at the hippocampus. The scale bar (1 mm) applies to both A and B, and the boxed regions are 2 × 2 mm. **C**. Number of detected cells in the 15 regions with the most labelled neurons, broken down by hemisphere. *Abbreviations*, in the order plotted: Cb-m, cerebellum, molecular layer; CPu, caudate putamen; Cb-u, cerebellum, unspecified; BS-u, brainstem, unspecified; V1, primary visual area; M2, secondary motor area; OB-u, olfactory bulb, unspecified; M1, primary motor area; S1-f, primary somatosensory area, face representation; S2, secondary somatosensory area; S1-bf, primary somatosensory area, barrel field; AOA-u, anterior olfactory area, unspecified; PIR3, piriform cortex, layer 3; BFR-u, basal forebrain region, unspecified; LO, lateral orbital area.

**Figure 7–figure supplement 1.**
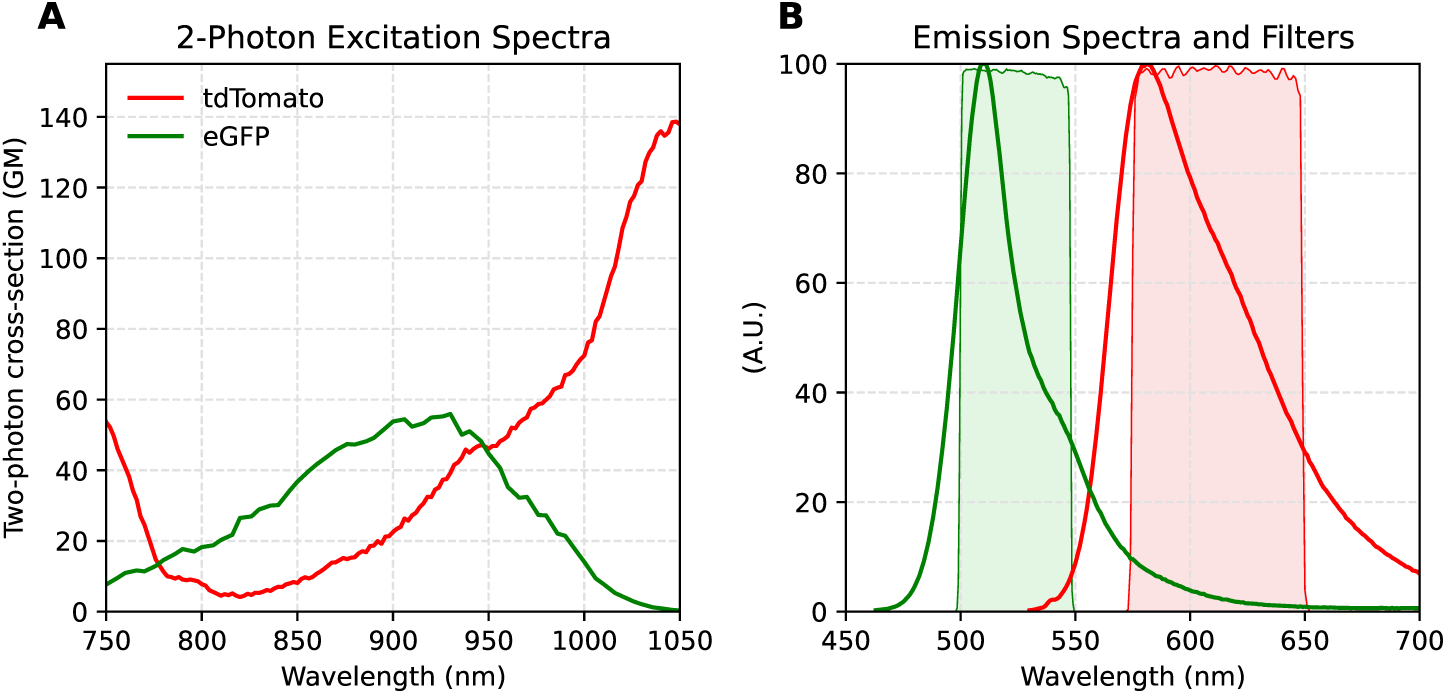
Excitation and emission spectra of tdTomato and eGFP. **A**. Two-photon excitation spectra for tdTomato (red) and eGFP (green). A wavelength of 920 nm provides optimal excitation of eGFP but remains far from the peak excitation of tdTomato. At 950 nm eGFP is still almost at its peak, and tdTomato signal will be substantially stronger. Note that tdTomato is a very bright fluorophore and can work well even in the 920–950 nm wavelength range. **B**. Emission spectra of tdTomato and eGFP along with the emission filter pass-bands typically used in our microscopes. There is significant and unavoidable bleed-through of green into red.

**Figure 8–figure supplement 1.**
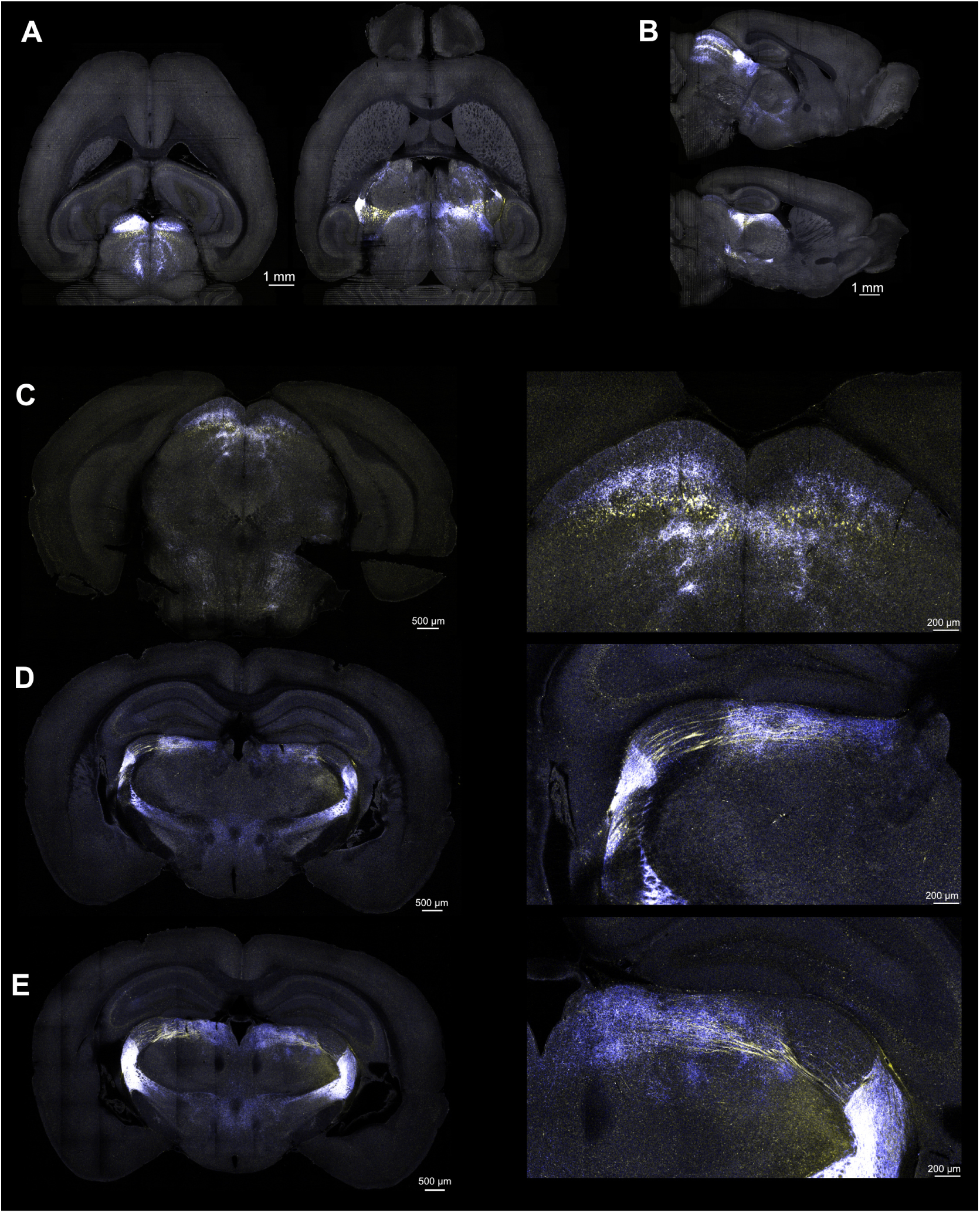
vLGN synaptic mapping supplementary figure. Images show data from a single pilot vgat-Cre mouse which received bilateral AAV[2/1]-hSyn-DIO-tdTomato-Synaptophysin-eGFP injections into the vLGN. Images were filtered with a Gaussian (0.5 pixel SD) to get rid of sparse bright single pixels. Data were acquired with 920 nm excitation, 130 mW at the sample. The red tdTomato signal is rendered in yellow and the green eGFP signal is rendered blue. Areas with co-labelling appear white. **A**. Two different horizontal planes. **B**. Two different sagittal planes. **C–E**. Three different coronal planes (left) with details from a sub-region (right, 2.5 *µ*m/pixel, resampled from 1.25 *µ*m/pixel raw data). Regions with only fibres of passage contrast obviously with regions also containing putative synaptic connections.

## Appendix 1: Costs

Assembling a BrainSaw can be very cost-effective. A typical resonant scanning two-photon microscope used for *in vivo* imaging can become a BrainSaw simply by adding a suitable XYZ stage (Supplementary Table 2) and vibratome. The cost estimate for this upgrade is 30,000–35,000 EUR. The cost of converting a linear (galvo/galvo) scope to resonant scanning is an additional 30,000– 35,000 EUR, plus costs for a fast modulator (such as an electro- or acousto-optic modulator) should that be needed. Exact costs vary by region and with any additional parts required to integrate upgrades into an existing system. The cost of building an entire system from scratch is about 200,000 to 350,000 EUR depending on the desired hardware and how much of the system is DIY-constructed in-house. This cost does not include the laser. Prices are indicative 2026 figures and exclude VAT. Consumable costs for running samples are in the region of 1 EUR per brain.

Ti:sapphire lasers such as Coherent’s Chameleon series or Spectra-Physics’ Mai Tai series are suitable for running a BrainSaw system. BrainSaw systems have much higher operational hours than a conventional *in vivo* two-photon microscope, so the choice of laser manufacturer should be guided as much by the local service environment as by price. The diodes that pump the Ti:sapphire crystal have an operational life of roughly 25,000 hours. Adopters should anticipate costs for laser refurbishment roughly every four to eight years depending on usage.

## Appendix 2: BakingTray as an upgrade path for legacy systems

BakingTray is a viable way to upgrade and extend the life of legacy TissueCyte 1000 systems. The upgrade has been performed on two systems: one at the SWC (‘NeuroVision’, Appendix 2–figure 1) and one at the École Polytechnique (‘ChroMS’, Supplementary Table 3), and both are used heavily. Even without the hardware modifications shown in Appendix 2–figure 1, conversion to BakingTray provides a variety of benefits (depending on the exact version of the microscope being upgraded).

- Acquisition on a single Windows 10/11 PC instead of two Windows XP PCs.
- The option to add a fourth detector channel.
- Easier sample setup with a preview tile scan.
- Automatic sample finding (autoROI).
- The ability to pause acquisitions.
- The ability to easily resume stopped acquisitions.
- Acquisitions will stop if laser mode lock fails or no tissue is detected.
- Laser and PMTs turn off at the end of acquisition.
- Acquisition blocked if there is insufficient disk space.

The system becomes easier to use, which simplifies user training. Furthermore, since BakingTray operates alongside ScanImage, users can do live ‘Focus’ previews whilst panning around the sample, quickly change zoom or image size, etc. This makes various quality control tasks, such as acquiring point-spread functions from fluorescent beads, much easier. Even without the resonant scanner upgrade, overall acquisition speed will improve substantially because of autoROI.

The upgrade re-uses all the original stages and motor controllers. If desired, the vibratome, detectors and pre-amplifiers, and scanners can all be re-used. The PI C-891 linear stage controllers will need a free firmware upgrade and the Z stage stepper controller will need a free USB dongle from the manufacturer in order to communicate with Windows 10/11. The linear scanners from Cambridge Technology used in the TissueCyte 1000 have unique tuning parameters and scan significantly faster than stock Cambridge galvos. Should a scanner fail, replacements with the same specifications cannot be purchased from the original manufacturer (Cambridge Technology). This is not a significant constraint, as the system will perform better overall if converted to resonant scanning, which provides much faster scan speed whilst retaining the ability to average frames when desired. Bidirectional scanning artefacts are also easier to manage with a resonant scanner. With bright labelling, resonant scanning is not a compromise in image quality: the high-resolution single-axon tracing work at Janelia was performed on a resonant-scanning STPT rig (Economo *et al*. 2016; Winnubst *et al*. 2019), as was the work from Liu *et al*. 2022 on BrainSaw.

The upgrade should be performed carefully: some hardware changes are tricky to make (in particular the coarse Z motor upgrade) and these changes are hard to revert once made. Once the upgrade is complete, the end-user becomes responsible for their own support. The exact upgrade costs depend upon how many changes are desired. Minimum requirements are a new acquisition PC, ScanImage licence, a new DAQ, and an EOM to rapidly modulate laser power (the TissueCyte’s LCVR is not supported by ScanImage). The cost of the above is roughly 50,000 EUR assuming a high-end DAQ was purchased. It is possible to re-package the TissueCyte’s sprawling open collection path into a more compact closed design (Appendix 2–figure 1A) whilst either retaining the original PMTs and pre-amplifiers or upgrading these also. It is also a good idea to add bandpass filters, if these were missing originally, since doing so should reduce spectral cross-talk. The collection path can be upgraded in this way for about 5,000 EUR. The addition of motorised coarse Z is very helpful and can be done for under 5,000 EUR. A resonant scanner upgrade is optional and only adds the cost of the scanner and associated opto-mechanics (around 5,000 EUR), assuming a high-end DAQ, such as a ScanImage vDAQ, was purchased for the base upgrade. A fully specified upgrade is therefore in the region of 65,000 EUR.

**Appendix 2–figure 1.**
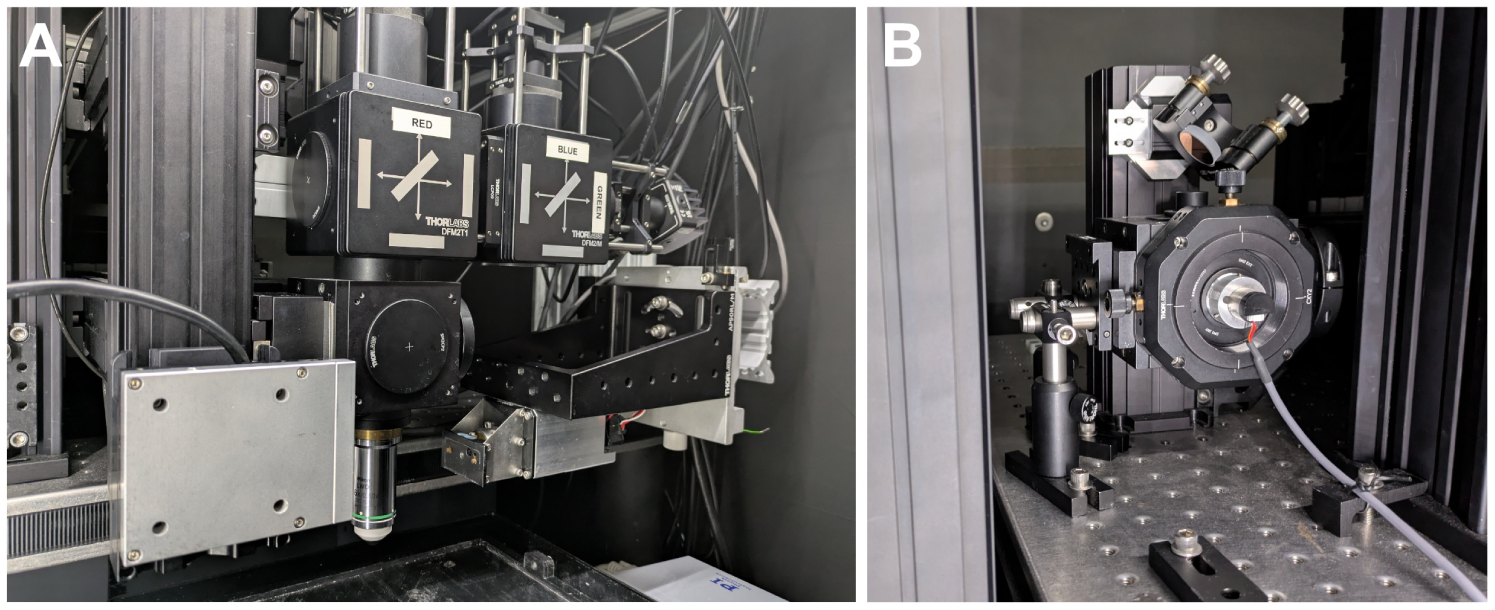
‘NeuroVision’ STPT system at the SWC. This is a modified version of the legacy TissueCyte 1000 (TissueVision). **A**. Front view showing objective with coarse Z motor, enclosed detection path based around SiPMs, and Leica vibratome. **B**. NeuroVision has a resonant scanner enclosed in a sealed Thorlabs optical cube that clamps directly onto the original beam expander. The resonant scanner conversion is optional but recommended.

## Appendix 3: Acquisition safeguards and making settings changes mid-run

**Appendix 3–table 1.** BakingTray safeguards. . This table itemises the safeguards BakingTray applies before and during acquisitions to help avoid errors. In addition to these, BakingTray has other useful features. Interrupted acquisitions can be easily resumed, and acquisitions can be paused at any time for sample inspection. Various key settings, such as frame averaging, laser power, and PMT gain, can be adjusted mid-run without stopping the acquisition.

| Pre-acquisition safeguards | Safeguards during acquisition |
| --- | --- |
| • Stage limits prevent the objective, blade, or blade-holder crashing into the water bath.<br>• Acquisition will not begin if there is insufficient disk space.<br>• Acquisition will not begin if the PMT channels enabled in ScanImage do not match those selected to be saved.<br>• Users are blocked from focusing by moving the sample stage in Z once the first trimming slice has been taken. | • Acquisition stops if no tissue is detected.<br>• Acquisition stops if the laser loses mode lock.<br>• A Slack message is sent if acquisition stops unexpectedly.<br>• The last completed section is stitched, down-sampled, and streamed to a web page for remote monitoring.<br>• Laser switched off at the end of acquisition to preserve diode lifetime.<br>• PMT gains set to zero at the end of acquisition to reduce chance of accidental damage. |

### Changing settings during an acquisition

It is sometimes necessary to tweak acquisition settings after the acquisition has begun. Some settings can be changed freely during acquisition; others require the acquisition to be stopped and restarted; and some cannot be changed at all, either because the resulting data could not be merged with what has already been acquired or because stitching would fail (Appendix 3–table 2).

The most common adjustment is **objective height**, which is used to correct any drift in the distance between the objective and the block face (see Discussion Section 3.5). **Laser power** can also be adjusted at any time, but a large change introduces a step in image brightness at the section where it is made, complicating comparisons of intensity through the volume. Changing the **laser wavelength** is occasionally necessary in older tunable lasers that are in need of service: lasers sometimes drop in and out of mode lock at a particular wavelength, and moving 5 to 10 nm away is often enough to restore stable operation (Appendix 3–figure 1). Two-photon excitation spectra are broad, so a shift of this size has little practical effect on the signal and is greatly preferable to generating noisy data or having the acquisition stop prematurely late at night. **Frame averaging** can be changed whilst the system is cutting but not whilst it is imaging.

BakingTray has the ability to resume an acquisition. This is useful if there was a system crash or if some settings, such as **cutting speed** or the **number of physical sections**, need changing. The acquisition is stopped via the Acquisition GUI. The user selects ‘Resume acquisition’ under the File menu and loads the ‘recipe’ (settings file) associated with the current acquisition. A window appears asking how the user wishes to proceed (e.g. re-image the last section or slice and carry on). The required settings can now be changed. The user presses ‘Bake’ and the acquisition resumes.

Settings related to the image **resolution**, **number of optical planes per section**, **detector channels**, or **section thickness** require a new acquisition in a new directory.

**Appendix 3–figure 1.**
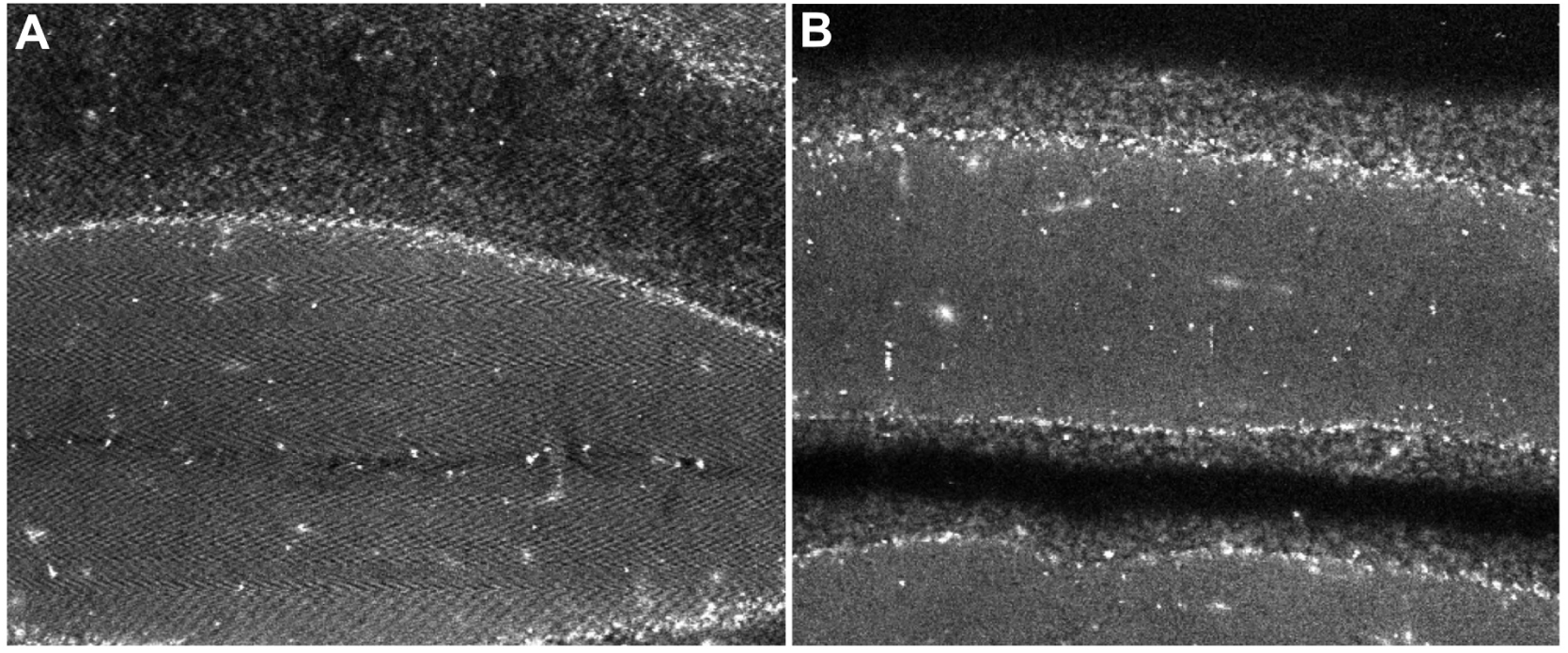
Lasers dropping in and out of mode lock produce imaging artefacts. **A**. Detail from a single frame in ScanImage showing the characteristic herringbone pattern of a laser dropping rapidly in and out of mode lock. The image comes from an 8 kHz resonant scanning microscope and shows autofluorescence in mouse cerebellum. **B**. Similar field of view to that in A after the wavelength is jogged by 10 nm: the laser is now mode locking perfectly.

**Appendix 3–table 2.** Changing acquisition settings once a run has started. . Settings that can be **changed freely** may be altered at any point without interrupting the run. Settings that **require a restart** need the acquisition to be stopped, the setting changed, and the acquisition resumed. Settings that are **fixed for the acquisition** cannot meaningfully be altered, as the resulting data could not be merged into a single dataset afterwards.

| Changed freely | Requires restart | Fixed for acquisition |
| --- | --- | --- |
| Objective height | Cutting speed | X/Y resolution |
| Laser power | Number of physical sections | Optical planes per section |
| Laser wavelength |  | Detector channels |
| PMT gain |  | Section thickness |
| Frame averaging<sup>†</sup> |  |  |

<sup>†</sup> Frame averaging can be altered without stopping the acquisition, but only during the slicing phase of the cycle; it cannot be changed while ScanImage is acquiring data.

## Appendix 4: System alignment

### Stage and imaging plane alignment

The plane of motion of the X/Y stages must be aligned with the imaging plane. Misalignment causes overlapping regions of adjacent tiles to sample different z-planes, which manifests as a discontinuity along tile borders that cannot be corrected during stitching. Because optical planes are typically spaced 5–20 *µ*m apart – insufficient to allow affine correction along the axial dimension – physical alignment is necessary.

The X/Y/Z stage assembly is mounted on three Thorlabs adjustable height pedestal posts (BLP01), which provide fine tilt control. Alignment is assessed by imaging a slide containing a thin fluorescent layer (FluoroCal, Valley Scientific). A cheaper alternative is cover-slipping a 1 *µ*L drop of secondary antibody conjugated to a fluorescent dye. With correct alignment and no field curvature, the fluorescent layer appears uniform across the field of view; field curvature produces a symmetric ring (‘donut’) pattern (Appendix 4–figure 1B). When unaligned, the FOV will show a curved edge (Appendix 4–figure 1A) or even a bright band (for more uses of these thin slide targets, see Campbell, Eifert, and Turner 2014 Fig. 7). The pedestal post heights are adjusted until the image is as uniform and symmetric as possible. Before finally securing the posts, the stage motion axes are checked for parallelism with the scanning axes by translating a small feature across the field of view and confirming it remains on the same scan line. Any error is corrected by rotating the stages about the axis of the objective. The tilt alignment can be sensitive and may change slightly when posts are clamped to the optical table. Note that any field curvature remains in the data and has no practical consequence, as the curvature is identical in every tile and introduces no discontinuity at tile boundaries after the above alignment procedure.

**Appendix 4–figure 1.**
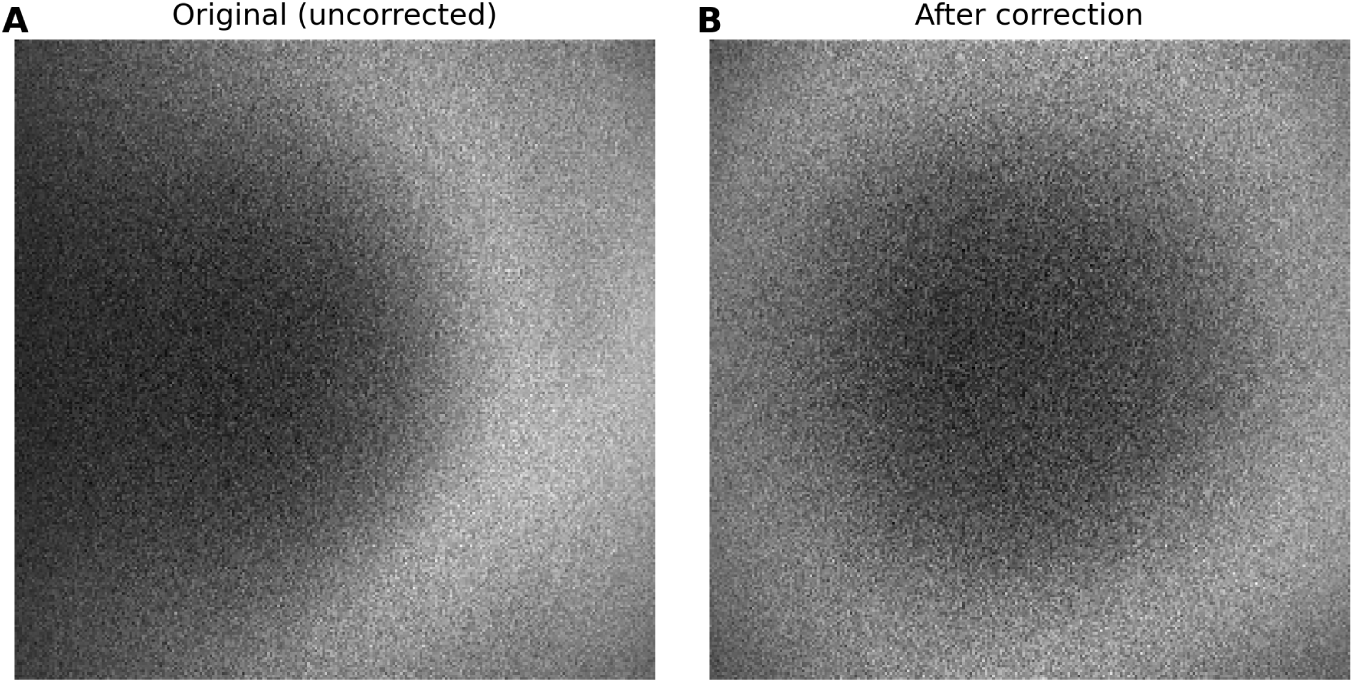
Example stage tilt calibration images. **A**. Image of a thin fluorescent layer taken on a microscope where the stage and imaging planes are not well aligned. The image is taken at ScanImage’s zoom=1 setting and is 1.1 × 1.1 mm. **B**. The same field of view after adjustment of the stage angle such that it is parallel with the imaging plane. The ring-like shape is due to field curvature. Objectives corrected for field curvature will not exhibit this pattern.

### Vibratome and stage alignment

The X stage translates the sample into the vibratome blade during cutting. We mount the vibratome to a Thorlabs AP90RL right-angle bracket (Appendix 4–figure 2). The vibratome blade edge must be parallel to the Y stage motion axis. The Y stage is the one that translates the sample toward and away from a user standing in front of the microscope. If the blade and Y axis are not parallel, the sliced block surface will vary in height along the Y direction with respect to the imaging plane, producing serious imaging artefacts (Appendix 4–figure 2A). Alignment is assessed using a fluorescent agarose block made by adding a small volume of fluorescent microsphere suspension, obtained cheaply from a highlighter pen, to the agarose during preparation. After slicing, the relative surface height of the block is measured at two distant points along the Y axis using the microscope coarse focus control. The vibratome angle is adjusted by pivoting it around the central screw of the AP90RL bracket. Precision adjustment is enabled by a Thorlabs KL02 adjustable kinematic positioner, which contacts the right-angle mount at one end (Appendix 4–figure 2B, arrowed). One turn of the KL02 adjuster will change the relative heights of two points on the agarose 20 mm apart in the Y axis by about 100 *µ*m. A residual height variation of 10–20 *µ*m over 20 mm of the agarose block is achievable and quite acceptable. This calibration needs to be performed when the system is first built, but some batches of blades are variable, so recalibration is occasionally needed. If a height difference is detected, the first step is to replace the blade; recalibration of the vibratome angle is only necessary if the problem persists. End-users typically find this calibration challenging, so they are usually trained to notice the issue and change the blade, rather than alter the vibratome angle.

We provide a CAD model illustrating how to modify the bracket as shown in Appendix 4–figure 2B to reduce play at the pivot point and require only a single screw (SWC-Advanced-Microscopy/BrainSaw on GitHub). The adjustment process remains feasible even with an unmodified bracket.

**Appendix 4–figure 2.**
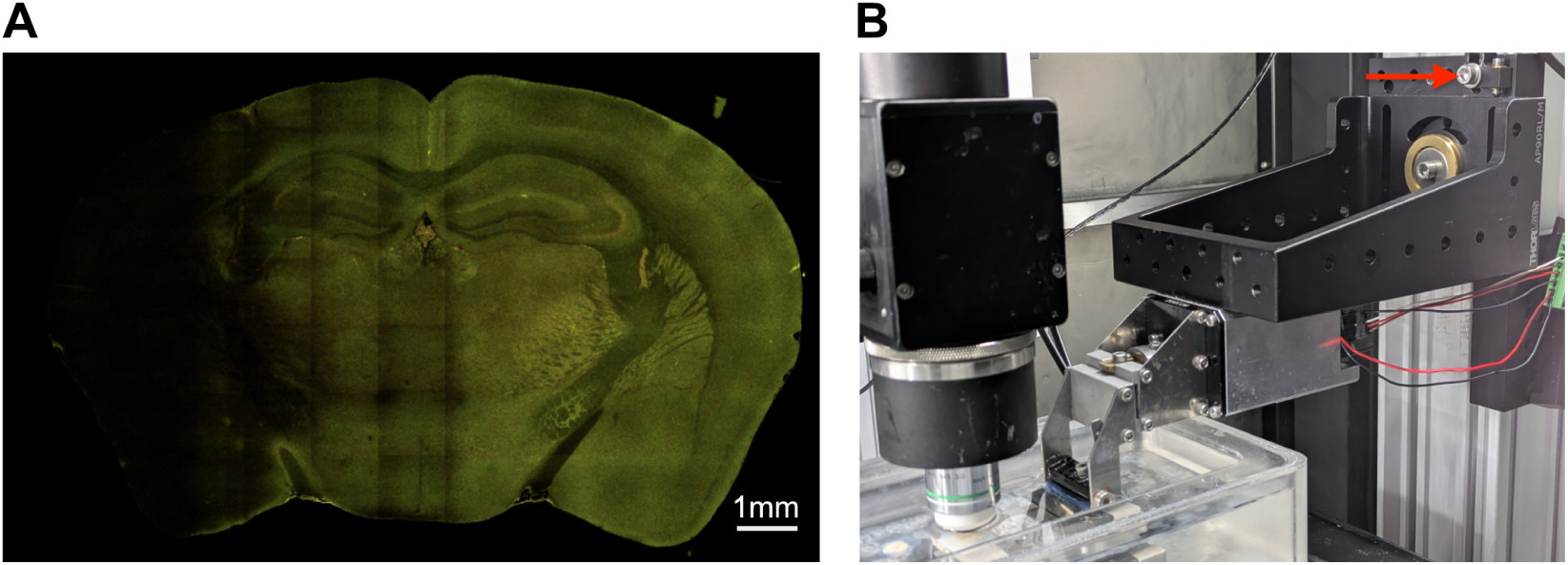
Vibratome alignment. **A**. An imaged plane showing evidence of severe misalignment between the blade and the Y stage axis. In the black region to the left, the focal plane is above the sample surface, meaning the blade is slicing deeper on that side. **B**. The vibratome is mounted on a Thorlabs AP90RL right-angle bracket. The right-angle bracket in this image has been modified to accept a brass bushing as a single clamping point (details at BrainSaw_CAD). This allows the unit to be easily clamped and unclamped with a single screw. The bushing also eliminates play, which makes adjustment more accurate. Unmodified brackets can also be used, but they require two screws for security and are slightly harder to work with. Note the critical Thorlabs KL02 adjustable kinematic positioner arrowed in the top right of the picture.

## Appendix 5: Vibratome failure modes and ‘Auto-Trim’

BakingTray’s acquisition procedure assumes slice thickness is consistent over sections, and therefore the distance between the objective and the sample surface does not vary during the acquisition. Because agarose blocks are somewhat elastic, they can introduce feedback into the cutting cycle that breaks the assumption of a fixed block surface height (objective to sample distance). Specifically, it is possible for a vibratome to alternately cut sections that are too thick and too thin. Alternating thick/thin sections may arise during setup and trimming; we have not observed it developing spontaneously once a run is underway. Once this cycle begins it rarely breaks on its own, and the vibratome continues cutting thin and thick sections alternately, manifesting as the block surface rising and falling by as much as 30–40 *µ*m from cut to cut with respect to the objective. Acquisitions should not be started if this is the case.

To minimise the chance of inconsistent cutting developing, once we have completed initial trimming of the block (which we typically do at around 200 *µ*m), we reduce cut thickness gradually to our target thickness (typically 40–50 *µ*m). This process is automated by BakingTray via its ‘Auto-Trim’ button. The Auto-Trim cycle finishes with three sections cut at the target thickness. Using the imaging system, the user measures the sample surface height directly across successive slices to confirm that it does not drift.

On occasion, surface drift persists despite the above measures. This can almost always be resolved by cutting a substantially thicker section and returning to the desired section thickness gradually; for instance, moving from the current 40 *µ*m slices to 80 *µ*m and stepping back down to 40 *µ*m in 10 *µ*m increments.

## Appendix 6: Whole-brain antibody labelling

This antibody labelling protocol takes two weeks and delipidates brains with tetrahydrofuran (THF) according to the EZ Clear method (Hsu *et al*. 2022) and labels using the permeabilising buffers from iDISCO (Renier *et al*. 2014). All steps below are conducted in conical-bottomed 15 mL polypropylene sample tubes, apart from the THF step, which requires glass tubes.

### Buffer recipes

- PTx.2 (1L): 100 mL 10X PBS, 900 mL dH_2_O, 2 mL Triton X-100
- PTwH (1L): 100 mL 10X PBS, 900 mL dH_2_O, 2 mL Tween-20, 1 mL of 10 mg/mL heparin stock
- Permeabilisation solution (500 mL): 400 mL PTx.2, 11.5 g glycine, 100 mL DMSO
- Blocking solution (50 mL): 42 mL PTx.2, 3 mL donkey serum, 5 mL DMSO

### Day 1: Delipidation

1. Prepare a total volume of 10 mL of THF diluted 50:50 in water in a glass vial with polypropylene screw cap. THF may melt other plastics
2. Immerse the brain in the diluted THF at room temperature for two days on a shaker, refreshing the solution after 24 hours
3. Wash: 3 × 1 hour Milli-Q water
4. Wash once with PBS

### Day 3: Permeabilisation

Although samples were delipidated with THF, this step still enhances antibody penetration through remaining membranes

- PTx.2, 1 hour, room temperature, 12 mL, shaking
- PTx.2, 1 hour, room temperature, 12 mL, shaking
- Permeabilisation solution, 37^◦^C, 7.5 mL, shaking for 3 nights

### Day 6: Block

Blocking solution 4 mL, 37^◦^C, 2 days, shaking

### Day 8: Primary antibody on

Add TH antibodies diluted 1:50 in 1.5 mL blocking solution, 37^◦^C, 7 days, shaking

### Day 15: Washes

All washes have a volume of 12 mL and are done at room temperature on a shaker.

- PTwH, 10 minutes
- PTwH, 20 minutes
- PTwH, 30 minutes
- PTwH, 1 hour
- PTwH, 2 hours
- PTwH, overnight
- 50 mM PB, 3 × 1 hour to remove the wash buffer and prepare brain for agarose embedding
- Image on BrainSaw with 800 nm excitation, 125 mW at the sample, collecting the far-red and green channels

## Appendix 7: Half-brain Congo red labelling

The protocol labels plaques in Alzheimer’s mouse brains using Congo red. Brains are first delipidated with CUBIC-L (Tainaka *et al*. 2018) before being further permeabilised with ScaleCUBIC-1a (Susaki *et al*. 2014) and labelled with Congo red. Brains are strengthened by embedding them in gelatin, which is then cross-linked with PFA, in order to improve sectioning quality for STPT imaging. This protocol was tested in half-brains.

### Solutions

- CUBIC-L: TCI Chemicals, Product No. T3740
- ScaleCUBIC-1a: 10% wt Triton X-100, 5% wt Quadrol, 10% wt urea, 500 mM NaCl, dissolved in dH_2_O. e.g. 70 mL dH_2_O, 10 g Triton X-100, 5 g Quadrol, 10 g urea, 2.922 g NaCl. Top up to 100 mL with dH_2_O

### Delipidation

1. Dilute 5 mL CUBIC-L 50:50 in water for 10 mL total
2. Immerse sample at room temperature on rollers for six hours or overnight
3. CUBIC-L, 7 mL, 37^◦^C, shaker, 3 days
4. Refresh CUBIC-L and shake at 37^◦^C for a further two days
5. PBS wash 3 × 2 hours, 37^◦^C on a shaker

### Labelling

- Congo red, 5 *µ*M (stock is 10 mM) diluted in ScaleCUBIC-1a, 37^◦^C, shaker, 4 days
- Wash 3 × 2 hours PBS 0.1% Triton X-100, at room temperature. One overnight wash in the fridge

### Tissue strengthening

CUBIC-L makes brains soft, so they cut poorly; to partially counteract this, we immersed the brains in gelatin and then cross-linked the gelatin with PFA. Shortening the delipidation step may leave the tissue firmer without compromising Congo red penetration; we have not tested this.

- Make 5% gelatin solution in dH_2_O
- Stir for around 10 minutes at 42^◦^C until the gelatin has melted
- Incubate brain in the gelatin at 37^◦^C for 4 hours
- Remove the brain from the gelatin and air dry, blowing on it and flipping it around so the gelatin solidifies
- Fix the brain in 4% PFA overnight in the fridge to cross-link the gelatin
- Wash in 50 mM PB before imaging on BrainSaw at 820 nm excitation, 80 mW at the sample, collecting blue and red channels

